# Mutations in filamentous bacteriophages spark eco-evolutionary feedbacks in *Pseudomonas aeruginosa*

**DOI:** 10.64898/2026.01.21.699487

**Authors:** Noah S. B. Houpt, Catherine A. Hernandez, Paul E. Turner

## Abstract

Microbial populations strongly shape their environment, which can re-route adaptation toward organism-generated fitness optima. However, the conditions that promote these eco-evolutionary feedbacks are unclear. Here, we used experimental evolution to test whether high population density, by strengthening niche construction, drives eco-evolutionary feedbacks in the bacterial pathogen *Pseudomonas aeruginosa* MPAO1. We then tested for adaptation to organism-modified environments by measuring the relative performance of ancestral and endpoint populations in filtrate generated by each evolutionary line sampled across generations. Contrary to expectations, we found that endpoint populations had higher performance than the ancestral strain in filtrate across nearly all evolutionary lines regardless of population density. This was caused by the emergence of hyperactive filamentous bacterio(phage) mutants during experimental passaging that inhibited the ancestral strain but not endpoint populations in modified media. Hyperactive phages emerged from one of two avirulent prophages in MPAO1’s genome (Pf4 or Pf6). Hyperactive phages drove the evolution of phage resistance in bacterial populations via mutations in the type IV pilus (TIVP), the phage’s binding receptor. In a follow-up experiment, we showed that these TIVP mutations pleiotropically reduced motility and decreased susceptibility to a TIVP-targeting virulent phage, both of which are important traits for *P. aeruginosa* infection and treatment. Overall, this work suggests that filamentous phage evolution can act as a driver of eco-evolutionary feedbacks in bacterial populations, causing phenotypic and genetic changes that would not be anticipated from adaptation to the extrinsic environment alone.

## Introduction

Classical conceptions of adaptive evolution ignore the evident: organisms strongly shape their environment and, thus, the selective pressures they face [1–4]. This process, often referred to as niche construction, links evolutionary and environmental change, rerouting evolution towards fitness peaks generated by the organism itself, rather than the extrinsic environment (ie. eco-evolutionary feedbacks) [3, 5–8]. Consequently, in some cases, adaptive evolution may only make sense in the light of niche construction.

Niche construction may be particularly important to making sense of adaptation in microbial systems. Microbes are prolific niche constructors, shaping their environments through resource consumption, pH modification, and the release of metabolites, among other mechanisms [9–12]. These mechanisms can, in turn, shape the ecology of microbial communities, tipping the outcome of competition towards species that thrive in modified environments [12–17]. As an apparent consequence, microbes are replete with traits that are beneficial specifically in organism-constructed environments and many species depend on niche construction for survival [11, 18–20]. Furthermore, microbial evolution experiments frequently result in populations adapting to organism-generated environmental change rather than the extrinsic environment [7, 13, 15, 16, 21–23]. Despite the importance of niche construction in microbial systems, when and where we should expect niche construction to be most pronounced in driving adaptation remains understudied [8]. Working this out requires generating and testing hypotheses on the conditions that promote adaptation in response to organism-generated environmental change.

One factor long thought to modulate the importance of niche construction is population density. High population density magnifies the impact of populations on their environment, strengthening feedbacks between the phenotypes of populations and the selective pressures they face [4, 24–26]. For example, prior work found that soil bacteria modify the pH of growth media in a density-dependent manner: pH change was minimal at low population density whereas high density populations changed the pH to such an extent that it could drive them extinct [25]. The density-dependence of niche construction implies that sustained high population density should favour adaptation to organism-generated environments over the extrinsic environment. However, no study has tested how population density affects the balance between these modes of adaptation.

Here, we explored how population density impacts adaptation to organism-generated environments in the bacterium *Pseudomonas aeruginosa* (*Pa*) over a 1,000-generation evolution experiment. *Pa* is a model organism and opportunistic pathogen that engages in several well-studied forms of niche construction, including the production of quorum sensing autoinducers, biofilm components, iron-scavenging siderophores, and pyocins [9, 27–29]. Given that these and other mechanisms of niche construction are more pronounced at high population density, we predicted that evolution under high population density would more frequently drive adaptation to organism-constructed environments. Contrary to these expectations, we found that endpoint populations adapted to organism-constructed environments in nearly all experimental lines regardless of population density. This was driven by the emergence of hyperactive filamentous bacteriophage (phage) mutants during experimental passaging that evolved from one of two prophages (Pf phages; Pf4 and Pf6) present in the ancestral strain’s genome. Although these prophages are normally avirulent, mutations that disrupt their respective repressor genes lead to high rates of phage replication (ie. hyperactivity), decreased host growth, and in some cases, host lysis [30–35]. The emergence of hyperactive phages sparked eco-evolutionary feedbacks, driving the evolution of phage resistance across populations. Collectively, our work suggests that organism-driven environmental change, in this case via mutations in an integrated virus, can strongly shape adaptive evolution across population densities.

## Materials and Methods

Please see the Supplementary Methods file for comprehensive description of the methods used in this study.

### Evolution experiment

16 populations of *Pa* MPAO1 were experimentally evolved at either high (HD) or low (LD) population density for 1,000 generations (Figure 1A). All populations were grown in M9 media (22 mM KH_2_PO_4_, 48 mM Na_2_HPO_4_•H_2_O, 8.6 mM NaCl, 19 mM NH_4_Cl, 1 mM MgSO_4_, 100 µM CaCl_2_, 18 µM FeSO_4_) supplemented with 37 mM sodium succinate as the sole carbon source. Populations were grown in well-mixed (200 rpm) 1.5 mL cultures in 24 well tissue culture plates (CellTreat, #229524) incubated at 30 °C for 23 h each day before being transferred to fresh media.

**Figure 1:**
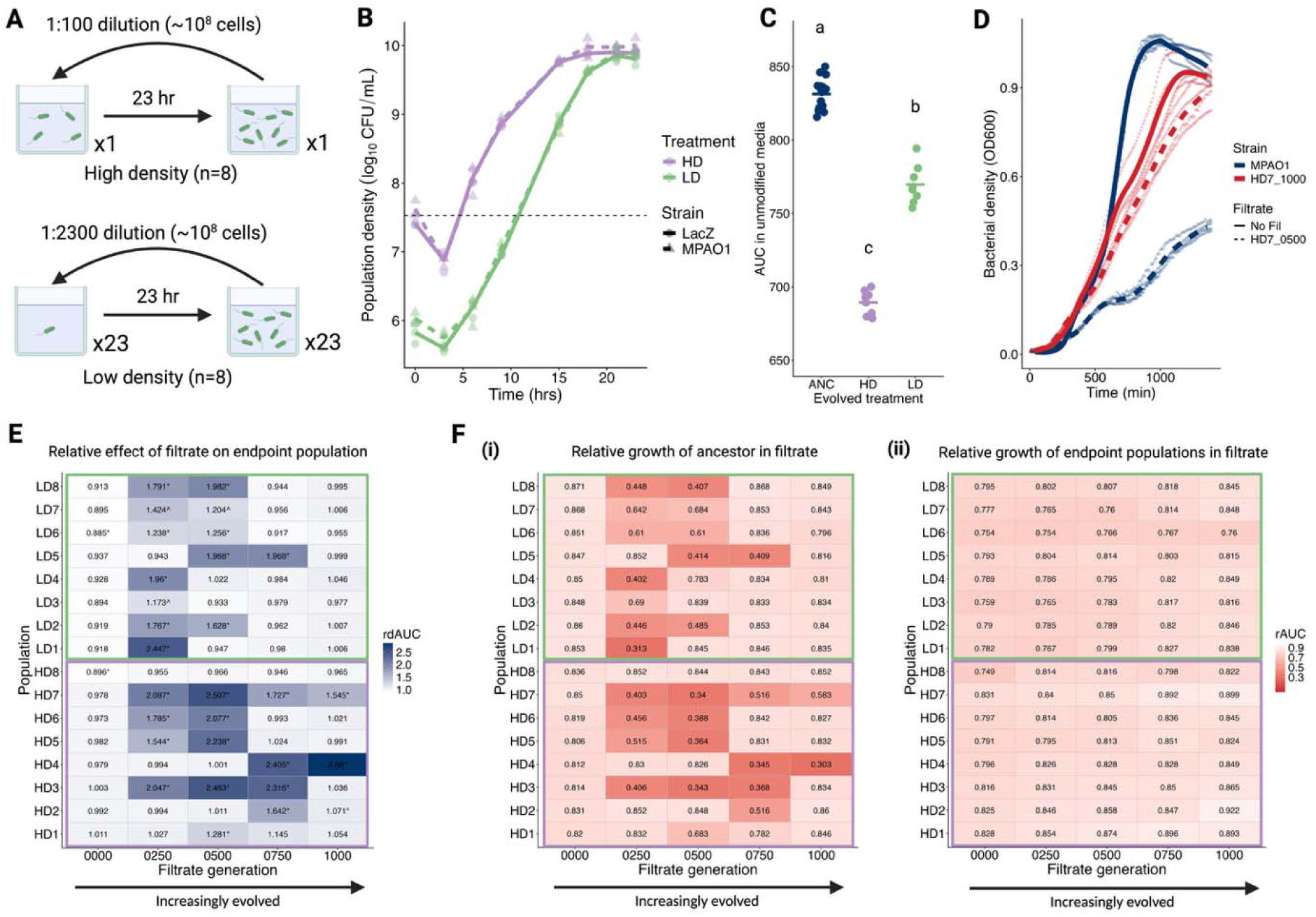
Passaging at both high and low population density leads to the evolution of inhibitory filtrate. **A**: Passaging design of evolution experiment. **B**: Population density of the unmarked (MPAO1) and genetically marked (LacZ) ancestral strain when growth in the high density (HD) and low density (LD) treatment for one growth cycle. Colored lines trace the average density of three replicate populations for each strain and treatment. Horizontal dashed line shows the average starting density for the HD treatment. **C**: Area-under-the-curve (AUC) of ancestral and endpoint populations in unmodified media. Each point represents the mean AUC across four replicate curves for each endpoint population (n = 8) or paired replicate curves of the ancestral strain (n = 16). Horizontal bars represent mean values for each group and letters denote statistically distinct groups. **D**: Ancestral (MPAO1) and endpoint (HD7_1000) population of line HD7 grown in the presence or absence of filtrate generated by HD7’s evolutionary midpoint (HD7_0500). Lines represent moving averages (bin width = 70 min) of 4 replicate growth curves, which are shown as faded circles (no filtrate) or triangles (HD7_0500 filtrate). **E**: Mean rdAUC for each endpoint population in filtrate sampled across generations (n = 4). * indicates a statistically significant advantage of either ancestral or endpoint populations in a given filtrate (*P* < 0.05 after correcting for multiple comparisons). ^ indicates filtrates where the endpoint populations had a relative growth advantage in some replicates but not others. **F**: Mean rAUC values for i) ancestral and ii) endpoint populations in filtrate sampled across generations (n = 4). Created in BioRender. Turner, P. (2026) https://BioRender.com/94m4mlj. Alt text: Multipaneled figure visualizing the relative performance of ancestral and endpoint bacterial populations grown in the presence and absence of filtrate in microplate growth curves.

HD lines were grown in a single well of a 24 well plate and diluted 1:100 each day into fresh media (6.64 generations/day). Conversely, LD lines were split across 23 wells which were combined and diluted 1:2300 into 23 new wells filled with fresh media daily (11.17 generations/day). This resulted in an equivalent population bottleneck across treatments (∼10^8^ cells), but a 23-fold lower starting density in LD lines each day. Consequently, LD populations experienced lower average population density during each growth cycle (Supplementary Figure 1). Differences in starting density of similar magnitude have been shown to alter the relative performance of bacterial strains varying in their performance in organism-constructed environments [13, 25, 36]. End-of-passage population density was measured every 2-7 passages by diluting and plating each population on agar plates (M9 + succinate media, 1.5% agar). Due to the different number of bacterial generations experienced each day across treatments, HD and LD lines were passaged for a different number of days (HD: 151 days, LD: 90 days).

### Microplate growth curves in filtrate

Filtrate was generated by first reviving populations from frozen stocks and allowing them to grow for two 23-h growth cycles as described above. We then centrifuged 1 mL of the second day’s culture in a 1.5 mL Eppendorf tube for 10 min at 10,000 g and filtered the supernatant using a syringe-tip 0.22 µm PDVF filter (Millex, SLGVR33R5). The leftover pellets were then resuspended in 1 mL of M9 salt solution and were centrifuged and resuspended once more to minimize carryover of spent media.

Growth curves were run in flat bottom 96-well plates (Falcon, #351172) in either unmodified media (190 µL 1x media, 10 µL OD_600_ = 0.5 bacterial culture) or in media supplemented with filtrate (126.7 µL media with 1.5x concentrated succinate, 63.3 µL filtrate, 10 µL OD_600_ = 0.5 bacterial culture). Curves were run for 23 h at 30 °C with constant double-orbital shaking and optical density (OD_600_) was measured every 10 min in a BioTek Epoch 2 plate reader.

The relative performance of ancestral and endpoint bacteria in filtrate was quantified using two metrics derived from their area-under-the-curve (AUC) across growth conditions. The AUC from growth curves was extracted using the R package *gcplyr* [37]. The impact of filtrate on a population was quantified using the quotient of its AUC in the presence vs. absence of filtrate, which we called the relative AUC, or rAUC (Supplementary Methods equation 1). Additionally, the extent to which endpoint populations were adapted to filtrate was quantified by calculating the quotient of rAUC values from endpoint populations and ancestral populations in the same filtrate, which we called the relative difference in AUC, or rdAUC (Supplementary Methods equation 2). rdAUC values exceeding 1 indicate that the endpoint population performed comparatively well in filtrate compared to unmodified media, relative to its ancestor.

### Phage quantification

The densities of hyperactive phage in filtrate were quantified using a conventional plaque assay on lawns of MPAO1 on lysogeny broth (LB) agar plates (15 g/L agar) and LB top agar (4 g/L agar).

### Phage removal/addition experiment

Hyperactive phage-containing filtrates from a subset of populations and evolutionary timepoints were generated as described above. In the phage removal experiment, phages were removed from filtrates using 10 kDa cellulose centrifugal filters (centrifuged at 3,220 rpm for 10 min; Amicon Ultra 4, Millipore, UFC9010). Centrifugal filtration successfully removed phages in all but one replicate, which was consequently not considered in our analysis (Supplementary Figure 2). In this experiment, MPAO1 was grown in microplate growth curves as above, but in the presence of unmodified ancestral or hyperactive phage-containing filtrate, or the same filtrates after centrifugal filtration.

In the phage addition experiment, three phage isolates were first collected from each of the filtrates used in the phage removal experiment. Then, MPAO1 was grown in microplate growth curves in the presence of ancestral filtrate, evolved, phage-containing filtrate, or ancestral filtrate with a mix of the three phages isolated from each filtrate added in at the same total phage density as was present in the evolved filtrate.

### Phage isolate sequencing and genomic analysis

Phenol chloroform DNA extractions were performed on our phage isolate collection using established protocols (https://barricklab.org/twiki/bin/view/Lab/ProtocolsPhageGenomicDNA). Briefly, we concentrated phage stocks by mixing them (4:1) with 5x phage precipitate solution (20% w/v PEG 8000, 2.5 M NaCl), incubating them overnight at 4 °C, and centrifuging them for 30 min at 3,220 rpm to pellet the precipitated phage. Phage were then resuspended in phage resuspension buffer (1M NaCl, 10 mM Tris•HCl (pH 7.6), 0.1 mM EDTA) before DNAse and RNAse treatment. Then, 500 µL of phenol:chloroform:isoamyl alcohol (25:24:1) was added to each sample, which was then mixed by inversion, and centrifuged for 5 min at 16,000 g. The aqueous phase was then transferred to fresh tubes and incubated overnight in 70% ethanol at −20 °C. Sampled were then centrifuged for 15 mins at 14,000 g and the supernatant was removed. DNA pellets were dislodged in 500 µL of pre-chilled 70% ethanol and centrifuged for 5 minutes at 14,000 g. Subsequently, the supernatant was removed and, after allowing the samples to air dry, pellets were resuspended in 200 µL of 10 mM Tris•HCl (pH 7.6). Extracted DNA was then purified using a DNA purification kit (Zymo Research DNA Clean & Concentrator Kit, D4013) and ssDNA concentrations were measured via spectroscopy (Nanopore One, Thermo Scientific). Extracted DNA was sent to Plasmidsaurus for Oxford Nanopore sequencing using their small linear dsDNA sequencing package. Raw sequencing reads are available on the NCBI Sequence Read Archive (PRJNA1420075).

Raw reads were filtered for quality using *fastplong* (mean base quality > Q20) [38] and then used to call variants in phage isolate genomes relative to reference genomes of both known prophages present in MPAO1 (Pf4 and Pf6). Prophage reference genomes were generated by first extracting the known sequence of these phages from the reference genomes of PAO1 (for Pf4; NC_002516.2) and MPAO1 (for Pf6; NZ_CP027857.1). Prophage reference genomes were annotated using *pharokka* (Bouras et al., 2023) to identify coding regions and gene names, locus IDs, and gene functions were adjusted based on relevant literature [34, 40]. Variants in phage isolates relative to both reference genomes were called using *breseq* in nanopore mode [41]. Finally, a phylogenetic tree of hyperactive phage isolates from population HD7 was generated using IQ-TREE [42] and visualized with *ggtree* [43].

For two phage isolates, a high number of genetic variants were detected. To investigate whether these isolates were recombinants of Pf4 and Pf6, the raw reads from four phage isolates (two putative recombinants, two non-recombinants) were mapped to Pf4 and Pf6 simultaneously using *bwa*. Recombinant phage were identified by searching for regions mapping to both reference phage genomes using the raw reads of single phage isolates.

### Phage virulence measurement

MPAO1 was grown in microplate growth curves as described above in the presence vs. absence of phage isolates (multiplicity of infection ∼ 0.01). Reduced AUC in the presence of phage isolates was used as a measure of virulence.

### Endpoint bacterial population sequencing

Endpoint bacterial populations and ancestral strains were revived from glycerol stocks in 23-h cultures. These cultures were then transferred to fresh media using dilution factors corresponding to their evolved treatment and were grown for an additional 23 h. The second growth step was intended to minimize any shifts in allele frequencies resulting from growth in the presence of glycerol from the frozen stocks. We extracted genomic DNA from these cultures using the protocol for Gram negative bacteria of a DNeasy Blood and Tissue DNA extraction kit (Qiagen, 69504). Short-read sequencing on extracted DNA was performed by SeqCenter on a NovaSeq X Plus System (Illumina), producing 2 x 151 bp paired-end reads. Raw sequencing reads are available on the NCBI Sequence Read Archive (PRJNA1420075).

*fastp* [38] was used to filter raw reads for quality using default parameters (Q >= 15, % unqualified bases >= 40%, minimum read length = 15 bp) and downsample reads to an appropriate sequencing depth for analysis (downsampled to ∼250-fold coverage for populations, ∼50-fold coverage for ancestors). Then, variants in the sequenced populations relative to the MPAO1 reference genome (NCBI Reference Sequence: NZ_CP027857.1) were called using *breseq* in polymorphism mode (frequency cutoff = 0.05). Summary data for all variants in endpoint populations was then filtered for variants in genes related to type IV pilus biosynthesis and function.

### Generating endpoint isolate library

To investigate the phenotypic consequences of hyperactive phage emergence, a library of three isolates from each endpoint population was generated. Isolates were randomly selected from separate colonies generated by plating frozen stocks of endpoint populations on LB agar plates. Isolates were re-streaked three times to ensure they were isogenic and frozen stocks of each were generated by freezing overnight cultures in 20% glycerol at −80 °C.

### Twitch assay

Each evolved isolate, ancestor, and the negative control strain (MPAO1ΔpilA) was grown in a 1.5 mL LB liquid culture for 23 h. Subsequently, sterile wooden toothpicks were used to inoculate each strain into the center of a freshly poured LB plate (10 g/L agar). Plates were then incubated upside down for 48 h at 37 °C. Following incubation, agar was removed and we stained the bottom of each plate using 0.1% w/v crystal violet. Plates were scanned using an Epson Perfection V850 Pro scanner. ImageJ was used to manually outline the stained region, which we used to measure the twitch area. We measured the twitch area of each strain in a single technical replicate on each of 5 days (n = 5).

### Virulent phage susceptibility assay

Lawns of each endpoint isolate, both ancestors, MPAO1ΔpilA, and MPAO1 algC::Tn were generated on LB agar plates as described above, but with 7.5 g/L LB top agar. High titer stocks of two virulent phages (LPS-5 and ΦKMV) were then plated at dilution factors ranging from 10^0^-10^7^ in 10 µL spots. The efficiency of plating (EOP) of each phage on each strain was calculated using the relevant reference strain (MPAO1 or LacZ; Supplementary Methods equation 3).

### Statistics

All statistical analyses were performed in R (version 4.2.0). Our modeling approach was to use linear mixed-effects models (*lme4* package) [44] to compare phenotypes across groups while controlling for factors like date and population ID using random effects where appropriate. Type III ANOVAs were used for overall significance testing (type III Wald χ² tests) of main effects and post-hoc contrasts of relevant groups were carried out using the *emmeans* package. Where appropriate, Bonferroni corrections were used to adjust *P* values from post-hoc contracts. Please see Supplementary File 1 for more detailed information on the specific statistical models used. All non-sequencing data used in this study are included in supplementary files associated with this article.

## Results

### Adaptation to organism-modified environments across population densities

We tracked the population density of our lines at regular intervals during experimental passaging (Figure 1B). End-of-passage population density was similar between HD and LD lines (ANOVA, *P* = 0.64) and modestly increased, on average, by the end of the experiment (ANOVA, *P* < 0.001; est. ± se = 0.18 ± 0.019; Supplementary Figure 3). Despite the general trend of increasing population density over time, we observed transient crashes in population density in most of our lines across both treatments (Figure 1B). Furthermore, the average performance of endpoint populations in microplate growth curves was lower than that of the ancestral strain in the media in which they were passaged (ANOVA post-hoc contrast, *t*_112_ > 9.4, *P* < 0.0001; Figure 1C). These results suggest that ecological and evolutionary dynamics may have been impacted by the production of an inhibitory factor by evolving populations in both passaging regimes.

To investigate adaptation to organism-modified environments, we measured the performance of ancestral and endpoint populations in the presence of filtrate sampled from the same experimental line across evolutionary timepoints. We quantified the effect of filtrate on performance using the relative area-under-the-curve (rAUC) of bacteria grown in the presence of filtrate compared to growth in unmodified media. We tested for adaptation of endpoint populations to organism-modified environments by comparing the rAUC of endpoint and ancestral populations across each filtrate. For example, the endpoint population of line HD7 had a relative growth advantage over the ancestral strain in filtrate from that line’s evolutionary midpoint, suggesting this population was well-adapted to environmental modification by a population sampled from its evolutionary past (ANOVA post-hoc contrast, *t*_29_ = −22.3, *P* < 0.0001; Figure 1D).

We found that endpoint populations had relatively high rAUC in filtrate generated from at least one evolutionary timepoint in 13/16 populations (ANOVA post-hoc contrasts, *P* < 0.05; 6/8 LD populations, 7/8 HD populations; Figure 1E). This pattern was not due to endpoint populations adapting to growth in filtrate in general, as we did not detect an advantage of any endpoint populations in ancestral filtrate (ancestor ≥ endpoint populations in ancestral filtrate in all lines; Supplementary Figure 4).

Although we failed to detect a statistically significant advantage of filtrate in two LD populations (LD3 and LD7), this appeared to result from variable effects of filtrate across days (Supplementary Figure 5). Despite this day-to-day variation, it was clear that these populations were well-adapted to environmental modification by their ancestors, and so we included them among the 13 endpoint populations for which we detected significant advantages in filtrate.

Elevated rAUC in endpoint populations could be driven by any combination of filtrate inhibiting the ancestor or benefiting endpoint populations. We found that elevated rAUC in endpoint populations was overwhelmingly caused by filtrates inhibiting the ancestor (Figure 1F). Relatively high rAUC in endpoint populations was associated with decreased performance of the ancestral strain in all but one case (filtrate from HD2_1000; Figure 1Fi). Conversely, the performance of endpoint populations did not strongly vary across filtrates (Figure 1Fii).

Overall, these results indicated that 15/16 populations evolved to produce one or more filterable factor that inhibited the growth of the ancestral strain. This process did not strongly vary according to population density, as most populations across both treatments showed this pattern (LD: 8/8, HD: 7/8; Fisher’s exact test, *P* = 1). Furthermore, our results suggested that the production of inhibitory environmental factors drove an eco-evolutionary feedback, as endpoint populations were well-adapted to this organism-caused inhibitory environmental change.

### Hyperactive bacteriophages are responsible for inhibitory filtrate

We hypothesized that the inhibitory effect of filtrate on the ancestral strain (MPAO1) was caused by the emergence of hyperactive filamentous phages. Throughout, we refer to phages with repressor mutations as “hyperactive” instead of the more widely used “superinfective”, as recent evidence suggests that Pf phages do not always confer superinfection exclusion to their hosts [45].

We tested for the presence of hyperactive Pf phages by plating each filtrate on lawns of MPAO1 and checking for the presence of phage plaques, which are only produced by hyperactive variants of Pf phages. As expected, we detected phage in all inhibitory filtrates (Figure 2AB; Supplementary Figure 6). Conversely, we did not detect phage in the vast majority of non-inhibitory filtrates (Figure 2AB; Supplementary Figure 6). Furthermore, phage presence was associated with reduced rAUC of ancestral populations when grown in filtrate (ANOVA post-hoc contrast, *t*_625_ = 51.3, *P* < 0.0001), whereas phage presence had no effect on the performance of endpoint populations (ANOVA post-hoc contrast, *t*_625_ = −0.84, *P* = 0.40; Figure 2B). The latter result was consistent with endpoint populations having evolved resistance to phage they encountered previously in their evolutionary history.

**Figure 2:**
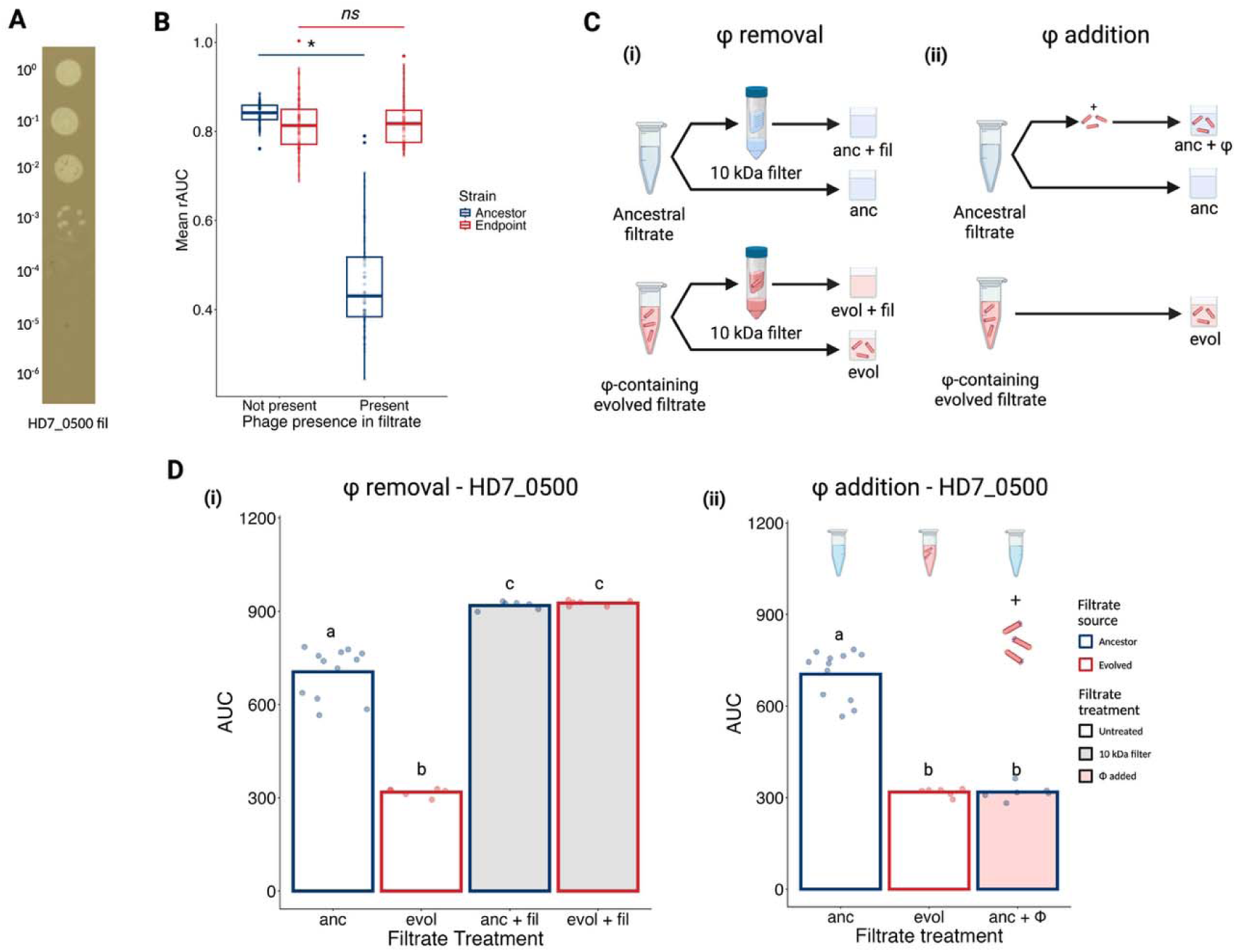
Phage presence explains inhibitory filtrate. **A**: Dilution series of filtrate from HD7_0500 plated on lawn of MPAO1. **B**: rAUC of ancestral and endpoint bacteria in filtrate with or without detectable hyperactive phage. * indicates a statistically significant difference across groups, whereas *ns* indicates no statistically detectable difference. **C**: Design of phage removal/addition experiment. **D**: Phage removal (i) and addition (ii) results for HD7_0500, which is a representative example of the filtrates tested (Supplementary Figure 7). i) AUC of MPAO1 when grown in ancestral (anc) or HD7_0500 (evol) filtrate that was either untreated or passed through a 10 kDa centrifugal filter to remove phage (+ fil). ii) AUC of MPAO1 in ancestral (anc) or HD7_0500 (evol) filtrate, as well as ancestral filtrate with a mix of phage isolated from HD7_0500 filtrate spiked in (+ Φ). Letters indicate statistically distinguishable groups. Created in BioRender. Turner, P. (2026) https://BioRender.com/tfxanrf. Alt text: Multipaneled figure demonstrating that the presence of hyperactive filamentous phages is responsible for the inhibitory effect of some filtrates on the ancestral bacterial strain via reanalysis of growth curve data and experimental removal and addition of hyperactive phages from/into filtrate.

Hyperactive phage were never detected in filtrate from population HD8 (Pf_hyp_-) but were detected in filtrate across all other populations (Pf_hyp_+). Hyperactive phage persisted in Pf_hyp_+ populations for up to 750 bacterial generations but were detected in endpoint filtrate in only two populations (HD7 and HD4; Supplementary Figure 6). This suggested that the evolution of phage resistance in bacterial populations tended to drive hyperactive phage populations extinct, although long term persistence was also possible.

Although hyperactive phage presence was strongly associated with inhibitory filtrate, it was possible that another unknown factor caused both inhibition and hyperactive phage production. To strengthen the causal link between phage presence and inhibition, we tested whether phage presence was necessary and sufficient to explain the decreased performance of MPAO1 in phage-containing filtrate (Figure 2C). We focussed on phage containing filtrates from two populations from each treatment that varied in the extent to which they inhibited MPAO1 (strong inhibition: HD7 and LD8, weak inhibition: HD1 and LD6; Figure 1Ei). Using 10 kDa centrifugal filters, we successfully removed all phage from phage-containing filtrates (Supplementary Figure 2). We found that centrifugal filtration of ancestral and phage-containing filtrate eliminated the difference in performance of MPAO1 across filtrates, suggesting that phage presence was necessary to cause the inhibitory effect (ANOVA post-hoc contrast, *t*_128_ = −0.36, *P* = 0.72; Figure 2Di, Supplementary Figure 7A). Furthermore, spiking in a mix of three contemporaneous phage isolates into ancestral filtrate caused quantitatively similar inhibition as phage-containing filtrate generated by evolved populations (ANOVA, χ_1_² = 3.55, *P* = 0.060; Figure 2Dii, Supplementary Figure 7B). Collectively, these results suggested that hyperactive phage were necessary and sufficient to explain inhibitory filtrate.

### Hyperactive phage isolates descend from both Pf4 and Pf6 across populations

We next sought to determine the identity of the hyperactive phages we detected across filtrates. We focussed on filtrates from the same four populations used in the phage removal/addition experiments described above. We collected and attempted to sequence three phage isolates from each timepoint at which phages were detected in these focal populations (10 total timepoints, 30 phage isolates). 27/30 phage isolates were successfully sequenced.

Phage species varied across populations (Figure 3A). Phage isolated from HD7 and LD8 filtrate descended from Pf4 (Figure 3Ai), whereas phage isolated from HD1 and LD6 filtrate were Pf6-derived (Figure 3Aii). Consistent with previous studies, every isolate we examined either had a deletion in the phage repressor gene or a mutation in the intergenic region upstream of the phage repressor (Figure 3A) [31, 32]. As in prior studies, these mutations likely drove the emergence of hyperactive phages from initially avirulent prophages present in the MPAO1 genome.

**Figure 3:**
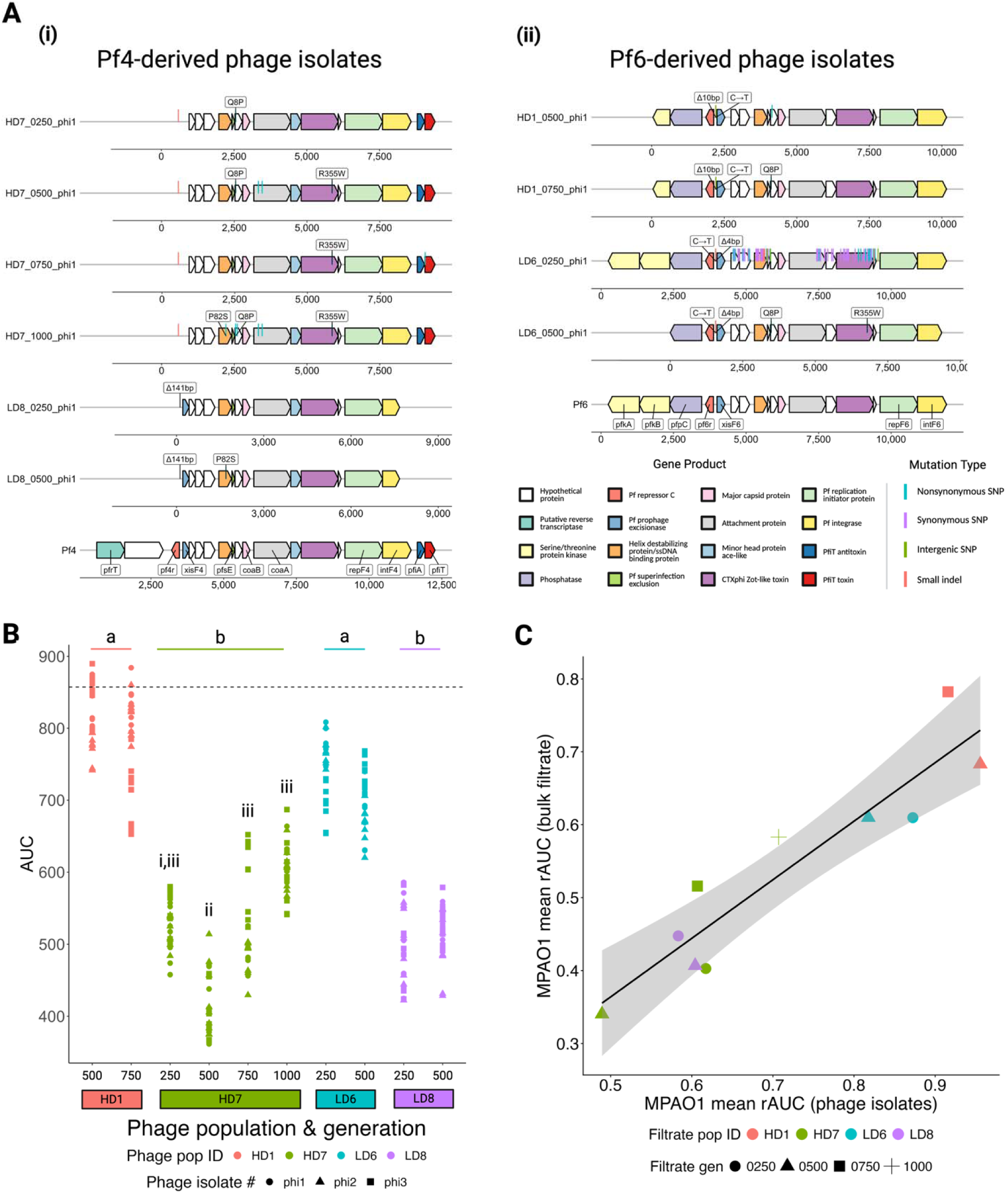
Mutations in both Pf4 and Pf6 are responsible for hyperactive phage emergence across populations. **A**: Annotated genomes of subset of i) Pf4-derived and ii) Pf6-derived hyperactive phage isolates. Annotated reference genomes of wildtype Pf4 and Pf6 are shown at the bottom of each subfigure. Mutations relative to reference genome are indicated by vertical bars. Mutations in the intergenic region upstream of the phage repressor or that occurred across multiple populations are highlighted above phage isolate genomes. Missing genes in phage genomes indicate large deletions. **B**: AUC of MPAO1 grown in the presence of phages isolated across populations and evolutionary timepoints (n = 6). Dashed line indicates average AUC of MPAO1 in the absence of phage. Letters indicate significant differences in AUC across phage populations (a, b), or within population HD7 (i, ii, iii). **C**: Relationship between virulence of phage isolates from a given filtrate and the extent of inhibition caused by that filtrate in growth curves (rAUC of MPAO1). Each point pools the mean virulence of the three phage isolates collected from a given filtrate. Linear relationship shown in black trendline with standard error shown in shaded grey region. Created in BioRender. Turner, P. (2026) https://BioRender.com/frpd3q1. Alt text: Multipaneled figure summarizing genetic variants in hyperactive Pf phages isolated from inhibitory filtrates, how virulence varies among these isolates, and the relationship between MPAO1 inhibition in the presence of bulk filtrate vs. phages isolated from the same filtrate.

### Hyperactive phage genomes contain large deletions, parallel nonsynonymous SNPs

Beyond phage repressor mutations, hyperactive phage isolates contained several other parallel genetic variants. For instance, at least one phage isolate from each timepoint examined contained a large deletion (> 1kb) in the accessory genome (Figure 3A). All Pf4-derived isolates had deleted the region upstream of *pf4r*, which contains genes encoding a reverse transcriptase (*pfrT*) and a hypothetical protein (Figure 3Ai). Further, isolates from LD8-filtrate also had deletions spanning the other end of the genome which encompassed the *pfiAT* toxin-antitoxin system, as well as the 3’-end of the integrase gene (*intF4;* Figure 3Ai). Similarly, Pf6-derived isolates from each timepoint had deletions spanning *pfkA* and all or part of *pfkB*, which encode serine/threonine kinases that act as toxins in a recently described kinase-kinase-phosphatase toxin-antitoxin system (Figure 3Aii) [34]. Only 2/27 of sequenced isolates contained no large deletions (LD6_0250_phi1 and LD6_0250_phi2). These two isolates appeared to be recombinants of Pf4 and Pf6, possessing the accessory genome of Pf6 and core genomes with alternating regions of greater similarity to Pf4 and Pf6 (Supplementary Figure 8).

We also detected three nonsynonymous single-nucleotide polymorphisms (NS SNPs) that were shared across multiple phage populations. Each population of Pf4-derived phage had the same NS SNP (P82S) in PA0720, which encodes a single-stranded DNA binding protein, in at least one isolate (Figure 3Ai). In a related filamentous phage, M13, a homologous gene encodes the protein pV, which binds and stabilizes the ssDNA phage genome and facilitates its transport to the inner membrane for viral assembly [46]. Also, some isolates from HD7 and LD6 shared a NS SNP (R355W) in the gene encoding the CTXΦ Zot-like toxin, which is essential for the assembly and export of virions in another related filamentous phage, f1 (Figure 3A) [47]. Finally, some isolates from HD7, HD1, and LD6 all share a NS SNP (Q8P) in PA0722, a hypothetical gene thought to encode a protein with functional similarity to the minor capsid protein pIX in f1(Figure 3A) [48, 49]. The latter two mutations were shared across isolates derived from different phage species, implying strong selection on these loci. All detected genetic variants in hyperactive phage isolates are available in Supplementary Table 1.

### Hyperactive Pf phage isolates vary in virulence across populations and over generations

Phage isolates varied in virulence on MPAO1 (Figure 3B). Pf6-derived isolates (HD1, LD6) were less virulent than Pf4-derived isolates (HD7, LD8), on average (ANOVA post-hoc contrast; |t_6_| > 3.78, P < 0.05; Figure 3B). Further, isolates from HD7 filtrate varied in virulence across filtrate generation. These phages initially increased in virulence between generation 250 and 500 (ANOVA post-hoc contrast, AUC_gen250_ > AUC_gen500_, *t*_8_ = 3.72, *P* = 0.024), which was followed by an attenuation of virulence from generation 500 to 1000 (ANOVA post-hoc contrast, AUC_gen500_ < AUC_gen750_ = AUC_gen1000_, |*t*_8_| > 3.46, *P* < 0.05; Figure 3B). When plotted on a phylogeny of HD7-derived Pf4 isolates, these transitions in virulence are coincident with SNPs in two genes, PA0720 and PA0722, both of which are involved in the initiation of viral assembly (Supplementary Figure 10) [50].

The extent of inhibition of MPAO1 in bulk filtrate was positively associated with the average virulence of phages isolated from the same filtrate (linear mixed-effects model, *t*_7_ = 5.51, *P* = 0.0009; Figure 3C). Conversely, we detected no relationship between filtrate inhibition and phage titer (linear mixed-effects model, *t*_7_ = -0.60, *P* = 0.57; Supplementary Figure 9). The latter results suggested that the variation in rAUC we observed across filtrates was explained by genetic variation in virulence across phage populations, rather than phage density.

### Emergence of hyperactive Pf phages associated with mutations in TIVP-related genes

We hypothesized that the emergence of hyperactive Pf phages would drive the evolution of phage resistance via mutations in genes related to the type IV pilus (TIVP), the cell-surface receptor for Pf phages. As expected, all Pf_hyp_+ populations had a high frequency (> 50%) of mutations in genes involved in TIVP biosynthesis and function (Figure 4). In these populations, mutations in *pilR* (7/15 populations), *pilS* (7/15), and *pilB* (5/15) were most common (Figure 4). Conversely, we did not detect mutations in any TIVP-related genes in HD8, the Pf_hyp_-population, suggesting these mutations were not strongly favoured in the absence of hyperactive phages (Figure 4).

**Figure 4:**
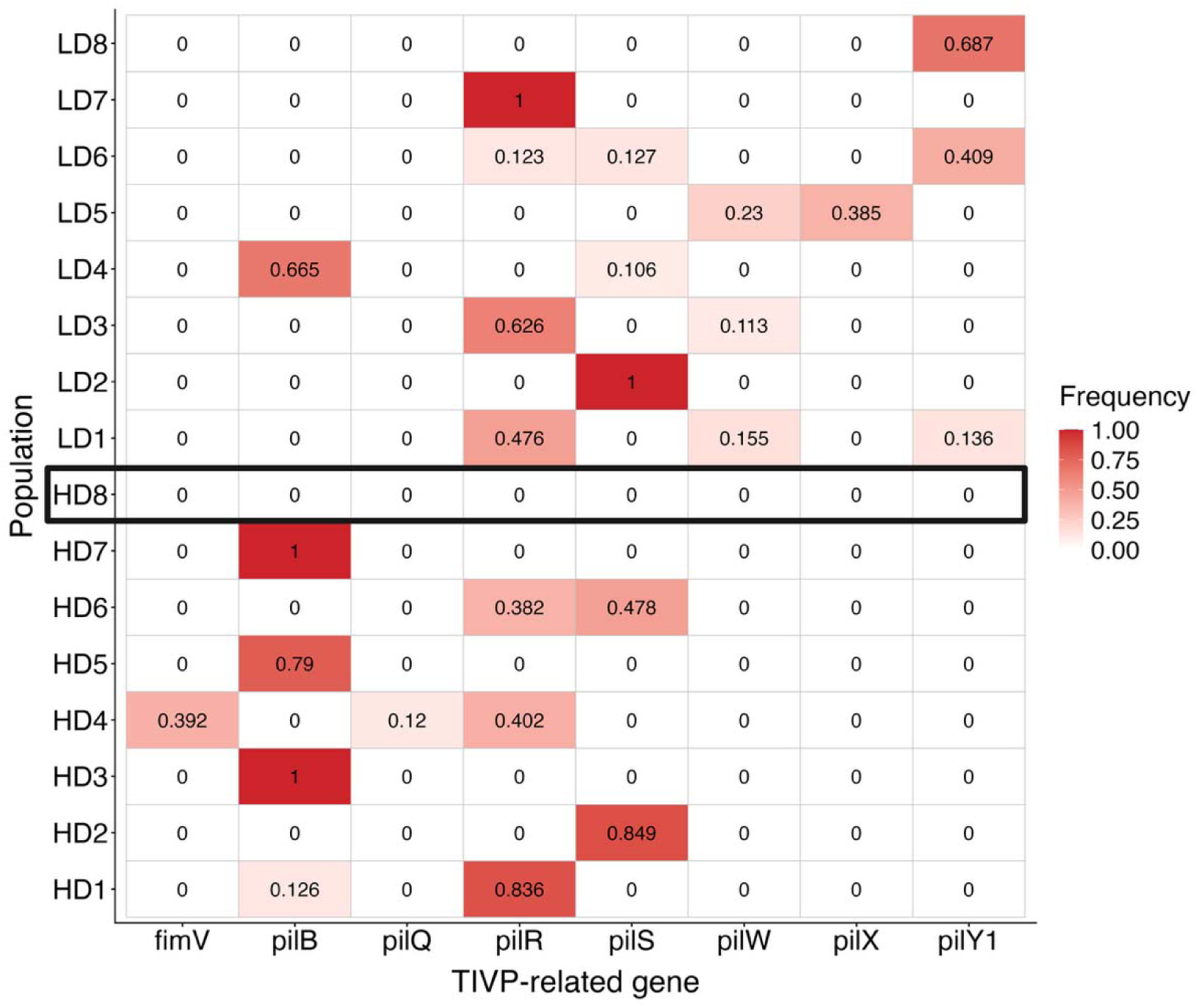
Hyperactive phage emergence drives mutations in genes related to phage host receptor across populations. Total frequency of mutations in TIVP-related genes in endpoint populations. Bolded rectangle highlights HD8, the population in which hyperactive phage were never detected (Pf_hyp_-). Mutations in genes unrelated to TIVP not shown. Created in BioRender. Turner, P. (2026) https://BioRender.com/8q9418s. Alt text: Tile plot summarizing the frequency of mutations in TIVP-related genes across endpoint populations.

We also examined whether endpoint populations contained mutations in either prophage region, which revealed that all endpoint populations still carried the ancestral sequence of Pf4 and Pf6. This was consistent with a model in which the cells from which hyperactive phages initially emerged did not persist at sufficient frequencies to be detected in endpoint population sequencing.

### Emergence of hyperactive Pf phages associated with decreased twitch motility, resistance to a TIVP-targeting virulent phage

To explore the pleiotropic consequences of phage resistance, we first measured twitch motility, which requires functional TIVP, in three bacterial isolates from each endpoint population. Twitch motility in isolates from Pf_hyp_+ populations was significantly lower than in the two ancestral strains (ANOVA post-hoc contrast, *t*_246_ > 7.04, *P* < 0.0001) and was indistinguishable from MPAO1ΔpilA, which is incapable of twitch motility (ANOVA post-hoc contrast, *t*_246_ = 0.137, *P* = 1.00; Figure 5A). Conversely, twitch motility was similar in isolates from HD8 and its ancestral strain (LacZ; ANOVA post-hoc contrast, *t*_246_ = 0.016, *P* = 1.00) and was significantly higher than both MPAO1ΔpilA and isolates from Pf_hyp_+ populations (ANOVA post-hoc contrast, |*t*_246_| > 6.02 *P* < 0.0001, Figure 5A).

**Figure 5:**
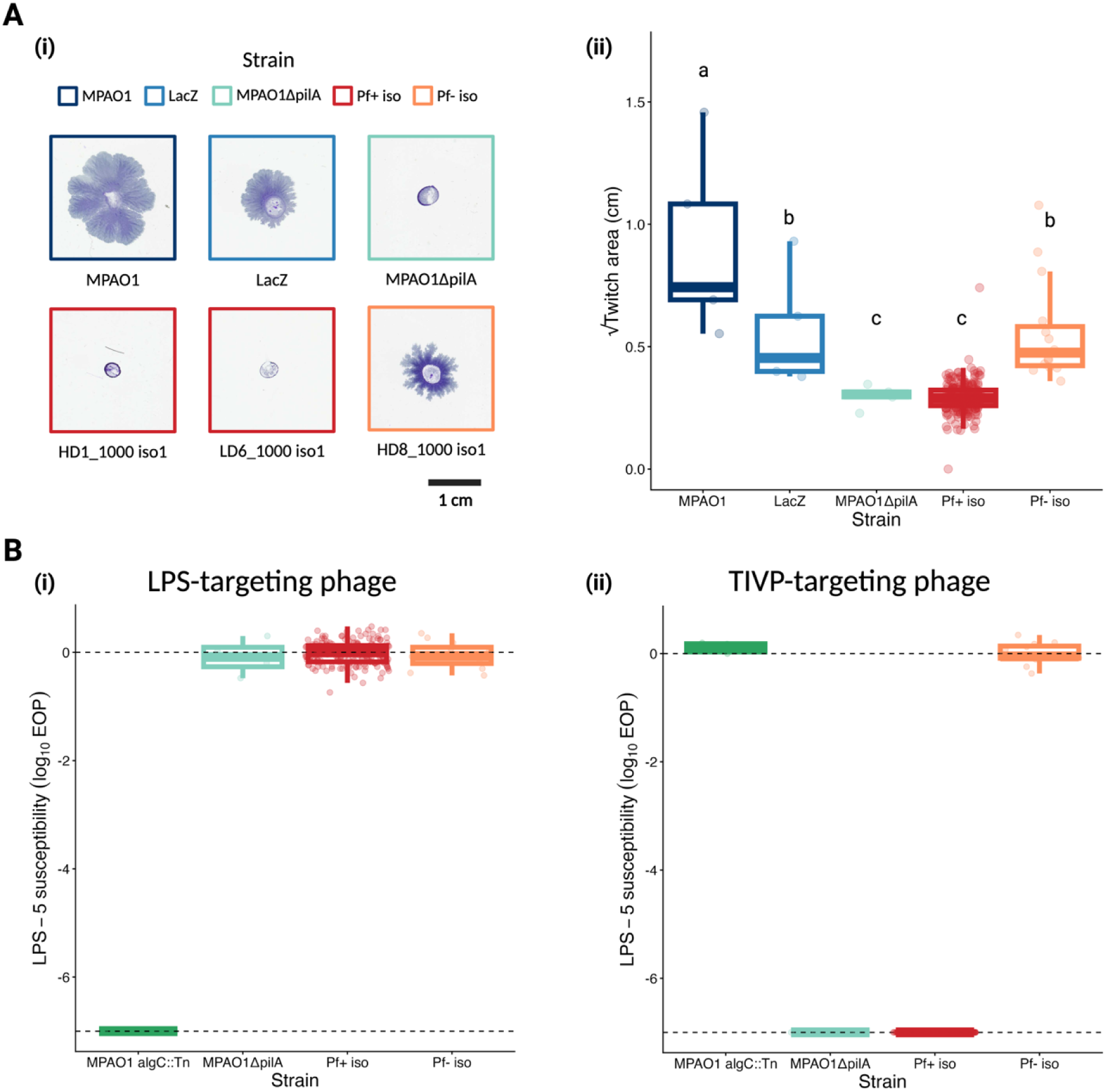
Hyperactive phage emergence drives decreased twitch motility and resistance to TIVP-targeting virulent phage. **A**: Twitch motility of endpoint and ancestral bacterial isolates. i) Example scans of twitch plates across isolates. ii) Twitch area across isolates (three isolates per endpoint population, n = 5 per isolate). Letters denote significant differences in twitch motility across strains. **B**: Efficiency of plating (EOP) of i) an LPS-targeting virulent phage (LPS-5) or ii) a TIVP-targeting virulent phage (ΦKMV) on endpoint isolates relative to MPAO1 (three isolates per endpoint population, n = 4 per isolate). Upper dashed line indicates equal infectivity on MPAO1 and the endpoint isolate. Lower dashed line indicates the lower limit of detection. Created in BioRender. Turner, P. (2026) https://BioRender.com/p1nqkqi. Alt text: Multipaneled figure demonstrating that bacterial isolates from populations in which hyperactive filamentous phage emerged had reduced twitch motility and reduced susceptibility to a TIVP-targeting virulent phage.

We also measured the susceptibility of endpoint isolates to two MPAO1-infecting virulent (strictly lytic) phages which both have shown potential for use in phage therapy to treat *Pa* infections (LPS-5 and ΦKMV) [51, 52]. We used two phages with different cell surface receptors to test whether any changes in phage susceptibility were general, and thus protective against both phages (LPS-5 and ΦKMV), or specific, and thus protective against only the TIVP-targeting phage (ΦKMV). We found that endpoint isolates from both Pf_hyp_+ populations and Pf_hyp_-populations remained susceptible to LPS-5 (Figure 5Bi). Conversely, whereas isolates from the Pf_hyp_-population remained susceptible to ΦKMV, isolates from Pf_hyp_+ populations were resistant to ΦKMV (Figure 5Bii). Taken together, these results suggest that the evolution of resistance to hyperactive Pf phages pleiotropically drove decreased twitch motility and resistance to a virulent TIVP-targeting phage.

## Discussion

Microbial populations strongly shape their environment, which can re-route adaptation toward organism-generated fitness optima. However, the conditions that promote this mode of adaptation are unclear. Here, we explored whether high population density, by magnifying niche construction, increases the frequency with which experimentally evolved populations of the bacterium *Pseudomonas aeruginosa* (*Pa*) MPAO1 adapt to organism-constructed environments. We quantified this form of adaptation by measuring the relative performance of endpoint vs. ancestral populations in filtrate generated by evolved populations sampled across evolutionary timepoints.

Unexpectedly, we found that populations adapted to organism-constructed environments in nearly every replicate population regardless of population density (Figure 1). In each case, this was caused by the presence of hyperactive filamentous phages in evolved filtrate (Figure 2), which emerged from one of two prophages present in MPAO1’s genome (Pf4 or Pf6) during experimental passaging (Figure 3). The emergence of hyperactive phages, in turn, drove the evolution of phage resistance via mutations in type IV pilus (TIVP)-related genes (Figure 4), which explains the performance advantage of endpoint populations in hyperactive phage-containing filtrate. Thus, the reason our results differed from expectations seems clear: unlike other agents of niche construction like toxins and metabolites, hyperactive phages are self-amplifying, capable of horizontal spread, and evolve independently from their host. In our case, these features allowed hyperactive phages to reach high density and, consequently, impose strong selection regardless of host population density or their negative impact on host fitness (Figure 6). Our data imply that prophage evolution deserves further consideration as a driver of niche construction in bacterial populations.

**Figure 6:**
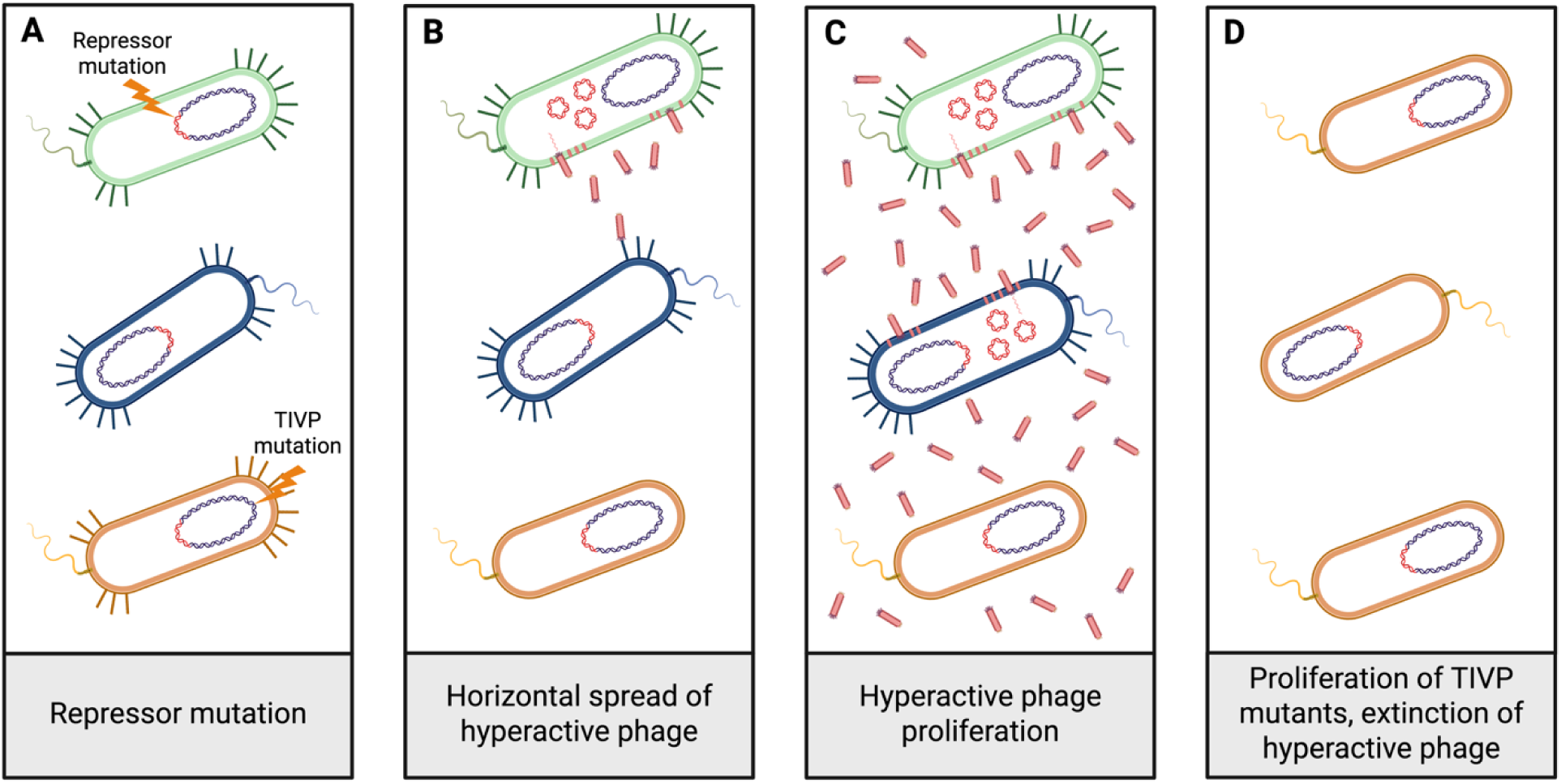
Conceptual overview of eco-evolutionary feedback observed in both HD and LD populations. Colours represent lineages of bacteria in a single population. Bacterial genomes represented with a single prophage region (in red) for simplicity. **A**: A mutation disabling the prophage repressor occurs in the green lineage. At an unknown timepoint, a mutation disabling the type IV pilus (TIVP) occurs in a different lineage (in orange). **B**: The repressor mutation causes hyperactive phage production that spreads to hosts in another lineage with functional TIVP (in blue). **C**: Hyperactive phage proliferate in the media, infecting most cells. **D**: By reducing the growth rate of their hosts, hyperactive phage selectively favour TIVP mutants, ultimately driving their own extinction. The ancestral prophage persists unaltered in the surviving lineage. Created in BioRender. Turner, P. (2026) https://BioRender.com/1ciha2y. Alt text: Conceptual overview of eco-evolutionary feedback observed in the present study. Bacterial lineages are represented as differently coloured cells, one of which sweeps to fixation as a result of a TIVP mutation that confers resistance to hyperactive filamentous phage which are produced by neighbouring cells.

Our results raise the question of how often filamentous phages drive eco-evolutionary feedbacks in bacterial populations. Filamentous phages are relatively common, as they are found in ∼7% of prokaryotic genomes [53]. Our work and others’ suggest that these phages can evolve higher virulence and subsequently drive bacterial evolution in various bacterial strains and species and with differing environmental contexts [30, 34, 54–57]. For example, Renda et al. (2015, 2016) found that a filamentous prophage in *Acinetobacter baylyi* evolved hyperactivity during experimental passaging, selecting for receptor-mediated phage resistance in its host population. The evolution of resistance to hyperactive filamentous phages also appears to drive trade-offs across systems. Just as we found phage resistance pleiotropically drove reduced motility and decreased susceptibility to a TIVP-targeting virulent phage (Figure 5), Renda et al. (2016) found that phage resistance diminished the ability of the host bacteria to take up extracellular DNA, an important mechanism of horizontal gene transfer. Collectively, these results suggest that filamentous phages should be investigated further for their potential to evolve hyperactivity and drive eco-evolutionary feedbacks with pleiotropic consequences. Whether filamentous phages tend to drive such dynamics in natural systems offers an exciting opportunity for future work in environmental microbiology.

Whether and how Pf phages shape *Pa* evolution during infection should be a focus for further research. *Pa* is an opportunistic pathogen that causes recalcitrant lung infections in immunocompromised individuals, including people with cystic fibrosis, in which viable bacterial densities can exceed 5×10^9^ CFU/mL [58–60]. After colonizing the lung, populations of *Pa* undergo characteristic pathoadaptation, which often involves the loss of TIVP-dependent motility [60–62]. Although loss of twitch motility is typically attributed to selection for resistance to phagocytosis by macrophages [61, 63], another non-mutually exclusive possibility is that, similar to what we observed here, loss of twitching during chronic infections is driven by the emergence of hyperactive Pf phages. Consistent with this hypothesis, *Pa* strains isolated from the lungs of pwCF often carry Pf phages and Pf virions are often found at high densities in the sputum of pwCF [64, 65]. Future work should investigate the link between Pf phage carriage and TIVP loss during pathoadaptation. If such a link is detected, it could make pathoadaptation, and therefore both disease severity and treatment efficacy, more predictable. In particular, the relationship between Pf carriage and the susceptibility of pathogenic bacteria to virulent phages deserves further attention (Figure 5B), as phage therapy offers an emerging strategy for treating chronic *Pa* lung infections [52, 66]. Beyond potentially driving TIVP loss, Pf phages also play a role in structuring *Pa* biofilms, contribute to *Pa* virulence, and modulate the human immune system during *Pa* infections [33]. Given the importance of Pf phages to these aspects of *Pa* infections, more work into their evolution in the context of infections is warranted.

Sequencing of hyperactive phages that emerged in our experiment revealed a pattern of parallel evolution across replicate populations and, strikingly, even across phage species (Figure 3A). Parallel evolution is a telltale sign that populations have experienced strong and similar selective pressures, raising the question of which selective pressures drove the changes we observed. One possibility is that strong selection arose from the dramatic change in phage transmission mode caused by the initial repressor mutations. Filamentous phage, like many parasites, maximize their fitness by carefully tailoring their phenotypes according to whether they transmit horizontally (infecting naive hosts) or vertically (from mother to daughter cells) [56, 67, 68]. From this standpoint, repressor mutations cause filamentous phages to transition from specialists in vertical transmission to specialists in horizontal transmission via increased reproduction at the expense of host growth. If horizontal transmission favours different phenotypes than vertical transmission, this may explain the parallel genetic changes we observed across phage populations. For example, vertical transmission often favours the acquisition of genes that benefit the host cell, whereas horizontal transmission in filamentous phages tends to favour genome reduction, which is thought to allow faster replication [57, 69, 70]. This could explain why toxin-antitoxin systems, which often protect host cells from further phage infection in other systems [71], were deleted in parallel across ¾ of our sequenced phage populations (Figure 3A). Similarly, the parallel SNPs we observed in the core phage genomes across populations may represent fine-tuning following the transition to a strictly horizontal mode of transmission (Figure 3A; Supplementary Table 1).

Although more work is needed to dissect the phenotypic consequences of the mutations we detected in Pf isolates, we did observe that phage isolates in one population decreased in virulence over time (Figure 3B). In this population, transitions in virulence were coincident with mutations in PA0720 and PA0722, which are thought to encode a ssDNA-binding protein homologous to pV in the filamentous phage f1 and a minor capsid protein similar to pIX in f1, respectively (Supplementary Figure 10; [48, 50]. In f1, both proteins are involved in the initiation of viral assembly, which is thought to impose membrane stress during Pf phage production in *P.* aeruginosa [30, 49, 72]. Furthermore, compensatory mutations in pIX in response to engineered mutations altering phage packaging signals increased virulence of f1 [49], consistent with our findings that mutations in PA0722 appear to alter virulence in Pf4. Thus, we hypothesize that mutations in PA0720 and PA0722 shape virulence in Pf4 by altering the balance between viral assembly and other, less-costly steps of the Pf4 life cycle, such as genome replication. Regardless of genetic mechanism, the virulence attenuation we observed is consistent with previous work showing that f1 attenuates its virulence when experimentally evolved without the periodic introduction of uninfected hosts [67]. In our experiment, this reduced virulence may have contributed to the longer persistence of hyperactive Pf phages in population HD7 (Supplementary Figure 6), although more data is needed to explain prolonged phage persistence in this population. Overall, our work may suggest that repressor mutations, by shifting the balance between reproductive modes, alter the fitness landscape for filamentous phages in a way that drives strong selection and, thus, parallel evolution. The factors shaping the evolution of virulence in Pf phages should be a priority for further research given their central role in *Pa* biofilms and the potential of virulent phages as antimicrobial agents to treat pathogenic bacterial infections [33, 66].

Our work suggests that eco-evolutionary feedbacks may frequently occur in bacterial populations across densities, however the generality of this conclusion may be limited by our study design. Specifically, our methodological choices may have impacted the likelihood of hyperactive phage emergence by increasing or decreasing Pf induction rate relative to alternative choices. Pf induction is promoted by several factors including high temperature, oxidative stress associated with growth in biofilms, nutrient starvation, and regulatory crosstalk with other prophages [30, 34, 73–75]. Thus, our decision to use a relatively low temperature (30 °C) and shaken planktonic liquid cultures, which should minimize biofilm formation, may have reduced Pf phage induction [30, 74]. Conversely, Pf induction rate may have been promoted by our use of batch culture, which causes daily periods of nutrient starvation, and our use of succinate as a carbon source, which causes oxidative stress in *P. aeruginosa* when provided in excess [73, 76, 77]. However, on the latter point, we note that much higher succinate concentrations than we supplied are needed to cause oxidative stress [30, 77]. Beyond these extrinsic factors, our choice of MPAO1 as the ancestral strain may have promoted hyperactive phage emergence due to regulatory crosstalk between Pf4 and Pf6 [34]. Recent work showed that a kinase-kinase-phosphatase system carried by Pf6 increases induction of both Pf6 and Pf4 in MPAO1 via interactions with the host protein MvaU [34]. Choosing another *P. aeruginosa* strain, particularly one lacking Pf phages altogether, may have allowed non-self-amplifying forms of niche construction to drive adaptation, possibly leading to different conclusions. Finally, our finding that eco-evolutionary feedbacks were similarly frequent across population densities may have been influenced by our use of differential dilution to manipulate population density. By imposing temporal heterogeneity in population density, this approach may have constrained selection on mutations with strong fitness trade-offs across densities. Other approaches, such as using turbidostat systems which control bacterial turbidity, would be useful for imposing constant differences in population density across treatments. Ultimately, experiments using a broader range of strains and species as well as media and passaging designs are needed to further elucidate the impact of population density on the frequency of eco-evolutionary feedbacks.

The selective pressures shaping organisms arise through an interaction between extrinsic factors and the environment-altering actions of the organism itself [1, 6, 7]. Our work failed to detect a connection between population density and adaptation to organism-driven environmental change. Instead, we uncovered a density-independent mechanism through which selective pressures can arise from non-extrinsic factors: the emergence of hyperactive filamentous phage from initially avirulent prophages. Much more work is needed to uncover the general conditions that promote adaptation to non-extrinsic selective pressures. However, our work and others’ [7, 13, 15, 16, 21–23] indicate that models of microbial evolution that assume organisms are only shaped by extrinsic selective pressures are frequently insufficient to explain the outcome of adaptive evolution.

## Acknowledgments

We thank members of the Turner Lab, especially Dallas L. Mould, Kaitlyn E. Kortright, Helen M. Stone, Michael Blazanin, Alita R. Burmeister, and Isaac Larbi Osew for their helpful input on the design and analysis of this study. We also thank Barbara I. Kazmierczak and Roxanne Morris for supplying the bacterial strains used herein. We thank Nanami Kubota for her insight into the analysis of Pf phage sequencing data. We acknowledge generous support from Yale University for the Center for Phage Biology & Therapy and from the Yale Institute for Biospheric Studies (YIBS), which funded this work. CAH was supported by a Gaylord Donnelley Postdoctoral Environmental Fellowship through YIBS and a Postdoctoral Research Fellowship in Biology through the National Science Foundation (Award No. 2109819). NSBH was supported by a National Sciences and Engineering Research Council of Canada Postgraduate Scholarship (CGSD3 - 559651 - 2021).

