## Supplementary Table 1 for "Mutations in filamentous bacteriophages spark eco-evolutionary feedbacks in *Pseudomonas aeruginosa*"

**Supplementary Table 1: Genetic variants detected in hyperactive filamentous phage isolates.**

| Phage Isolate ID | Phage Species | Gene Name | Gene Product | Type | SNP Type | Genetic Change | Amino Acid Change |
|---|---|---|---|---|---|---|---|
| HD1_0500_phi1 | Pf6 | MPAO1_RS28990/MPAO1_RS24230 | transcriptional regulator/hypothetical protein | DEL | | Δ10bp | N/A |
| HD1_0500_phi1 | Pf6 | MPAO1_RS28990/MPAO1_RS24230 | transcriptional regulator/hypothetical protein | SNP | intergenic | C→T | N/A |
| HD1_0500_phi1 | Pf6 | MPAO1_RS24215 | hypothetical protein | SNP | nonsynonymous | A→G | E18G |
| HD1_0750_phi1 | Pf6 | MPAO1_RS28990/MPAO1_RS24230 | transcriptional regulator/hypothetical protein | DEL | | Δ10bp | N/A |
| HD1_0750_phi1 | Pf6 | MPAO1_RS28990/MPAO1_RS24230 | transcriptional regulator/hypothetical protein | SNP | intergenic | C→T | N/A |
| HD1_0750_phi1 | Pf6 | MPAO1_RS24215 | hypothetical protein | SNP | nonsynonymous | A→C | Q8P |
| HD1_0750_phi2 | Pf6 | MPAO1_RS28990/MPAO1_RS24230 | transcriptional regulator/hypothetical protein | DEL | | Δ10bp | N/A |
| HD1_0750_phi2 | Pf6 | MPAO1_RS28990/MPAO1_RS24230 | transcriptional regulator/hypothetical protein | SNP | intergenic | C→T | N/A |
| HD1_0750_phi2 | Pf6 | MPAO1_RS24215 | hypothetical protein | SNP | nonsynonymous | A→C | Q8P |
| HD7_0250_phi1 | Pf4 | PA0714.1/PA0715 | PhrD/polymerase | SNP | intergenic | G→C | N/A |
| HD7_0250_phi1 | Pf4 | [PA0715]–PA0716.1 | [PA0715],PA0716,PA0716.1 | DEL | | Δ2859bp | N/A |
| HD7_0250_phi1 | Pf4 | PA0722 | hypothetical protein | SNP | nonsynonymous | A→C | Q8P |
| HD7_0250_phi2 | Pf4 | PA0714.1/PA0715 | PhrD/polymerase | SNP | intergenic | G→C | N/A |
| HD7_0250_phi2 | Pf4 | PA0722 | hypothetical protein | SNP | nonsynonymous | A→C | Q8P |
| HD7_0250_phi3 | Pf4 | PA0714.1/PA0715 | PhrD/polymerase | SNP | intergenic | G→C | N/A |
| HD7_0250_phi3 | Pf4 | [PA0715]–PA0716.1 | [PA0715],PA0716,PA0716.1 | DEL | | Δ2859bp | N/A |
| HD7_0250_phi3 | Pf4 | PA0722 | hypothetical protein | SNP | nonsynonymous | A→C | Q8P |
| HD7_0250_phi3 | Pf4 | PA0726 | CTXphi Zot-like toxin | SNP | nonsynonymous | C→T | T41I |
| HD7_0500_phi1 | Pf4 | PA0714.1/PA0715 | PhrD/polymerase | SNP | intergenic | G→C | N/A |
| HD7_0500_phi1 | Pf4 | PA0722 | hypothetical protein | SNP | nonsynonymous | A→C | Q8P |
| HD7_0500_phi1 | Pf4 | PA0724 | hypothetical protein | SNP | nonsynonymous | G→A | R53H |
| HD7_0500_phi1 | Pf4 | PA0724 | hypothetical protein | SNP | nonsynonymous | A→G | D95G |
| HD7_0500_phi1 | Pf4 | PA0726 | CTXphi Zot-like toxin | SNP | nonsynonymous | C→T | R355W |
| HD7_0500_phi2 | Pf4 | PA0714.1/PA0715 | PhrD/polymerase | SNP | intergenic | G→C | N/A |
| HD7_0500_phi2 | Pf4 | PA0722 | hypothetical protein | SNP | nonsynonymous | A→C | Q8P |
| HD7_0500_phi2 | Pf4 | PA0724 | hypothetical protein | SNP | nonsynonymous | G→A | R53H |
| HD7_0500_phi2 | Pf4 | PA0724 | hypothetical protein | SNP | nonsynonymous | A→G | D95G |
| HD7_0500_phi2 | Pf4 | PA0726 | CTXphi Zot-like toxin | SNP | nonsynonymous | C→T | R355W |
| HD7_0500_phi3 | Pf4 | PA0714.1/PA0715 | PhrD/polymerase | SNP | intergenic | G→C | N/A |
| HD7_0500_phi3 | Pf4 | PA0722 | hypothetical protein | SNP | nonsynonymous | A→C | Q8P |
| HD7_0500_phi3 | Pf4 | PA0724 | hypothetical protein | SNP | nonsynonymous | G→A | R53H |
| HD7_0500_phi3 | Pf4 | PA0724 | hypothetical protein | SNP | nonsynonymous | A→G | D95G |
| HD7_0500_phi3 | Pf4 | PA0726 | CTXphi Zot-like toxin | SNP | nonsynonymous | C→T | R355W |
| HD7_0750_phi1 | Pf4 | PA0714.1/PA0715 | PhrD/polymerase | SNP | intergenic | G→C | N/A |
| HD7_0750_phi1 | Pf4 | [PA0715]–PA0716.1 | [PA0715],PA0716,PA0716.1 | DEL | | Δ2859bp | N/A |

| | | | | | | | |
|---|---|---|---|---|---|---|---|
| HD7_0750_phi1 | Pf4 | PA0726 | CTXphi Zot-like toxin | SNP | nonsynonymous | C→T | R355W |
| HD7_0750_phi1 | Pf4 | PA0729 | hypothetical protein | SNP | nonsynonymous | C→T | P3L |
| HD7_0750_phi2 | Pf4 | PA0714.1/PA0715 | PhrD/polymerase | SNP | intergenic | G→C | N/A |
| HD7_0750_phi2 | Pf4 | PA0724 | hypothetical protein | SNP | nonsynonymous | G→A | R53H |
| HD7_0750_phi2 | Pf4 | PA0724 | hypothetical protein | SNP | nonsynonymous | A→G | D95G |
| HD7_0750_phi2 | Pf4 | PA0726 | CTXphi Zot-like toxin | SNP | nonsynonymous | C→T | R355W |
| HD7_0750_phi3 | Pf4 | PA0714.1/PA0715 | PhrD/polymerase | SNP | intergenic | G→C | N/A |
| HD7_0750_phi3 | Pf4 | PA0720 | single strand DNA binding protein | SNP | nonsynonymous | C→T | P82S |
| HD7_0750_phi3 | Pf4 | PA0722 | hypothetical protein | SNP | nonsynonymous | A→C | Q8P |
| HD7_0750_phi3 | Pf4 | PA0722 | hypothetical protein | SNP | nonsynonymous | G→T | A27S |
| HD7_0750_phi3 | Pf4 | PA0724 | hypothetical protein | SNP | nonsynonymous | G→A | R53H |
| HD7_0750_phi3 | Pf4 | PA0724 | hypothetical protein | SNP | nonsynonymous | A→G | D95G |
| HD7_0750_phi3 | Pf4 | PA0726 | CTXphi Zot-like toxin | SNP | nonsynonymous | C→T | R355W |
| HD7_1000_phi1 | Pf4 | PA0714.1/PA0715 | PhrD/polymerase | SNP | intergenic | G→C | N/A |
| HD7_1000_phi1 | Pf4 | PA0720 | single strand DNA binding protein | SNP | nonsynonymous | C→T | P82S |
| HD7_1000_phi1 | Pf4 | PA0722 | hypothetical protein | SNP | nonsynonymous | A→C | Q8P |
| HD7_1000_phi1 | Pf4 | PA0722 | hypothetical protein | SNP | nonsynonymous | G→T | A27S |
| HD7_1000_phi1 | Pf4 | PA0724 | hypothetical protein | SNP | nonsynonymous | G→A | R53H |
| HD7_1000_phi1 | Pf4 | PA0724 | hypothetical protein | SNP | nonsynonymous | A→G | D95G |
| HD7_1000_phi1 | Pf4 | PA0726 | CTXphi Zot-like toxin | SNP | nonsynonymous | C→T | R355W |
| HD7_1000_phi2 | Pf4 | PA0714.1/PA0715 | PhrD/polymerase | SNP | intergenic | G→C | N/A |
| HD7_1000_phi2 | Pf4 | [PA0715]–PA0716.1 | [PA0715],PA0716,PA0716.1 | DEL | | Δ2859bp | N/A |
| HD7_1000_phi2 | Pf4 | PA0720 | single strand DNA binding protein | SNP | nonsynonymous | C→T | P82S |
| HD7_1000_phi2 | Pf4 | PA0722 | hypothetical protein | SNP | nonsynonymous | A→C | Q8P |
| HD7_1000_phi2 | Pf4 | PA0722 | hypothetical protein | SNP | nonsynonymous | G→T | A27S |
| HD7_1000_phi2 | Pf4 | PA0724 | hypothetical protein | SNP | nonsynonymous | G→A | R53H |
| HD7_1000_phi2 | Pf4 | PA0724 | hypothetical protein | SNP | nonsynonymous | A→G | D95G |
| HD7_1000_phi2 | Pf4 | PA0726 | CTXphi Zot-like toxin | SNP | nonsynonymous | C→T | R355W |
| HD7_1000_phi3 | Pf4 | PA0714.1/PA0715 | PhrD/polymerase | SNP | intergenic | G→C | N/A |
| HD7_1000_phi3 | Pf4 | [PA0715]–PA0716.1 | [PA0715],PA0716,PA0716.1 | DEL | | Δ2859bp | N/A |
| HD7_1000_phi3 | Pf4 | PA0720 | single strand DNA binding protein | SNP | nonsynonymous | C→T | P82S |
| HD7_1000_phi3 | Pf4 | PA0722 | hypothetical protein | SNP | nonsynonymous | A→C | Q8P |
| HD7_1000_phi3 | Pf4 | PA0722 | hypothetical protein | SNP | nonsynonymous | G→T | A27S |
| HD7_1000_phi3 | Pf4 | PA0724 | hypothetical protein | SNP | nonsynonymous | G→A | R53H |
| HD7_1000_phi3 | Pf4 | PA0724 | hypothetical protein | SNP | nonsynonymous | A→G | D95G |
| HD7_1000_phi3 | Pf4 | PA0726 | CTXphi Zot-like toxin | SNP | nonsynonymous | C→T | R355W |
| LD6_0250_phi1 | Pf6* | MPAO1_RS28990/MPAO1_RS24230 | transcriptional regulator/hypothetical protein | SNP | intergenic | C→T | N/A |
| LD6_0250_phi1 | Pf6* | MPAO1_RS28990/MPAO1_RS24230 | transcriptional regulator/hypothetical protein | DEL | | Δ4bp | N/A |
| LD6_0250_phi1 | Pf6* | MPAO1_RS24225 | hypothetical protein | SNP | nonsynonymous | C→T | A27V |
| LD6_0250_phi1 | Pf6* | MPAO1_RS24225 | hypothetical protein | SNP | synonymous | G→A | P37P |
| LD6_0250_phi1 | Pf6* | MPAO1_RS24225 | hypothetical protein | SNP | synonymous | T→C | D38D |
| LD6_0250_phi1 | Pf6* | MPAO1_RS24225 | hypothetical protein | SNP | synonymous | C→T | C39C |
| LD6_0250_phi1 | Pf6* | MPAO1_RS24225 | hypothetical protein | SNP | nonsynonymous | A→G | D49G |

| | | | | | | | |
|---|---|---|---|---|---|---|---|
| LD6_0250_phi1 | Pf6* | MPAO1_RS24225 | hypothetical protein | SNP | synonymous | A→G | K81K |
| LD6_0250_phi1 | Pf6* | MPAO1_RS24225 | hypothetical protein | SNP | synonymous | T→C | P82P |
| LD6_0250_phi1 | Pf6* | MPAO1_RS24225 | hypothetical protein | SNP | synonymous | C→T | F85F |
| LD6_0250_phi1 | Pf6* | MPAO1_RS24225/ZDSLLIWT_CDS_0006 | hypothetical protein/hypothetical protein | SNP | intergenic | C→A | N/A |
| LD6_0250_phi1 | Pf6* | ZDSLLIWT_CDS_0006 | hypothetical protein | SNP | synonymous | C→T | L2L |
| LD6_0250_phi1 | Pf6* | ZDSLLIWT_CDS_0006 | hypothetical protein | SNP | nonsynonymous | T→C | L36P |
| LD6_0250_phi1 | Pf6* | ZDSLLIWT_CDS_0006 | hypothetical protein | SNP | synonymous | C→G | R46R |
| LD6_0250_phi1 | Pf6* | ZDSLLIWT_CDS_0006 | hypothetical protein | SNP | synonymous | T→G | L60L |
| LD6_0250_phi1 | Pf6* | ZDSLLIWT_CDS_0006 | hypothetical protein | INS | | | N/A |
| LD6_0250_phi1 | Pf6* | ZDSLLIWT_CDS_0006 | hypothetical protein | SNP | synonymous | C→A | G102G |
| LD6_0250_phi1 | Pf6* | ZDSLLIWT_CDS_0006 | hypothetical protein | SNP | nonsynonymous | C→T | T109I |
| LD6_0250_phi1 | Pf6* | ZDSLLIWT_CDS_0006 | hypothetical protein | SNP | nonsynonymous | A→C | T111P |
| LD6_0250_phi1 | Pf6* | ZDSLLIWT_CDS_0006 | hypothetical protein | SNP | synonymous | T→C | T111P |
| LD6_0250_phi1 | Pf6* | ZDSLLIWT_CDS_0006 | hypothetical protein | SNP | nonsynonymous | T→C | *115R |
| LD6_0250_phi1 | Pf6* | MPAO1_RS24220 | single strand DNA binding protein | SNP | synonymous | T→C | T18T |
| LD6_0250_phi1 | Pf6* | MPAO1_RS24220 | single strand DNA binding protein | SNP | synonymous | C→T | Y21Y |
| LD6_0250_phi1 | Pf6* | MPAO1_RS24220 | single strand DNA binding protein | SNP | synonymous | G→A | Q34Q |
| LD6_0250_phi1 | Pf6* | MPAO1_RS24220 | single strand DNA binding protein | SNP | synonymous | C→T | G42G |
| LD6_0250_phi1 | Pf6* | MPAO1_RS24220 | single strand DNA binding protein | SNP | synonymous | C→A | G49G |
| LD6_0250_phi1 | Pf6* | MPAO1_RS24220 | single strand DNA binding protein | SNP | synonymous | A→G | P71P |
| LD6_0250_phi1 | Pf6* | MPAO1_RS24220 | single strand DNA binding protein | SNP | synonymous | C→T | R81R |
| LD6_0250_phi1 | Pf6* | MPAO1_RS24220 | single strand DNA binding protein | SNP | synonymous | T→C | G88G |
| LD6_0250_phi1 | Pf6* | MPAO1_RS24220 | single strand DNA binding protein | SNP | synonymous | A→G | Q94Q |
| LD6_0250_phi1 | Pf6* | MPAO1_RS24220 | single strand DNA binding protein | SNP | synonymous | T→A | L96L |
| LD6_0250_phi1 | Pf6* | MPAO1_RS24220 | single strand DNA binding protein | SNP | synonymous | T→C | G104G |
| LD6_0250_phi1 | Pf6* | MPAO1_RS24220 | single strand DNA binding protein | SUB | | Δ2bp | N/A |
| LD6_0250_phi1 | Pf6* | MPAO1_RS24220 | single strand DNA binding protein | SUB | | Δ2bp | N/A |
| LD6_0250_phi1 | Pf6* | MPAO1_RS24220 | single strand DNA binding protein | SNP | nonsynonymous | T→A | S139T |
| LD6_0250_phi1 | Pf6* | MPAO1_RS24220 | single strand DNA binding protein | SNP | synonymous | C→G | A144A |
| LD6_0250_phi1 | Pf6* | MPAO1_RS24220/ZDSLLIWT_CDS_0008 | single strand DNA binding protein/hypothetical protein | SNP | intergenic | G→C | N/A |
| LD6_0250_phi1 | Pf6* | ZDSLLIWT_CDS_0008 | hypothetical protein | SNP | synonymous | T→C | F8F |
| LD6_0250_phi1 | Pf6* | ZDSLLIWT_CDS_0008 | hypothetical protein | SNP | nonsynonymous | A→G | T29A |
| LD6_0250_phi1 | Pf6* | ZDSLLIWT_CDS_0008/MPAO1_RS24215 | hypothetical protein/hypothetical protein | SNP | intergenic | T→C | N/A |
| LD6_0250_phi1 | Pf6* | MPAO1_RS24205 | hypothetical protein | SNP | synonymous | A→C | P315P |
| LD6_0250_phi1 | Pf6* | MPAO1_RS24205 | hypothetical protein | SNP | synonymous | T→C | P318P |
| LD6_0250_phi1 | Pf6* | MPAO1_RS24205 | hypothetical protein | SNP | synonymous | T→C | T329T |
| LD6_0250_phi1 | Pf6* | MPAO1_RS24205 | hypothetical protein | SNP | nonsynonymous | A→C | E343D |
| LD6_0250_phi1 | Pf6* | MPAO1_RS24205 | hypothetical protein | SNP | synonymous | C→T | D357D |
| LD6_0250_phi1 | Pf6* | MPAO1_RS24205 | hypothetical protein | SNP | synonymous | T→C | G361G |
| LD6_0250_phi1 | Pf6* | MPAO1_RS24205 | hypothetical protein | SNP | synonymous | A→G | L406L |
| LD6_0250_phi1 | Pf6* | MPAO1_RS24205 | hypothetical protein | SNP | synonymous | T→C | A412A |
| LD6_0250_phi1 | Pf6* | ZDSLLIWT_CDS_0012 | minor head protein Ace-like | SNP | synonymous | T→C | D20D |

| | | | | | | | |
|---|---|---|---|---|---|---|---|
| LD6_0250_phi1 | Pf6* | ZDSLLIWT_CDS_0012 | minor head protein Ace-like | SNP | synonymous | T→C | S66S |
| LD6_0250_phi1 | Pf6* | ZDSLLIWT_CDS_0012 | minor head protein Ace-like | SNP | synonymous | G→T | G82G |
| LD6_0250_phi1 | Pf6* | MPAO1_RS24195 | CTXphi Zot-like toxin | SNP | synonymous | C→T | F49F |
| LD6_0250_phi1 | Pf6* | MPAO1_RS24195 | CTXphi Zot-like toxin | SNP | synonymous | G→T | G82G |
| LD6_0250_phi1 | Pf6* | MPAO1_RS24195 | CTXphi Zot-like toxin | SNP | synonymous | G→A | K96K |
| LD6_0250_phi1 | Pf6* | MPAO1_RS24195 | CTXphi Zot-like toxin | SNP | synonymous | A→G | A117A |
| LD6_0250_phi1 | Pf6* | MPAO1_RS24195 | CTXphi Zot-like toxin | SNP | synonymous | C→T | H131H |
| LD6_0250_phi1 | Pf6* | MPAO1_RS24195 | CTXphi Zot-like toxin | SNP | synonymous | C→T | I138I |
| LD6_0250_phi1 | Pf6* | MPAO1_RS24195 | CTXphi Zot-like toxin | SNP | synonymous | A→G | Q215Q |
| LD6_0250_phi1 | Pf6* | MPAO1_RS24195 | CTXphi Zot-like toxin | SNP | nonsynonymous | G→C | G254A |
| LD6_0250_phi1 | Pf6* | MPAO1_RS24195 | CTXphi Zot-like toxin | SNP | nonsynonymous | G→A | A281T |
| LD6_0250_phi1 | Pf6* | MPAO1_RS24195 | CTXphi Zot-like toxin | SNP | synonymous | C→T | L318L |
| LD6_0250_phi1 | Pf6* | MPAO1_RS24195 | CTXphi Zot-like toxin | SNP | synonymous | T→C | L334L |
| LD6_0250_phi1 | Pf6* | MPAO1_RS24195 | CTXphi Zot-like toxin | SNP | nonsynonymous | G→C | E335D |
| LD6_0250_phi1 | Pf6* | MPAO1_RS24195 | CTXphi Zot-like toxin | SNP | synonymous | G→A | V356V |
| LD6_0250_phi1 | Pf6* | MPAO1_RS24195 | CTXphi Zot-like toxin | SNP | nonsynonymous | C→G | D370E |
| LD6_0250_phi1 | Pf6* | MPAO1_RS24195 | CTXphi Zot-like toxin | SNP | nonsynonymous | G→A | A378T |
| LD6_0250_phi1 | Pf6* | MPAO1_RS24195 | CTXphi Zot-like toxin | SNP | nonsynonymous | G→C | A382P |
| LD6_0250_phi1 | Pf6* | MPAO1_RS24195 | CTXphi Zot-like toxin | SNP | nonsynonymous | C→T | A389V |
| LD6_0250_phi1 | Pf6* | MPAO1_RS24195 | CTXphi Zot-like toxin | SNP | synonymous | C→T | V394V |
| LD6_0250_phi1 | Pf6* | MPAO1_RS24195 | CTXphi Zot-like toxin | SNP | synonymous | G→C | A396A |
| LD6_0250_phi1 | Pf6* | MPAO1_RS24195 | CTXphi Zot-like toxin | SNP | nonsynonymous | T→C | V400A |
| LD6_0250_phi1 | Pf6* | MPAO1_RS24195 | CTXphi Zot-like toxin | SNP | synonymous | T→G | G404G |
| LD6_0250_phi1 | Pf6* | MPAO1_RS24195 | CTXphi Zot-like toxin | SNP | nonsynonymous | C→G | H423Q |
| LD6_0250_phi1 | Pf6* | ZDSLLIWT_CDS_0014 | hypothetical protein | SUB | | Δ2bp | N/A |
| LD6_0250_phi1 | Pf6* | ZDSLLIWT_CDS_0014 | hypothetical protein | SNP | nonsynonymous | T→A | S21T |
| LD6_0250_phi1 | Pf6* | ZDSLLIWT_CDS_0014/MPAO1_RS24190 | hypothetical protein/hypothetical protein | SNP | intergenic | T→C | N/A |
| LD6_0250_phi2 | Pf6* | MPAO1_RS28990/MPAO1_RS24230 | transcriptional regulator/hypothetical protein | SNP | intergenic | C→T | N/A |
| LD6_0250_phi2 | Pf6* | MPAO1_RS28990/MPAO1_RS24230 | transcriptional regulator/hypothetical protein | DEL | | Δ4bp | N/A |
| LD6_0250_phi2 | Pf6* | MPAO1_RS24205 | hypothetical protein | SNP | synonymous | T→C | P318P |
| LD6_0250_phi2 | Pf6* | MPAO1_RS24205 | hypothetical protein | SNP | synonymous | T→C | T329T |
| LD6_0250_phi2 | Pf6* | MPAO1_RS24205 | hypothetical protein | SNP | nonsynonymous | A→C | E343D |
| LD6_0250_phi2 | Pf6* | MPAO1_RS24205 | hypothetical protein | SNP | synonymous | C→T | D357D |
| LD6_0250_phi2 | Pf6* | MPAO1_RS24205 | hypothetical protein | SNP | synonymous | T→C | G361G |
| LD6_0250_phi2 | Pf6* | MPAO1_RS24205 | hypothetical protein | SNP | synonymous | A→G | L406L |
| LD6_0250_phi2 | Pf6* | MPAO1_RS24205 | hypothetical protein | SNP | synonymous | T→C | A412A |
| LD6_0250_phi2 | Pf6* | ZDSLLIWT_CDS_0012 | minor head protein Ace-like | SNP | synonymous | T→C | D20D |
| LD6_0250_phi2 | Pf6* | ZDSLLIWT_CDS_0012 | minor head protein Ace-like | SNP | synonymous | T→C | S66S |
| LD6_0250_phi2 | Pf6* | ZDSLLIWT_CDS_0012 | minor head protein Ace-like | SNP | synonymous | G→T | G82G |
| LD6_0250_phi2 | Pf6* | MPAO1_RS24195 | CTXphi Zot-like toxin | SNP | synonymous | C→T | F49F |
| LD6_0250_phi2 | Pf6* | MPAO1_RS24195 | CTXphi Zot-like toxin | SNP | synonymous | G→T | G82G |
| LD6_0250_phi2 | Pf6* | MPAO1_RS24195 | CTXphi Zot-like toxin | SNP | synonymous | G→A | K96K |

| | | | | | | | |
|---|---|---|---|---|---|---|---|
| LD6_0250_phi2 | Pf6* | MPAO1_RS24195 | CTXphi Zot-like toxin | SNP | synonymous | A→G | A117A |
| LD6_0250_phi2 | Pf6* | MPAO1_RS24195 | CTXphi Zot-like toxin | SNP | synonymous | C→T | H131H |
| LD6_0250_phi2 | Pf6* | MPAO1_RS24195 | CTXphi Zot-like toxin | SNP | synonymous | C→T | I138I |
| LD6_0250_phi2 | Pf6* | MPAO1_RS24195 | CTXphi Zot-like toxin | SNP | synonymous | A→G | Q215Q |
| LD6_0250_phi2 | Pf6* | MPAO1_RS24195 | CTXphi Zot-like toxin | SNP | nonsynonymous | G→C | G254A |
| LD6_0250_phi2 | Pf6* | MPAO1_RS24195 | CTXphi Zot-like toxin | SNP | nonsynonymous | G→A | A281T |
| LD6_0250_phi2 | Pf6* | MPAO1_RS24195 | CTXphi Zot-like toxin | SNP | synonymous | C→T | L318L |
| LD6_0250_phi2 | Pf6* | MPAO1_RS24195 | CTXphi Zot-like toxin | SNP | synonymous | T→C | L334L |
| LD6_0250_phi2 | Pf6* | MPAO1_RS24195 | CTXphi Zot-like toxin | SNP | nonsynonymous | G→C | E335D |
| LD6_0250_phi2 | Pf6* | MPAO1_RS24195 | CTXphi Zot-like toxin | SNP | synonymous | G→A | V356V |
| LD6_0250_phi2 | Pf6* | MPAO1_RS24195 | CTXphi Zot-like toxin | SNP | nonsynonymous | C→G | D370E |
| LD6_0250_phi2 | Pf6* | MPAO1_RS24195 | CTXphi Zot-like toxin | SNP | nonsynonymous | G→A | A378T |
| LD6_0250_phi2 | Pf6* | MPAO1_RS24195 | CTXphi Zot-like toxin | SNP | nonsynonymous | G→C | A382P |
| LD6_0250_phi2 | Pf6* | MPAO1_RS24195 | CTXphi Zot-like toxin | SNP | nonsynonymous | C→T | A389V |
| LD6_0250_phi2 | Pf6* | MPAO1_RS24195 | CTXphi Zot-like toxin | SNP | synonymous | C→T | V394V |
| LD6_0250_phi2 | Pf6* | MPAO1_RS24195 | CTXphi Zot-like toxin | SNP | synonymous | G→C | A396A |
| LD6_0250_phi2 | Pf6* | MPAO1_RS24195 | CTXphi Zot-like toxin | SNP | nonsynonymous | T→C | V400A |
| LD6_0250_phi2 | Pf6* | MPAO1_RS24195 | CTXphi Zot-like toxin | SNP | synonymous | T→G | G404G |
| LD6_0250_phi2 | Pf6* | MPAO1_RS24195 | CTXphi Zot-like toxin | SNP | nonsynonymous | C→G | H423Q |
| LD6_0250_phi2 | Pf6* | ZDSLLIWT_CDS_0014 | hypothetical protein | SUB | | Δ2bp | N/A |
| LD6_0250_phi2 | Pf6* | ZDSLLIWT_CDS_0014 | hypothetical protein | SNP | nonsynonymous | T→A | S21T |
| LD6_0250_phi2 | Pf6* | ZDSLLIWT_CDS_0014/MPAO1_RS24190 | hypothetical protein/hypothetical protein | SNP | intergenic | T→C | N/A |
| LD6_0250_phi3 | Pf6 | MPAO1_RS28990/MPAO1_RS24230 | transcriptional regulator/hypothetical protein | SNP | intergenic | C→T | N/A |
| LD6_0250_phi3 | Pf6 | MPAO1_RS28990/MPAO1_RS24230 | transcriptional regulator/hypothetical protein | DEL | | Δ4bp | N/A |
| LD6_0250_phi3 | Pf6 | MPAO1_RS24215 | hypothetical protein | SNP | nonsynonymous | A→C | Q8P |
| LD6_0500_phi1 | Pf6 | MPAO1_RS28990/MPAO1_RS24230 | transcriptional regulator/hypothetical protein | SNP | intergenic | C→T | N/A |
| LD6_0500_phi1 | Pf6 | MPAO1_RS28990/MPAO1_RS24230 | transcriptional regulator/hypothetical protein | DEL | | Δ4bp | N/A |
| LD6_0500_phi1 | Pf6 | MPAO1_RS24215 | hypothetical protein | SNP | nonsynonymous | A→C | Q8P |
| LD6_0500_phi1 | Pf6 | MPAO1_RS24195 | CTXphi Zot-like toxin | SNP | nonsynonymous | C→T | R355W |
| LD6_0500_phi2 | Pf6 | MPAO1_RS28990/MPAO1_RS24230 | transcriptional regulator/hypothetical protein | SNP | intergenic | C→T | N/A |
| LD6_0500_phi2 | Pf6 | MPAO1_RS28990/MPAO1_RS24230 | transcriptional regulator/hypothetical protein | DEL | | Δ4bp | N/A |
| LD6_0500_phi2 | Pf6 | MPAO1_RS24215 | hypothetical protein | SNP | nonsynonymous | A→C | Q8P |
| LD6_0500_phi3 | Pf6 | MPAO1_RS28990/MPAO1_RS24230 | transcriptional regulator/hypothetical protein | SNP | intergenic | C→T | N/A |
| LD6_0500_phi3 | Pf6 | MPAO1_RS28990/MPAO1_RS24230 | transcriptional regulator/hypothetical protein | DEL | | Δ4bp | N/A |
| LD6_0500_phi3 | Pf6 | MPAO1_RS24215 | hypothetical protein | SNP | nonsynonymous | A→C | Q8P |
| LD8_0250_phi1 | Pf4 | [PA0716.1] | [PA0716.1] | DEL | | Δ141bp | N/A |

| | | | | | | | |
|---|---|---|---|---|---|---|---|
| LD8_0250_phi2 | Pf4 | [PA0716.1] | [PA0716.1] | DEL | | Δ141bp | N/A |
| LD8_0500_phi1 | Pf4 | [PA0716.1] | [PA0716.1] | DEL | | Δ141bp | N/A |
| LD8_0500_phi1 | Pf4 | PA0720 | single strand DNA binding protein | SNP | nonsynonymous | C→TRUE | P82S |
| LD8_0500_phi2 | Pf4 | [PA0716.1] | [PA0716.1] | DEL | | Δ141bp | N/A |
| LD8_0500_phi3 | Pf4 | [PA0716.1] | [PA0716.1] | DEL | | Δ141bp | N/A |
| LD8_0500_phi3 | Pf4 | PA0720 | single strand DNA binding protein | SNP | nonsynonymous | C→TRUE | P82S |

*indicates recombinant phage isolates.
