## Supplementary File 1 for "Mutations in filamentous bacteriophages spark eco-evolutionary feedbacks in *Pseudomonas aeruginosa*"

NCEE - Prophage Paper - Stats


### NCEE - Prophage Paper - Stats

###### Noah Houpt

#### 2025-08-21

### Overview

This RMarkdown file contains the code that generated the analyses
discussed in Houpt, Hernandez, and Turner (2026).

This document is organized by result section. These are sections in
the manuscript that each focus on one main figure. For example, the
first section below generates statistical models related to statements
in the first section of the results, which concerns main figure 1.

Paths to data files are based on the first author’s machine, so
please adjust accordingly when replicating analyses.

#### Loading Packages

```
library(tidyverse) #Version 1.3.2
```

```
## ── Attaching core tidyverse packages ──────────────────────── tidyverse 2.0.0 ──
## ✔ dplyr     1.2.1     ✔ readr     2.2.0
## ✔ forcats   1.0.1     ✔ stringr   1.6.0
## ✔ ggplot2   4.0.2     ✔ tibble    3.3.1
## ✔ lubridate 1.9.5     ✔ tidyr     1.3.2
## ✔ purrr     1.2.2     
## ── Conflicts ────────────────────────────────────────── tidyverse_conflicts() ──
## ✖ dplyr::filter() masks stats::filter()
## ✖ dplyr::lag()    masks stats::lag()
## ℹ Use the conflicted package (<http://conflicted.r-lib.org/>) to force all conflicts to become errors
```

```
library(lme4) #version 1.1-29
```

```
## Loading required package: Matrix
## 
## Attaching package: 'Matrix'
## 
## The following objects are masked from 'package:tidyr':
## 
##     expand, pack, unpack
```

```
library(lmerTest) #Version 3.1-3
```

```
## 
## Attaching package: 'lmerTest'
## 
## The following object is masked from 'package:lme4':
## 
##     lmer
## 
## The following object is masked from 'package:stats':
## 
##     step
```

```
library(emmeans) #Version 1.8.1-1
```

```
## Welcome to emmeans.
## Caution: You lose important information if you filter this package's results.
## See '? untidy'
```

```
library(car) #Version 3.1-3
```

```
## Loading required package: carData
## Registered S3 method overwritten by 'car':
##   method           from
##   na.action.merMod lme4
## 
## Attaching package: 'car'
## 
## The following object is masked from 'package:dplyr':
## 
##     recode
## 
## The following object is masked from 'package:purrr':
## 
##     some
```

```
library(gcplyr) #Version 1.5.2
```

```
## ## 
## ## gcplyr (Version 1.12.0, Build Date: 2025-07-28)
## ## See http://github.com/mikeblazanin/gcplyr for additional documentation
## ## Please cite software as:
## ##   Blazanin, Michael. gcplyr: an R package for microbial growth
## ##   curve data analysis. BMC Bioinformatics 25, 232 (2024).
## ##   https://doi.org/10.1186/s12859-024-05817-3
## ##
```

```
library(lubridate) #Version 1.8.0
library(vctrs) #Version 0.6.5
```

```
## 
## Attaching package: 'vctrs'
## 
## The following object is masked from 'package:dplyr':
## 
##     data_frame
## 
## The following object is masked from 'package:tibble':
## 
##     data_frame
```

```
library(nlme) #Version 3.1-157
```

```
## 
## Attaching package: 'nlme'
## 
## The following object is masked from 'package:lme4':
## 
##     lmList
## 
## The following object is masked from 'package:dplyr':
## 
##     collapse
```

#### Figure 1

The results section centered on Figure 1 involves the following
statistical models:

Model 1.1: End-of-day population density (CFU/mL) over generations
during experimental evolution.

Model 1.2: Area-under-the-curve of ancestral, evolved HD, and evolved
LD populations in unmodified media.

Model 1.3.xxx: For populations HD/LD1-8: area-under-the-curve of
endpoint and ancestral bacterial populations across filtrates. Separate
model for each population.

Note: In this section and throughout, I refer to “supernatant”, “st”,
“filtrate”, and “fil” interchangeably.

##### Model 1.1

###### Importing Data

```
dens_data <- read.csv("~/Documents/PhD Folder/Research/2023 Fall/evol_exp_dens_data.csv")
```

###### Calculating density in CFU/mL, averaging across replicates, and removing unecessary columns

```
dens_data$dens <- (dens_data$count/dens_data$vol)*10^dens_data$dil #Computing density in CFU/mL from raw data

dens_data <- dens_data %>% aggregate(dens~pop_id+strain+treat+day+vol, mean) #Averaging density measurements across replicates (populations plated in duplicate)

dens_data <- dens_data[,c(1:4,6)] #Removing unnecessary columns

dens_data <- dens_data %>% mutate(gen = case_when(treat=="ANC" ~ 0, treat=="HD" ~ day*6.643856, treat=="LD" ~ day*11.16742)) #Calculating generation times based on the number of days multiplied by the unique scaling factors across treatments (LD populations go through more generations per day due to their higher daily dilution factor)
```

###### Generating Model 1.1

Generating mixed-effects model with log10 population density (CFU/mL)
as the response variable, generation and treatment as main effects, and
population ID as a random effect.

Excluding population density measurements of ancestors from day 0
(day before first passage).

```
#Generating Model 1.1
mod1.1 <- lmer(data=dens_data[dens_data$treat != "ANC" & dens_data$gen < 1020,], log10(dens) ~ gen + treat + (1|pop_id))

#Summarizing Model 1.1 main effects
Anova(mod1.1, type="III")
```

```
## Analysis of Deviance Table (Type III Wald chisquare tests)
## 
## Response: log10(dens)
##                  Chisq Df Pr(>Chisq)    
## (Intercept) 2.3756e+05  1  < 2.2e-16 ***
## gen         6.6582e+01  1  3.356e-16 ***
## treat       2.2370e-01  1     0.6362    
## ---
## Signif. codes:  0 '***' 0.001 '**' 0.01 '*' 0.05 '.' 0.1 ' ' 1
```

```
#Generating model 1.1.1 in response to reviewer request for magnitude of density increase over experiment

#subsetting density data for starting and ending densities for each population
dens_data_sub <- dens_data[dens_data$gen < 12 & dens_data$treat != "ANC" | dens_data$gen < 1020 & dens_data$gen >980 & dens_data$treat != "ANC",]

#creating a new column called "start" for day 1 densities and "end" for densities at gen ~ 1000
dens_data_sub$stage <- if_else(dens_data_sub$gen < 20, "start", "end")

#generating model
mod1.1.1 <- lmer(data=dens_data_sub, log10(dens) ~ stage + treat + (1|pop_id))

#generating model output
summary(mod1.1.1)
```

```
## Linear mixed model fit by REML. t-tests use Satterthwaite's method [
## lmerModLmerTest]
## Formula: log10(dens) ~ stage + treat + (1 | pop_id)
##    Data: dens_data_sub
## 
## REML criterion at convergence: -73.2
## 
## Scaled residuals: 
##      Min       1Q   Median       3Q      Max 
## -2.16876 -0.53702 -0.08709  0.46365  2.31633 
## 
## Random effects:
##  Groups   Name        Variance  Std.Dev.
##  pop_id   (Intercept) 0.0008171 0.02858 
##  Residual             0.0029060 0.05391 
## Number of obs: 32, groups:  pop_id, 16
## 
## Fixed effects:
##             Estimate Std. Error       df t value Pr(>|t|)    
## (Intercept)  9.97848    0.01935 22.26626 515.582  < 2e-16 ***
## stagestart  -0.18123    0.01906 15.00000  -9.509 9.65e-08 ***
## treatLD      0.02089    0.02382 14.00000   0.877    0.395    
## ---
## Signif. codes:  0 '***' 0.001 '**' 0.01 '*' 0.05 '.' 0.1 ' ' 1
## 
## Correlation of Fixed Effects:
##            (Intr) stgstr
## stagestart -0.492       
## treatLD    -0.615  0.000
```

###### Model 1.1 Summary

1. We detected no overall difference in end-of-day population
   density across treatments.
2. End-of-day population density increased, on average, over
   generations during the evolution experiment.
3. log10 population density increased by an average of 0.000209 per
   generation (standard error = 0.000026).
4. log10 population density increased by an average of 0.18 over the
   experiment (se = 0.019)

##### Model 1.2

###### Importing growth curve data

```
h1_l1_st_r1 <- read_wides(files = "~/Documents/PhD Folder/Research/2025 Spring/ncee_gcs/ncee_gc_data/2025.01.16_h1-l1-st_r1/nh_2025.01.16_h1-l1-st_r1.csv", startrow=34, endrow=173) #Importing single plate of growth curve data (repeated below)

h1_l1_st_r1 <- trans_wide_to_tidy(wides=h1_l1_st_r1[,-c(1,3)], id_cols=c("Time")) #Converting imported growth curve to tidy format (repeated below)

h1_l1_st_r2 <- read_wides(files = "~/Documents/PhD Folder/Research/2025 Spring/ncee_gcs/ncee_gc_data/2025.01.28_h1-l1-st_r2/nh_2025.01.28_h1-l1-st_r2.csv", startrow=27, endrow=166)

h1_l1_st_r2 <- trans_wide_to_tidy(wides=h1_l1_st_r2[,-c(1,2,4)], id_cols=c("Time"))

h2_l2_st_r1 <- read_wides(files = "~/Documents/PhD Folder/Research/2025 Spring/ncee_gcs/ncee_gc_data/2025.01.16_h2-l2-st_r1/nh_2025.01.16_h2-l2-st_r1.csv", startrow=27, endrow=166)

h2_l2_st_r1 <- trans_wide_to_tidy(wides=h2_l2_st_r1[,-c(1,2,4)], id_cols=c("Time"))

h2_l2_st_r2 <- read_wides(files = "~/Documents/PhD Folder/Research/2025 Spring/ncee_gcs/ncee_gc_data/2025.01.28_h2-l2-st_r2/nh_2025.01.28_h2-l2-st_r2.csv", startrow=34, endrow=173)

h2_l2_st_r2 <- trans_wide_to_tidy(wides=h2_l2_st_r2[,-c(1,3)], id_cols=c("Time"))

h3_l3_st_r1 <- read_wides(files = "~/Documents/PhD Folder/Research/2025 Spring/ncee_gcs/ncee_gc_data/2025.01.17_h3-l3-st_r1/nh_2025.01.17_h3-l3-st_r1.csv", startrow=34, endrow=173)

h3_l3_st_r1 <- trans_wide_to_tidy(wides=h3_l3_st_r1[,-c(1,3)], id_cols=c("Time"))

h3_l3_st_r2 <- read_wides(files = "~/Documents/PhD Folder/Research/2025 Spring/ncee_gcs/ncee_gc_data/2025.01.30_h3-l3-st_r2/nh_2025.01.30_h3-l3-st_r2.csv", startrow=27, endrow=166)

h3_l3_st_r2 <- trans_wide_to_tidy(wides=h3_l3_st_r2[,-c(1,2,4)], id_cols=c("Time"))

h4_l4_st_r1 <- read_wides(files = "~/Documents/PhD Folder/Research/2025 Spring/ncee_gcs/ncee_gc_data/2025.01.17_h4-l4-st_r1/nh_2025.01.17_h4-l4-st_r1.csv", startrow=27, endrow=166)

h4_l4_st_r1 <- trans_wide_to_tidy(wides=h4_l4_st_r1[,-c(1,2,4)], id_cols=c("Time"))

h4_l4_st_r2 <- read_wides(files = "~/Documents/PhD Folder/Research/2025 Spring/ncee_gcs/ncee_gc_data/2025.01.30_h4-l4-st_r2/nh_2025.01.30_h4-l4-st_r2.csv", startrow=34, endrow=173)

h4_l4_st_r2 <- trans_wide_to_tidy(wides=h4_l4_st_r2[,-c(1,3)], id_cols=c("Time"))

h5_l5_st_r1 <- read_wides(files = "~/Documents/PhD Folder/Research/2025 Spring/ncee_gcs/ncee_gc_data/2025.01.20_h5-l5-st_r1/nh_2025.01.20_h5-l5-st_r1.csv", startrow=34, endrow=173)

h5_l5_st_r1 <- trans_wide_to_tidy(wides=h5_l5_st_r1[,-c(1,3)], id_cols=c("Time"))

h5_l5_st_r2 <- read_wides(files = "~/Documents/PhD Folder/Research/2025 Spring/ncee_gcs/ncee_gc_data/2025.02.05_h5-l5-st_r2/nh_2025.02.05_h5-l5-st_r2.csv", startrow=27, endrow=166)

h5_l5_st_r2 <- trans_wide_to_tidy(wides=h5_l5_st_r2[,-c(1,2,4)], id_cols=c("Time"))

h6_l6_st_r1 <- read_wides(files = "~/Documents/PhD Folder/Research/2025 Spring/ncee_gcs/ncee_gc_data/2025.01.20_h6-l6-st_r1/nh_2025.01.20_h6-l6-st_r1.csv", startrow=27, endrow=166)

h6_l6_st_r1 <- trans_wide_to_tidy(wides=h6_l6_st_r1[,-c(1,2,4)], id_cols=c("Time"))

h6_l6_st_r2 <- read_wides(files = "~/Documents/PhD Folder/Research/2025 Spring/ncee_gcs/ncee_gc_data/2025.02.05_h6-l6-st_r2/nh_2025.02.05_h6-l6-st_r2.csv", startrow=34, endrow=173)

h6_l6_st_r2 <- trans_wide_to_tidy(wides=h6_l6_st_r2[,-c(1,3)], id_cols=c("Time"))

h7_l7_st_r1 <- read_wides(files = "~/Documents/PhD Folder/Research/2025 Spring/ncee_gcs/ncee_gc_data/2025.01.21_h7-l7-st_r1/nh_2025.01.21_h7-l7-st_r1.csv", startrow=34, endrow=173)

h7_l7_st_r1 <- trans_wide_to_tidy(wides=h7_l7_st_r1[,-c(1,3)], id_cols=c("Time"))

h7_l7_st_r2 <- read_wides(files = "~/Documents/PhD Folder/Research/2025 Spring/ncee_gcs/ncee_gc_data/2025.02.07_h7-l7-st_r2/nh_2025.02.07_h7-l7-st_r2.csv", startrow=27, endrow=166)

h7_l7_st_r2 <- trans_wide_to_tidy(wides=h7_l7_st_r2[,-c(1,2,4)], id_cols=c("Time"))

h8_l8_st_r1 <- read_wides(files = "~/Documents/PhD Folder/Research/2025 Spring/ncee_gcs/ncee_gc_data/2025.01.21_h8-l8-st_r1/nh_2025.01.21_h8-l8-st_r1.csv", startrow=27, endrow=166)

h8_l8_st_r1 <- trans_wide_to_tidy(wides=h8_l8_st_r1[,-c(1,2,4)], id_cols=c("Time"))

h8_l8_st_r2 <- read_wides(files = "~/Documents/PhD Folder/Research/2025 Spring/ncee_gcs/ncee_gc_data/2025.02.07_h8-l8-st_r2/nh_2025.02.07_h8-l8-st_r2.csv", startrow=34, endrow=173)

h8_l8_st_r2 <- trans_wide_to_tidy(wides=h8_l8_st_r2[,-c(1,3)], id_cols=c("Time"))
```

###### Importing growth curve design files

```
h1_l1_st_r1_design<-import_blockdesigns(files=c("~/Documents/PhD Folder/Research/2025 Spring/ncee_gcs/ncee_gc_data/2025.01.16_h1-l1-st_r1/h1-l1-st_r1_st.csv", "~/Documents/PhD Folder/Research/2025 Spring/ncee_gcs/ncee_gc_data/2025.01.16_h1-l1-st_r1/h1-l1-st_r1_pop-id.csv", "~/Documents/PhD Folder/Research/2025 Spring/ncee_gcs/ncee_gc_data/2025.01.16_h1-l1-st_r1/h1-l1-st_r1_machine.csv", "~/Documents/PhD Folder/Research/2025 Spring/ncee_gcs/ncee_gc_data/2025.01.16_h1-l1-st_r1/h1-l1-st_r1_evol-treat.csv", "~/Documents/PhD Folder/Research/2025 Spring/ncee_gcs/ncee_gc_data/2025.01.16_h1-l1-st_r1/h1-l1-st_r1_date.csv", "~/Documents/PhD Folder/Research/2025 Spring/ncee_gcs/ncee_gc_data/2025.01.16_h1-l1-st_r1/h1-l1-st_r1_bac-id.csv", "~/Documents/PhD Folder/Research/2025 Spring/ncee_gcs/ncee_gc_data/2025.01.16_h1-l1-st_r1/h1-l1-st_r1_bac-gen.csv"), block_names = c("st", "pop_id", "machine", "evol_treat", "date", "bac_id", "bac_gen")) #Importing set of design files for one plate of growth curves (repeated below)
```

```
## Inferred 'into' column names as: st, pop_id, machine, evol_treat, date, bac_id, bac_gen
```

```
h1_l1_st_r2_design<-import_blockdesigns(files=c("~/Documents/PhD Folder/Research/2025 Spring/ncee_gcs/ncee_gc_data/2025.01.28_h1-l1-st_r2/h1-l1-st_r2_st.csv", "~/Documents/PhD Folder/Research/2025 Spring/ncee_gcs/ncee_gc_data/2025.01.28_h1-l1-st_r2/h1-l1-st_r2_pop-id.csv", "~/Documents/PhD Folder/Research/2025 Spring/ncee_gcs/ncee_gc_data/2025.01.28_h1-l1-st_r2/h1-l1-st_r2_machine.csv", "~/Documents/PhD Folder/Research/2025 Spring/ncee_gcs/ncee_gc_data/2025.01.28_h1-l1-st_r2/h1-l1-st_r2_evol-treat.csv", "~/Documents/PhD Folder/Research/2025 Spring/ncee_gcs/ncee_gc_data/2025.01.28_h1-l1-st_r2/h1-l1-st_r2_date.csv", "~/Documents/PhD Folder/Research/2025 Spring/ncee_gcs/ncee_gc_data/2025.01.28_h1-l1-st_r2/h1-l1-st_r2_bac-id.csv", "~/Documents/PhD Folder/Research/2025 Spring/ncee_gcs/ncee_gc_data/2025.01.28_h1-l1-st_r2/h1-l1-st_r2_bac-gen.csv"), block_names = c("st", "pop_id", "machine", "evol_treat", "date", "bac_id", "bac_gen"))
```

```
## Inferred 'into' column names as: st, pop_id, machine, evol_treat, date, bac_id, bac_gen
```

```
h2_l2_st_r1_design<-import_blockdesigns(files=c("~/Documents/PhD Folder/Research/2025 Spring/ncee_gcs/ncee_gc_data/2025.01.16_h2-l2-st_r1/h2-l2-st_r1_st.csv", "~/Documents/PhD Folder/Research/2025 Spring/ncee_gcs/ncee_gc_data/2025.01.16_h2-l2-st_r1/h2-l2-st_r1_pop-id.csv", "~/Documents/PhD Folder/Research/2025 Spring/ncee_gcs/ncee_gc_data/2025.01.16_h2-l2-st_r1/h2-l2-st_r1_machine.csv", "~/Documents/PhD Folder/Research/2025 Spring/ncee_gcs/ncee_gc_data/2025.01.16_h2-l2-st_r1/h2-l2-st_r1_evol-treat.csv", "~/Documents/PhD Folder/Research/2025 Spring/ncee_gcs/ncee_gc_data/2025.01.16_h2-l2-st_r1/h2-l2-st_r1_date.csv", "~/Documents/PhD Folder/Research/2025 Spring/ncee_gcs/ncee_gc_data/2025.01.16_h2-l2-st_r1/h2-l2-st_r1_bac-id.csv", "~/Documents/PhD Folder/Research/2025 Spring/ncee_gcs/ncee_gc_data/2025.01.16_h2-l2-st_r1/h2-l2-st_r1_bac-gen.csv"), block_names = c("st", "pop_id", "machine", "evol_treat", "date", "bac_id", "bac_gen"))
```

```
## Inferred 'into' column names as: st, pop_id, machine, evol_treat, date, bac_id, bac_gen
```

```
h2_l2_st_r2_design<-import_blockdesigns(files=c("~/Documents/PhD Folder/Research/2025 Spring/ncee_gcs/ncee_gc_data/2025.01.28_h2-l2-st_r2/h2-l2-st_r2_st.csv", "~/Documents/PhD Folder/Research/2025 Spring/ncee_gcs/ncee_gc_data/2025.01.28_h2-l2-st_r2/h2-l2-st_r2_pop-id.csv", "~/Documents/PhD Folder/Research/2025 Spring/ncee_gcs/ncee_gc_data/2025.01.28_h2-l2-st_r2/h2-l2-st_r2_machine.csv", "~/Documents/PhD Folder/Research/2025 Spring/ncee_gcs/ncee_gc_data/2025.01.28_h2-l2-st_r2/h2-l2-st_r2_evol-treat.csv", "~/Documents/PhD Folder/Research/2025 Spring/ncee_gcs/ncee_gc_data/2025.01.28_h2-l2-st_r2/h2-l2-st_r2_date.csv", "~/Documents/PhD Folder/Research/2025 Spring/ncee_gcs/ncee_gc_data/2025.01.28_h2-l2-st_r2/h2-l2-st_r2_bac-id.csv", "~/Documents/PhD Folder/Research/2025 Spring/ncee_gcs/ncee_gc_data/2025.01.28_h2-l2-st_r2/h2-l2-st_r2_bac-gen.csv"), block_names = c("st", "pop_id", "machine", "evol_treat", "date", "bac_id", "bac_gen"))
```

```
## Inferred 'into' column names as: st, pop_id, machine, evol_treat, date, bac_id, bac_gen
```

```
h3_l3_st_r1_design<-import_blockdesigns(files=c("~/Documents/PhD Folder/Research/2025 Spring/ncee_gcs/ncee_gc_data/2025.01.17_h3-l3-st_r1/h3-l3-st_r1_st.csv", "~/Documents/PhD Folder/Research/2025 Spring/ncee_gcs/ncee_gc_data/2025.01.17_h3-l3-st_r1/h3-l3-st_r1_pop-id.csv", "~/Documents/PhD Folder/Research/2025 Spring/ncee_gcs/ncee_gc_data/2025.01.17_h3-l3-st_r1/h3-l3-st_r1_machine.csv", "~/Documents/PhD Folder/Research/2025 Spring/ncee_gcs/ncee_gc_data/2025.01.17_h3-l3-st_r1/h3-l3-st_r1_evol-treat.csv", "~/Documents/PhD Folder/Research/2025 Spring/ncee_gcs/ncee_gc_data/2025.01.17_h3-l3-st_r1/h3-l3-st_r1_date.csv", "~/Documents/PhD Folder/Research/2025 Spring/ncee_gcs/ncee_gc_data/2025.01.17_h3-l3-st_r1/h3-l3-st_r1_bac-id.csv", "~/Documents/PhD Folder/Research/2025 Spring/ncee_gcs/ncee_gc_data/2025.01.17_h3-l3-st_r1/h3-l3-st_r1_bac-gen.csv"), block_names = c("st", "pop_id", "machine", "evol_treat", "date", "bac_id", "bac_gen"))
```

```
## Inferred 'into' column names as: st, pop_id, machine, evol_treat, date, bac_id, bac_gen
```

```
h3_l3_st_r2_design<-import_blockdesigns(files=c("~/Documents/PhD Folder/Research/2025 Spring/ncee_gcs/ncee_gc_data/2025.01.30_h3-l3-st_r2/h3-l3-st_r2_st.csv", "~/Documents/PhD Folder/Research/2025 Spring/ncee_gcs/ncee_gc_data/2025.01.30_h3-l3-st_r2/h3-l3-st_r2_pop-id.csv", "~/Documents/PhD Folder/Research/2025 Spring/ncee_gcs/ncee_gc_data/2025.01.30_h3-l3-st_r2/h3-l3-st_r2_machine.csv", "~/Documents/PhD Folder/Research/2025 Spring/ncee_gcs/ncee_gc_data/2025.01.30_h3-l3-st_r2/h3-l3-st_r2_evol-treat.csv", "~/Documents/PhD Folder/Research/2025 Spring/ncee_gcs/ncee_gc_data/2025.01.30_h3-l3-st_r2/h3-l3-st_r2_date.csv", "~/Documents/PhD Folder/Research/2025 Spring/ncee_gcs/ncee_gc_data/2025.01.30_h3-l3-st_r2/h3-l3-st_r2_bac-id.csv", "~/Documents/PhD Folder/Research/2025 Spring/ncee_gcs/ncee_gc_data/2025.01.30_h3-l3-st_r2/h3-l3-st_r2_bac-gen.csv"), block_names = c("st", "pop_id", "machine", "evol_treat", "date", "bac_id", "bac_gen"))
```

```
## Inferred 'into' column names as: st, pop_id, machine, evol_treat, date, bac_id, bac_gen
```

```
h4_l4_st_r1_design<-import_blockdesigns(files=c("~/Documents/PhD Folder/Research/2025 Spring/ncee_gcs/ncee_gc_data/2025.01.17_h4-l4-st_r1/h4-l4-st_r1_st.csv", "~/Documents/PhD Folder/Research/2025 Spring/ncee_gcs/ncee_gc_data/2025.01.17_h4-l4-st_r1/h4-l4-st_r1_pop-id.csv", "~/Documents/PhD Folder/Research/2025 Spring/ncee_gcs/ncee_gc_data/2025.01.17_h4-l4-st_r1/h4-l4-st_r1_machine.csv", "~/Documents/PhD Folder/Research/2025 Spring/ncee_gcs/ncee_gc_data/2025.01.17_h4-l4-st_r1/h4-l4-st_r1_evol-treat.csv", "~/Documents/PhD Folder/Research/2025 Spring/ncee_gcs/ncee_gc_data/2025.01.17_h4-l4-st_r1/h4-l4-st_r1_date.csv", "~/Documents/PhD Folder/Research/2025 Spring/ncee_gcs/ncee_gc_data/2025.01.17_h4-l4-st_r1/h4-l4-st_r1_bac-id.csv", "~/Documents/PhD Folder/Research/2025 Spring/ncee_gcs/ncee_gc_data/2025.01.17_h4-l4-st_r1/h4-l4-st_r1_bac-gen.csv"), block_names = c("st", "pop_id", "machine", "evol_treat", "date", "bac_id", "bac_gen"))
```

```
## Inferred 'into' column names as: st, pop_id, machine, evol_treat, date, bac_id, bac_gen
```

```
h4_l4_st_r2_design<-import_blockdesigns(files=c("~/Documents/PhD Folder/Research/2025 Spring/ncee_gcs/ncee_gc_data/2025.01.30_h4-l4-st_r2/h4-l4-st_r2_st.csv", "~/Documents/PhD Folder/Research/2025 Spring/ncee_gcs/ncee_gc_data/2025.01.30_h4-l4-st_r2/h4-l4-st_r2_pop-id.csv", "~/Documents/PhD Folder/Research/2025 Spring/ncee_gcs/ncee_gc_data/2025.01.30_h4-l4-st_r2/h4-l4-st_r2_machine.csv", "~/Documents/PhD Folder/Research/2025 Spring/ncee_gcs/ncee_gc_data/2025.01.30_h4-l4-st_r2/h4-l4-st_r2_evol-treat.csv", "~/Documents/PhD Folder/Research/2025 Spring/ncee_gcs/ncee_gc_data/2025.01.30_h4-l4-st_r2/h4-l4-st_r2_date.csv", "~/Documents/PhD Folder/Research/2025 Spring/ncee_gcs/ncee_gc_data/2025.01.30_h4-l4-st_r2/h4-l4-st_r2_bac-id.csv", "~/Documents/PhD Folder/Research/2025 Spring/ncee_gcs/ncee_gc_data/2025.01.30_h4-l4-st_r2/h4-l4-st_r2_bac-gen.csv"), block_names = c("st", "pop_id", "machine", "evol_treat", "date", "bac_id", "bac_gen"))
```

```
## Inferred 'into' column names as: st, pop_id, machine, evol_treat, date, bac_id, bac_gen
```

```
h5_l5_st_r1_design<-import_blockdesigns(files=c("~/Documents/PhD Folder/Research/2025 Spring/ncee_gcs/ncee_gc_data/2025.01.20_h5-l5-st_r1/h5-l5-st_r1_st.csv", "~/Documents/PhD Folder/Research/2025 Spring/ncee_gcs/ncee_gc_data/2025.01.20_h5-l5-st_r1/h5-l5-st_r1_pop-id.csv", "~/Documents/PhD Folder/Research/2025 Spring/ncee_gcs/ncee_gc_data/2025.01.20_h5-l5-st_r1/h5-l5-st_r1_machine.csv", "~/Documents/PhD Folder/Research/2025 Spring/ncee_gcs/ncee_gc_data/2025.01.20_h5-l5-st_r1/h5-l5-st_r1_evol-treat.csv", "~/Documents/PhD Folder/Research/2025 Spring/ncee_gcs/ncee_gc_data/2025.01.20_h5-l5-st_r1/h5-l5-st_r1_date.csv", "~/Documents/PhD Folder/Research/2025 Spring/ncee_gcs/ncee_gc_data/2025.01.20_h5-l5-st_r1/h5-l5-st_r1_bac-id.csv", "~/Documents/PhD Folder/Research/2025 Spring/ncee_gcs/ncee_gc_data/2025.01.20_h5-l5-st_r1/h5-l5-st_r1_bac-gen.csv"), block_names = c("st", "pop_id", "machine", "evol_treat", "date", "bac_id", "bac_gen"))
```

```
## Inferred 'into' column names as: st, pop_id, machine, evol_treat, date, bac_id, bac_gen
```

```
h5_l5_st_r2_design<-import_blockdesigns(files=c("~/Documents/PhD Folder/Research/2025 Spring/ncee_gcs/ncee_gc_data/2025.02.05_h5-l5-st_r2/h5-l5-st_r2_st.csv", "~/Documents/PhD Folder/Research/2025 Spring/ncee_gcs/ncee_gc_data/2025.02.05_h5-l5-st_r2/h5-l5-st_r2_pop-id.csv", "~/Documents/PhD Folder/Research/2025 Spring/ncee_gcs/ncee_gc_data/2025.02.05_h5-l5-st_r2/h5-l5-st_r2_machine.csv", "~/Documents/PhD Folder/Research/2025 Spring/ncee_gcs/ncee_gc_data/2025.02.05_h5-l5-st_r2/h5-l5-st_r2_evol-treat.csv", "~/Documents/PhD Folder/Research/2025 Spring/ncee_gcs/ncee_gc_data/2025.02.05_h5-l5-st_r2/h5-l5-st_r2_date.csv", "~/Documents/PhD Folder/Research/2025 Spring/ncee_gcs/ncee_gc_data/2025.02.05_h5-l5-st_r2/h5-l5-st_r2_bac-id.csv", "~/Documents/PhD Folder/Research/2025 Spring/ncee_gcs/ncee_gc_data/2025.02.05_h5-l5-st_r2/h5-l5-st_r2_bac-gen.csv"), block_names = c("st", "pop_id", "machine", "evol_treat", "date", "bac_id", "bac_gen"))
```

```
## Inferred 'into' column names as: st, pop_id, machine, evol_treat, date, bac_id, bac_gen
```

```
h6_l6_st_r1_design<-import_blockdesigns(files=c("~/Documents/PhD Folder/Research/2025 Spring/ncee_gcs/ncee_gc_data/2025.01.20_h6-l6-st_r1/h6-l6-st_r1_st.csv", "~/Documents/PhD Folder/Research/2025 Spring/ncee_gcs/ncee_gc_data/2025.01.20_h6-l6-st_r1/h6-l6-st_r1_pop-id.csv", "~/Documents/PhD Folder/Research/2025 Spring/ncee_gcs/ncee_gc_data/2025.01.20_h6-l6-st_r1/h6-l6-st_r1_machine.csv", "~/Documents/PhD Folder/Research/2025 Spring/ncee_gcs/ncee_gc_data/2025.01.20_h6-l6-st_r1/h6-l6-st_r1_evol-treat.csv", "~/Documents/PhD Folder/Research/2025 Spring/ncee_gcs/ncee_gc_data/2025.01.20_h6-l6-st_r1/h6-l6-st_r1_date.csv", "~/Documents/PhD Folder/Research/2025 Spring/ncee_gcs/ncee_gc_data/2025.01.20_h6-l6-st_r1/h6-l6-st_r1_bac-id.csv", "~/Documents/PhD Folder/Research/2025 Spring/ncee_gcs/ncee_gc_data/2025.01.20_h6-l6-st_r1/h6-l6-st_r1_bac-gen.csv"), block_names = c("st", "pop_id", "machine", "evol_treat", "date", "bac_id", "bac_gen"))
```

```
## Inferred 'into' column names as: st, pop_id, machine, evol_treat, date, bac_id, bac_gen
```

```
h6_l6_st_r2_design<-import_blockdesigns(files=c("~/Documents/PhD Folder/Research/2025 Spring/ncee_gcs/ncee_gc_data/2025.02.05_h6-l6-st_r2/h6-l6-st_r2_st.csv", "~/Documents/PhD Folder/Research/2025 Spring/ncee_gcs/ncee_gc_data/2025.02.05_h6-l6-st_r2/h6-l6-st_r2_pop-id.csv", "~/Documents/PhD Folder/Research/2025 Spring/ncee_gcs/ncee_gc_data/2025.02.05_h6-l6-st_r2/h6-l6-st_r2_machine.csv", "~/Documents/PhD Folder/Research/2025 Spring/ncee_gcs/ncee_gc_data/2025.02.05_h6-l6-st_r2/h6-l6-st_r2_evol-treat.csv", "~/Documents/PhD Folder/Research/2025 Spring/ncee_gcs/ncee_gc_data/2025.02.05_h6-l6-st_r2/h6-l6-st_r2_date.csv", "~/Documents/PhD Folder/Research/2025 Spring/ncee_gcs/ncee_gc_data/2025.02.05_h6-l6-st_r2/h6-l6-st_r2_bac-id.csv", "~/Documents/PhD Folder/Research/2025 Spring/ncee_gcs/ncee_gc_data/2025.02.05_h6-l6-st_r2/h6-l6-st_r2_bac-gen.csv"), block_names = c("st", "pop_id", "machine", "evol_treat", "date", "bac_id", "bac_gen"))
```

```
## Inferred 'into' column names as: st, pop_id, machine, evol_treat, date, bac_id, bac_gen
```

```
h7_l7_st_r1_design<-import_blockdesigns(files=c("~/Documents/PhD Folder/Research/2025 Spring/ncee_gcs/ncee_gc_data/2025.01.21_h7-l7-st_r1/h7-l7-st_r1_st.csv", "~/Documents/PhD Folder/Research/2025 Spring/ncee_gcs/ncee_gc_data/2025.01.21_h7-l7-st_r1/h7-l7-st_r1_pop-id.csv", "~/Documents/PhD Folder/Research/2025 Spring/ncee_gcs/ncee_gc_data/2025.01.21_h7-l7-st_r1/h7-l7-st_r1_machine.csv", "~/Documents/PhD Folder/Research/2025 Spring/ncee_gcs/ncee_gc_data/2025.01.21_h7-l7-st_r1/h7-l7-st_r1_evol-treat.csv", "~/Documents/PhD Folder/Research/2025 Spring/ncee_gcs/ncee_gc_data/2025.01.21_h7-l7-st_r1/h7-l7-st_r1_date.csv", "~/Documents/PhD Folder/Research/2025 Spring/ncee_gcs/ncee_gc_data/2025.01.21_h7-l7-st_r1/h7-l7-st_r1_bac-id.csv", "~/Documents/PhD Folder/Research/2025 Spring/ncee_gcs/ncee_gc_data/2025.01.21_h7-l7-st_r1/h7-l7-st_r1_bac-gen.csv"), block_names = c("st", "pop_id", "machine", "evol_treat", "date", "bac_id", "bac_gen"))
```

```
## Inferred 'into' column names as: st, pop_id, machine, evol_treat, date, bac_id, bac_gen
```

```
h7_l7_st_r2_design<-import_blockdesigns(files=c("~/Documents/PhD Folder/Research/2025 Spring/ncee_gcs/ncee_gc_data/2025.02.07_h7-l7-st_r2/h7-l7-st_r2_st.csv", "~/Documents/PhD Folder/Research/2025 Spring/ncee_gcs/ncee_gc_data/2025.02.07_h7-l7-st_r2/h7-l7-st_r2_pop-id.csv", "~/Documents/PhD Folder/Research/2025 Spring/ncee_gcs/ncee_gc_data/2025.02.07_h7-l7-st_r2/h7-l7-st_r2_machine.csv", "~/Documents/PhD Folder/Research/2025 Spring/ncee_gcs/ncee_gc_data/2025.02.07_h7-l7-st_r2/h7-l7-st_r2_evol-treat.csv", "~/Documents/PhD Folder/Research/2025 Spring/ncee_gcs/ncee_gc_data/2025.02.07_h7-l7-st_r2/h7-l7-st_r2_date.csv", "~/Documents/PhD Folder/Research/2025 Spring/ncee_gcs/ncee_gc_data/2025.02.07_h7-l7-st_r2/h7-l7-st_r2_bac-id.csv", "~/Documents/PhD Folder/Research/2025 Spring/ncee_gcs/ncee_gc_data/2025.02.07_h7-l7-st_r2/h7-l7-st_r2_bac-gen.csv"), block_names = c("st", "pop_id", "machine", "evol_treat", "date", "bac_id", "bac_gen"))
```

```
## Inferred 'into' column names as: st, pop_id, machine, evol_treat, date, bac_id, bac_gen
```

```
h8_l8_st_r1_design<-import_blockdesigns(files=c("~/Documents/PhD Folder/Research/2025 Spring/ncee_gcs/ncee_gc_data/2025.01.21_h8-l8-st_r1/h8-l8-st_r1_st.csv", "~/Documents/PhD Folder/Research/2025 Spring/ncee_gcs/ncee_gc_data/2025.01.21_h8-l8-st_r1/h8-l8-st_r1_pop-id.csv", "~/Documents/PhD Folder/Research/2025 Spring/ncee_gcs/ncee_gc_data/2025.01.21_h8-l8-st_r1/h8-l8-st_r1_machine.csv", "~/Documents/PhD Folder/Research/2025 Spring/ncee_gcs/ncee_gc_data/2025.01.21_h8-l8-st_r1/h8-l8-st_r1_evol-treat.csv", "~/Documents/PhD Folder/Research/2025 Spring/ncee_gcs/ncee_gc_data/2025.01.21_h8-l8-st_r1/h8-l8-st_r1_date.csv", "~/Documents/PhD Folder/Research/2025 Spring/ncee_gcs/ncee_gc_data/2025.01.21_h8-l8-st_r1/h8-l8-st_r1_bac-id.csv", "~/Documents/PhD Folder/Research/2025 Spring/ncee_gcs/ncee_gc_data/2025.01.21_h8-l8-st_r1/h8-l8-st_r1_bac-gen.csv"), block_names = c("st", "pop_id", "machine", "evol_treat", "date", "bac_id", "bac_gen"))
```

```
## Inferred 'into' column names as: st, pop_id, machine, evol_treat, date, bac_id, bac_gen
```

```
h8_l8_st_r2_design<-import_blockdesigns(files=c("~/Documents/PhD Folder/Research/2025 Spring/ncee_gcs/ncee_gc_data/2025.02.07_h8-l8-st_r2/h8-l8-st_r2_st.csv", "~/Documents/PhD Folder/Research/2025 Spring/ncee_gcs/ncee_gc_data/2025.02.07_h8-l8-st_r2/h8-l8-st_r2_pop-id.csv", "~/Documents/PhD Folder/Research/2025 Spring/ncee_gcs/ncee_gc_data/2025.02.07_h8-l8-st_r2/h8-l8-st_r2_machine.csv", "~/Documents/PhD Folder/Research/2025 Spring/ncee_gcs/ncee_gc_data/2025.02.07_h8-l8-st_r2/h8-l8-st_r2_evol-treat.csv", "~/Documents/PhD Folder/Research/2025 Spring/ncee_gcs/ncee_gc_data/2025.02.07_h8-l8-st_r2/h8-l8-st_r2_date.csv", "~/Documents/PhD Folder/Research/2025 Spring/ncee_gcs/ncee_gc_data/2025.02.07_h8-l8-st_r2/h8-l8-st_r2_bac-id.csv", "~/Documents/PhD Folder/Research/2025 Spring/ncee_gcs/ncee_gc_data/2025.02.07_h8-l8-st_r2/h8-l8-st_r2_bac-gen.csv"), block_names = c("st", "pop_id", "machine", "evol_treat", "date", "bac_id", "bac_gen"))
```

```
## Inferred 'into' column names as: st, pop_id, machine, evol_treat, date, bac_id, bac_gen
```

###### Merging raw growth curve data with design files

```
h1_l1_st_r1_merged <- merge_dfs(h1_l1_st_r1, h1_l1_st_r1_design) #merges one plate of growth curves with dataframe of design elements for those curves (repeated below)
```

```
## Joining with `by = join_by(Well)`
```

```
h1_l1_st_r2_merged <- merge_dfs(h1_l1_st_r2, h1_l1_st_r2_design)
```

```
## Joining with `by = join_by(Well)`
```

```
h2_l2_st_r1_merged <- merge_dfs(h2_l2_st_r1, h2_l2_st_r1_design)
```

```
## Joining with `by = join_by(Well)`
```

```
h2_l2_st_r2_merged <- merge_dfs(h2_l2_st_r2, h2_l2_st_r2_design)
```

```
## Joining with `by = join_by(Well)`
```

```
h3_l3_st_r1_merged <- merge_dfs(h3_l3_st_r1, h3_l3_st_r1_design)
```

```
## Joining with `by = join_by(Well)`
```

```
h3_l3_st_r2_merged <- merge_dfs(h3_l3_st_r2, h3_l3_st_r2_design)
```

```
## Joining with `by = join_by(Well)`
```

```
h4_l4_st_r1_merged <- merge_dfs(h4_l4_st_r1, h4_l4_st_r1_design)
```

```
## Joining with `by = join_by(Well)`
```

```
h4_l4_st_r2_merged <- merge_dfs(h4_l4_st_r2, h4_l4_st_r2_design)
```

```
## Joining with `by = join_by(Well)`
```

```
h5_l5_st_r1_merged <- merge_dfs(h5_l5_st_r1, h5_l5_st_r1_design)
```

```
## Joining with `by = join_by(Well)`
```

```
h5_l5_st_r2_merged <- merge_dfs(h5_l5_st_r2, h5_l5_st_r2_design)
```

```
## Joining with `by = join_by(Well)`
```

```
h6_l6_st_r1_merged <- merge_dfs(h6_l6_st_r1, h6_l6_st_r1_design)
```

```
## Joining with `by = join_by(Well)`
```

```
h6_l6_st_r2_merged <- merge_dfs(h6_l6_st_r2, h6_l6_st_r2_design)
```

```
## Joining with `by = join_by(Well)`
```

```
h7_l7_st_r1_merged <- merge_dfs(h7_l7_st_r1, h7_l7_st_r1_design)
```

```
## Joining with `by = join_by(Well)`
```

```
h7_l7_st_r2_merged <- merge_dfs(h7_l7_st_r2, h7_l7_st_r2_design)
```

```
## Joining with `by = join_by(Well)`
```

```
h8_l8_st_r1_merged <- merge_dfs(h8_l8_st_r1, h8_l8_st_r1_design)
```

```
## Joining with `by = join_by(Well)`
```

```
h8_l8_st_r2_merged <- merge_dfs(h8_l8_st_r2, h8_l8_st_r2_design)
```

```
## Joining with `by = join_by(Well)`
```

###### Removing blanks, setting time column to minutes, subtracting min OD values, and combining datasets

The way I subtracted the blanks (see below) is by splitting the
curves by date, machine, population ID, and st generation first. Then, I
subtracted all measurements from those curves by the minimum OD of those
growth curves. This ensures I’m subtracting the ODs by the appropriate
blanks (those corresponding to the same day and the same supernatant
(ie. filtrate) added).

```
st_gc_merged <- rbind(h1_l1_st_r1_merged, h2_l2_st_r1_merged, h3_l3_st_r1_merged, h4_l4_st_r1_merged, h5_l5_st_r1_merged, h6_l6_st_r1_merged, h7_l7_st_r1_merged, h8_l8_st_r1_merged, h1_l1_st_r2_merged, h2_l2_st_r2_merged, h3_l3_st_r2_merged, h4_l4_st_r2_merged, h5_l5_st_r2_merged, h6_l6_st_r2_merged, h7_l7_st_r2_merged, h8_l8_st_r2_merged) #merging all growth curves

#splitting growth curves by date, machine, pop_id, and st_id to subtract from appropriate blank
st_gc_merged_list <- split(st_gc_merged, list(st_gc_merged$date, st_gc_merged$machine, st_gc_merged$pop_id, st_gc_merged$st))

#removing empty elements from list (not all combinations of all elements occurred)
st_gc_merged_list <- list_drop_empty(st_gc_merged_list)

#subtracting blanks from each group (min value of grouped growth curves)
for (i in 1:length(st_gc_merged_list)) {
    st_gc_merged_list[[i]]$Measurements <- st_gc_merged_list[[i]]$Measurements - min(st_gc_merged_list[[i]]$Measurements)
}

#recombining elements of the list
st_gc_merged <- do.call("rbind", st_gc_merged_list)

#removing blanks
st_gc_merged <- st_gc_merged[st_gc_merged$bac_id!="bl",]

st_gc_merged <- st_gc_merged[st_gc_merged$bac_id!="na",]

#Converting time to minutes
st_gc_merged$Time<-hms(st_gc_merged$Time) #Converting time into Hour-Minutes-Seconds
st_gc_merged$Time<-hour(st_gc_merged$Time)*60+minute(st_gc_merged$Time) #Converting time to just minutes (ignoring seconds)
```

##### calculating GC statistics

```
#Computing percapita derivative of each growth curve
st_gc_merged <- mutate(
group_by(st_gc_merged, machine, date, Well),
percap_deriv = calc_deriv(y = Measurements, x = Time, percapita = TRUE,
blank = 0, window_width_n = 7))

#calculating growth curve parameters (lag time, max percapita growth rate, max density, and area under the curve) of each growth curve
data_sum <- st_gc_merged %>% 
group_by(machine, date, pop_id, evol_treat, bac_id, bac_gen, st, Well) %>% summarize(lag_time = lag_time(x = Time, y = Measurements, deriv = percap_deriv, trans_y="linear"),
max_percap = max(percap_deriv, na.rm = TRUE),
max_dens = max(Measurements),
auc = auc(y = Measurements, x = as.numeric(Time)))
```

```
## `summarise()` has regrouped the output.
## ℹ Summaries were computed grouped by machine, date, pop_id, evol_treat, bac_id,
##   bac_gen, st, and Well.
## ℹ Output is grouped by machine, date, pop_id, evol_treat, bac_id, bac_gen, and
##   st.
## ℹ Use `summarise(.groups = "drop_last")` to silence this message.
## ℹ Use `summarise(.by = c(machine, date, pop_id, evol_treat, bac_id, bac_gen,
##   st, Well))` for per-operation grouping (`?dplyr::dplyr_by`) instead.
```

###### Computing relative (r) values

```
#creating a new dataframe containing only rows from data_sum in which bacteria were grown without supernatant ("NST" rows). Then, averaging the 4 growth curve metrics across replicate NST curves, sorting by bacteria ID and date.. In essence, taking the average value of each growth curve parameter for each bacterial strain on each day.
data_sum_nst <- data_sum %>% subset(st == "NST") %>% group_by(bac_id, date) %>% summarize(nst_lag_time = mean(lag_time), nst_max_percap = mean(max_percap), nst_max_dens = mean(max_dens), nst_auc = mean(auc))
```

```
## `summarise()` has regrouped the output.
## ℹ Summaries were computed grouped by bac_id and date.
## ℹ Output is grouped by bac_id.
## ℹ Use `summarise(.groups = "drop_last")` to silence this message.
## ℹ Use `summarise(.by = c(bac_id, date))` for per-operation grouping
##   (`?dplyr::dplyr_by`) instead.
```

```
#recombining these average NST parameters with the original data frame. This will allow me to compute relative growth parameters (that is, the growth parameter for each curve relative to the average value of that parameter on that day for that strain when grown without st).
data_sum_combined <- merge_dfs(data_sum, data_sum_nst)
```

```
## Joining with `by = join_by(date, bac_id)`
```

```
#the next four lines compute r_measures.. that is, relative measures of the four growth curve parameters for each curve relative to that strain on that day when grown without st
data_sum_combined$r_lag_time <- data_sum_combined$lag_time/data_sum_combined$nst_lag_time
data_sum_combined$r_max_percap <- data_sum_combined$max_percap/data_sum_combined$nst_max_percap
data_sum_combined$r_max_dens <- data_sum_combined$max_dens/data_sum_combined$nst_max_dens
data_sum_combined$r_auc <- data_sum_combined$auc/data_sum_combined$nst_auc

#Now, I'm creating a new dataset that includes only the rows for the ancestral strains. This then allows me to compute the average values for each relative growth curve parameter for each ancestral strain in each supernatant on each date (averaging across the replicates taken for each of these combinations in a given day). Calling these averages "anc_r_measure". Each anc_r_auc value, for example, is the auc of the ancestral strain in a given supernatant treatment divided by the ancestor's auc in the NST treatment.
data_sum_combined_anc <- data_sum_combined %>% subset(evol_treat == "ANC") %>% group_by(date, machine, pop_id, st) %>% summarize(anc_r_lag_time = mean(r_lag_time), anc_r_max_percap = mean(r_max_percap), anc_r_max_dens = mean(r_max_dens), anc_r_auc = mean(r_auc))
```

```
## `summarise()` has regrouped the output.
## ℹ Summaries were computed grouped by date, machine, pop_id, and st.
## ℹ Output is grouped by date, machine, and pop_id.
## ℹ Use `summarise(.groups = "drop_last")` to silence this message.
## ℹ Use `summarise(.by = c(date, machine, pop_id, st))` for per-operation
##   grouping (`?dplyr::dplyr_by`) instead.
```

```
#recombining the dataframe with the average ancestral relative growth parameter values with the prior dataframe with all the other relative values. This will allow me in the next step to calculate the rd_measures. That is, the difference in relative growth curve parameters in a given st treatment compared with the NST treatment between a focal strain and the ancestral strain. 
data_sum_recombined <- merge_dfs(data_sum_combined, data_sum_combined_anc)
```

```
## Joining with `by = join_by(machine, date, pop_id, st)`
```

```
#In the next for lines, I will calculate the rd_measures for the four growth curve parameters by dividing each r_measure by the avg ancestor r_measure from that date in that st treatment.
data_sum_recombined$rd_lag_time <- data_sum_recombined$r_lag_time/data_sum_recombined$anc_r_lag_time
data_sum_recombined$rd_max_percap <- data_sum_recombined$r_max_percap/data_sum_recombined$anc_r_max_percap
data_sum_recombined$rd_max_dens <- data_sum_recombined$r_max_dens/data_sum_recombined$anc_r_max_dens
data_sum_recombined$rd_auc <- data_sum_recombined$r_auc/data_sum_recombined$anc_r_auc

#averaging r_ and rd_ growth curve parameters across replicates
data_sum_recombined_avg <- data_sum_recombined %>% group_by(date, machine, evol_treat, pop_id, bac_gen, bac_id, st) %>% summarize(mean_lag_time = mean(lag_time), mean_max_percap = mean(max_percap), mean_max_dens = mean(max_dens), mean_auc = mean(auc), mean_r_lag_time = mean(r_lag_time), mean_r_max_percap = mean(r_max_percap), mean_r_max_dens = mean(r_max_dens), mean_r_auc = mean(r_auc), mean_rd_lag_time = mean(rd_lag_time), mean_rd_max_percap = mean(rd_max_percap), mean_rd_max_dens = mean(rd_max_dens), mean_rd_auc = mean(rd_auc))
```

```
## `summarise()` has regrouped the output.
## ℹ Summaries were computed grouped by date, machine, evol_treat, pop_id,
##   bac_gen, bac_id, and st.
## ℹ Output is grouped by date, machine, evol_treat, pop_id, bac_gen, and bac_id.
## ℹ Use `summarise(.groups = "drop_last")` to silence this message.
## ℹ Use `summarise(.by = c(date, machine, evol_treat, pop_id, bac_gen, bac_id,
##   st))` for per-operation grouping (`?dplyr::dplyr_by`) instead.
```

```
#Now just one more big problem to solve...
#In order to plot/analyze these rd_measures properly, I need to create a new column reflecting the generation of the st-donors in each curve ("NST" for no st treatment, "gen0000" for the ancestral st, and "genXXXX" for the subsequent sts...).. To achieve this, I'll first split the df in three (NST, anc, and non-anc st donors), as each needs to be treated differently.

#starting with NST rows
data_sum_recombined_avg_nst <- data_sum_recombined_avg %>% subset(st == "NST")

#Now generating a new column for st-donor
data_sum_recombined_avg_nst$st <- rep("NST", length(data_sum_recombined_avg_nst$st))

#Now moving on to the ancestral st rows
data_sum_recombined_avg_anc <- data_sum_recombined_avg %>% subset(st == "HD1_0000" | st=="LD1_0000" | st=="HD2_0000" | st=="LD2_0000" | st=="HD3_0000" | st=="LD3_0000" | st=="HD4_0000" | st=="LD4_0000" | st=="HD5_0000" | st=="LD5_0000" | st=="HD6_0000" | st=="LD6_0000" | st=="HD7_0000" | st=="LD7_0000" | st=="HD8_0000" | st=="LD8_0000")

#Now generating a new column for st-donor generation
data_sum_recombined_avg_anc$st <- rep("0000", length(data_sum_recombined_avg_anc$st))

#Now moving on to the evolved bacteria st-donors
data_sum_recombined_avg_evo <- data_sum_recombined_avg %>% subset(st != "NST" & st != "HD1_0000" & st!="LD1_0000" & st!="HD2_0000" & st!="LD2_0000" & st!="HD3_0000" & st!="LD3_0000" & st!="HD4_0000" & st!="LD4_0000" & st!="HD5_0000" & st!="LD5_0000" & st!="HD6_0000" & st!="LD6_0000" & st!="HD7_0000" & st!="LD7_0000" & st!="HD8_0000" & st!="LD8_0000")

#Now generating a new column for st-donor generations by ripping apart the "st" column to isolate the generation number
vector1 <- unlist(strsplit(data_sum_recombined_avg_evo$st, "_")) #creates a vector twice the length of the dataframe containing the "st" column of the df split into two parts, the pop_id and the "genXXX_st"

vector2 <- rep(0,length(data_sum_recombined_avg_evo$st)) #creates empty vector for loop to fill

#the following for loop essentially subsets vector 1 for just the "genXXX_st" values by selecting every 2nd value from the vector
for (i in 1:length(data_sum_recombined_avg_evo$st)) {
  vector2[i] <- vector1[2*i]
}

#Now we need to repeat the process by subsetting vector 2 for just the "genXXX" part of the character
vector2 <- unlist(strsplit(vector2, "_"))

#now we add vector2 as a new column in the evolved-bacteria dataframe reflecting st-donor gen
data_sum_recombined_avg_evo$st <- vector2

#Now we recombine the three dfs using rbind
data_sum_final <- rbind(data_sum_recombined_avg_nst, data_sum_recombined_avg_anc, data_sum_recombined_avg_evo)

#subsetting the dataset for complete set of st donor gens
data_sum_final_complete <- data_sum_final %>% subset(st == "0000" | st == "0250" | st == "0500" | st == "0750" | st == "1000" | st =="NST")

#removing ancestors and gen 500 bacteria
data_sum_final_complete_1000 <- data_sum_final_complete %>% subset(evol_treat != "ANC") %>% subset(bac_gen != 500)
```

###### Generating Model 1.2

Model 1.2 tests for the effect of evolved treatment (ancestor, HD
endpoint, LD endpoint) on area under the curve in growth curves in the
absence of supernatant (ie. regular growth media).

```
#Subsetting unaggregarted growth curve parameter data for just endpoint and ancestral curves and just curves with no st.
nst_auc_dat_stats <- data_sum %>% subset(bac_gen!= 500 & st == "NST")

#Generating model 1.1.2
mod1.2 <- lmer(data=nst_auc_dat_stats, auc ~ evol_treat + (1|pop_id) + (1|date))
```

```
## boundary (singular) fit: see help('isSingular')
```

```
#Running ANOVA to test for overall effect of evolved treat (ANC vs. HD endpoint vs. LD endpoint)
Anova(mod1.2, type="III")
```

```
## Analysis of Deviance Table (Type III Wald chisquare tests)
## 
## Response: auc
##                Chisq Df Pr(>Chisq)    
## (Intercept) 32679.26  1  < 2.2e-16 ***
## evol_treat    494.42  2  < 2.2e-16 ***
## ---
## Signif. codes:  0 '***' 0.001 '**' 0.01 '*' 0.05 '.' 0.1 ' ' 1
```

```
#Post-hoc contrast across levels of evolved treat
pairs(emmeans(mod1.2, ~evol_treat))
```

```
##  contrast estimate   SE    df t.ratio p.value
##  ANC - HD    142.0 6.52 112.0  21.767 <0.0001
##  ANC - LD     61.6 6.52 112.0   9.452 <0.0001
##  HD - LD     -80.3 7.76  41.9 -10.352 <0.0001
## 
## Degrees-of-freedom method: kenward-roger 
## P value adjustment: tukey method for comparing a family of 3 estimates
```

###### Summarizing Model 1.2

1. AUC of ancestral populations in unmodified media is higher than that
   of LD endpoint populations, which is higher still than that of HD
   endpoint populations.

###### Generating Model 1.3.XXX

Model 1.3.XXX tests the effect of filtrates sampled across
evolutionary timepoints on the relative area-under-the-curve of endpoint
vs. ancestral populations across all 16 evolutionary lines (separate
models for each line). Separate model for each population.

```
#Subsetting data for Model 1.3.HD1
#Just rows from ancestor and endpoint populations, no NST rows, and just rows from HD1 experiment
mod1.3_hd1_dat <- data_sum_recombined %>% subset(bac_gen != 500 & st != "NST" & pop_id == "HD1")

#Generating Model 2.1.HD1
mod1.3.hd1 <- lmer(data=mod1.3_hd1_dat, r_auc ~ evol_treat*st + (1|date))

#Running type III Anova on mod2.1.hd1
Anova(mod1.3.hd1, type="III")

#Running emmeans on mod1.3.hd1
#considers all pairwise comparisons between endpoint and ancestral rAUCs within each filtrate treatment
#repeated below
mod1.3.hd1_output <- as.data.frame(pairs(emmeans(mod1.3.hd1, ~evol_treat:st), by="st"))

#Subsetting data for Model 2.1 HD2.
#Just rows from ancestor and endpoint populations, no NST rows, and just rows from HD2 experiment
mod1.3_hd2_dat <- data_sum_recombined %>% subset(bac_gen != 500 & st != "NST" & pop_id == "HD2")

#Generating Model 2.1.HD2
mod1.3.hd2 <- lmer(data=mod1.3_hd2_dat, r_auc ~ evol_treat*st + (1|date))

#Running type III Anova on mod2.1.hd2
Anova(mod1.3.hd2, type="III")

#Running emmeans on mod1.3.hd2
mod1.3.hd2_output <- as.data.frame(pairs(emmeans(mod1.3.hd2, ~evol_treat:st), by="st"))

#Subsetting data for Model 2.1 HD3.
#Just rows from ancestor and endpoint populations, no NST rows, and just rows from HD2 experiment
mod1.3_hd3_dat <- data_sum_recombined %>% subset(bac_gen != 500 & st != "NST" & pop_id == "HD3")

#Generating Model 2.1.HD2
mod1.3.hd3 <- lmer(data=mod1.3_hd3_dat, r_auc ~ evol_treat*st + (1|date))

#Running type III Anova on mod2.1.hd3
Anova(mod1.3.hd3, type="III")

#Running emmeans on mod1.3.hd3
mod1.3.hd3_output <- as.data.frame(pairs(emmeans(mod1.3.hd3, ~evol_treat:st), by="st"))

#Subsetting data for Model 2.1 HD4.
#Just rows from ancestor and endpoint populations, no NST rows, and just rows from HD2 experiment
mod1.3_hd4_dat <- data_sum_recombined %>% subset(bac_gen != 500 & st != "NST" & pop_id == "HD4")

#Generating Model 2.1.HD2
mod1.3.hd4 <- lmer(data=mod1.3_hd4_dat, r_auc ~ evol_treat*st + (1|date))

#Running type III Anova on mod2.1.hd4
Anova(mod1.3.hd4, type="III")

#Running emmeans on mod1.3.hd4
mod1.3.hd4_output <- as.data.frame(pairs(emmeans(mod1.3.hd4, ~evol_treat:st), by="st"))

#Subsetting data for Model 2.1 HD5.
#Just rows from ancestor and endpoint populations, no NST rows, and just rows from HD2 experiment
mod1.3_hd5_dat <- data_sum_recombined %>% subset(bac_gen != 500 & st != "NST" & pop_id == "HD5")

#Generating Model 2.1.HD2
mod1.3.hd5 <- lmer(data=mod1.3_hd5_dat, r_auc ~ evol_treat*st + (1|date))

#Running type III Anova on mod2.1.hd5
Anova(mod1.3.hd5, type="III")

#Running emmeans on mod1.3.hd5
mod1.3.hd5_output <- as.data.frame(pairs(emmeans(mod1.3.hd5, ~evol_treat:st), by="st"))

#Subsetting data for Model 2.1 HD6.
#Just rows from ancestor and endpoint populations, no NST rows, and just rows from HD2 experiment
mod1.3_hd6_dat <- data_sum_recombined %>% subset(bac_gen != 500 & st != "NST" & pop_id == "HD6")

#Generating Model 2.1.HD2
mod1.3.hd6 <- lmer(data=mod1.3_hd6_dat, r_auc ~ evol_treat*st + (1|date))

#Running type III Anova on mod2.1.hd6
Anova(mod1.3.hd6, type="III")

#Running emmeans on mod1.3.hd6
mod1.3.hd6_output <- as.data.frame(pairs(emmeans(mod1.3.hd6, ~evol_treat:st), by="st"))

#Subsetting data for Model 2.1 HD7.
#Just rows from ancestor and endpoint populations, no NST rows, and just rows from HD2 experiment
mod1.3_hd7_dat <- data_sum_recombined %>% subset(bac_gen != 500 & st != "NST" & pop_id == "HD7")

#Generating Model 2.1.HD2
mod1.3.hd7 <- lmer(data=mod1.3_hd7_dat, r_auc ~ evol_treat*st + (1|date))

#Running type III Anova on mod2.1.hd7
Anova(mod1.3.hd7, type="III")

#Running emmeans on mod1.3.hd7
mod1.3.hd7_output <- as.data.frame(pairs(emmeans(mod1.3.hd7, ~evol_treat:st), by="st"))

#Subsetting data for Model 2.1 HD8
#Just rows from ancestor and endpoint populations, no NST rows, and just rows from HD2 experiment
mod1.3_hd8_dat <- data_sum_recombined %>% subset(bac_gen != 500 & st != "NST" & pop_id == "HD8")

#Generating Model 2.1.HD2
mod1.3.hd8 <- lmer(data=mod1.3_hd8_dat, r_auc ~ evol_treat*st + (1|date))

#Running type III Anova on mod2.1.hd8
Anova(mod1.3.hd8, type="III")

#Running emmeans on mod1.3.hd8
mod1.3.hd8_output <- as.data.frame(pairs(emmeans(mod1.3.hd8, ~evol_treat:st), by="st"))

#Subsetting data for Model 2.1 LD1.
#Just rows from ancestor and endpoint populations, no NST rows, and just rows from HD1 experiment
mod1.3_ld1_dat <- data_sum_recombined %>% subset(bac_gen != 500 & st != "NST" & pop_id == "LD1")

#Generating Model 2.1.LD1
mod1.3.ld1 <- lmer(data=mod1.3_ld1_dat, r_auc ~ evol_treat*st + (1|date))

#Running type III Anova on mod2.1.ld1
Anova(mod1.3.ld1, type="III")

#Running emmeans on mod1.3.ld1
#considers all pairwise comparisons between endpoint and ancestral rAUCs within each filtrate treatment
#repeated below
mod1.3.ld1_output <- as.data.frame(pairs(emmeans(mod1.3.ld1, ~evol_treat:st), by="st"))

#Subsetting data for Model 2.1 LD2.
#Just rows from ancestor and endpoint populations, no NST rows, and just rows from LD2 experiment
mod1.3_ld2_dat <- data_sum_recombined %>% subset(bac_gen != 500 & st != "NST" & pop_id == "LD2")

#Generating Model 2.1.LD2
mod1.3.ld2 <- lmer(data=mod1.3_ld2_dat, r_auc ~ evol_treat*st + (1|date))
```

```
## boundary (singular) fit: see help('isSingular')
```

```
#Running type III Anova on mod2.1.ld2
Anova(mod1.3.ld2, type="III")

#Running emmeans on mod1.3.ld2
mod1.3.ld2_output <- as.data.frame(pairs(emmeans(mod1.3.ld2, ~evol_treat:st), by="st"))

#Subsetting data for Model 2.1 LD3.
#Just rows from ancestor and endpoint populations, no NST rows, and just rows from LD2 experiment
mod1.3_ld3_dat <- data_sum_recombined %>% subset(bac_gen != 500 & st != "NST" & pop_id == "LD3")

#Generating Model 2.1.LD2
mod1.3.ld3 <- lmer(data=mod1.3_ld3_dat, r_auc ~ evol_treat*st + (1|date))

#Running type III Anova on mod2.1.ld3
Anova(mod1.3.ld3, type="III")

#Running emmeans on mod1.3.ld3
mod1.3.ld3_output <- as.data.frame(pairs(emmeans(mod1.3.ld3, ~evol_treat:st), by="st"))

#Subsetting data for Model 2.1 LD4.
#Just rows from ancestor and endpoint populations, no NST rows, and just rows from LD2 experiment
mod1.3_ld4_dat <- data_sum_recombined %>% subset(bac_gen != 500 & st != "NST" & pop_id == "LD4")

#Generating Model 2.1.LD2
mod1.3.ld4 <- lmer(data=mod1.3_ld4_dat, r_auc ~ evol_treat*st + (1|date))

#Running type III Anova on mod2.1.ld4
Anova(mod1.3.ld4, type="III")

#Running emmeans on mod1.3.ld4
mod1.3.ld4_output <- as.data.frame(pairs(emmeans(mod1.3.ld4, ~evol_treat:st), by="st"))

#Subsetting data for Model 2.1 LD5.
#Just rows from ancestor and endpoint populations, no NST rows, and just rows from LD2 experiment
mod1.3_ld5_dat <- data_sum_recombined %>% subset(bac_gen != 500 & st != "NST" & pop_id == "LD5")

#Generating Model 2.1.LD2
mod1.3.ld5 <- lmer(data=mod1.3_ld5_dat, r_auc ~ evol_treat*st + (1|date))

#Running type III Anova on mod2.1.ld5
Anova(mod1.3.ld5, type="III")

#Running emmeans on mod1.3.ld5
mod1.3.ld5_output <- as.data.frame(pairs(emmeans(mod1.3.ld5, ~evol_treat:st), by="st"))

#Subsetting data for Model 2.1 LD6.
#Just rows from ancestor and endpoint populations, no NST rows, and just rows from LD2 experiment
mod1.3_ld6_dat <- data_sum_recombined %>% subset(bac_gen != 500 & st != "NST" & pop_id == "LD6")

#Generating Model 2.1.LD2
mod1.3.ld6 <- lmer(data=mod1.3_ld6_dat, r_auc ~ evol_treat*st + (1|date))

#Running type III Anova on mod2.1.ld6
Anova(mod1.3.ld6, type="III")

#Running emmeans on mod1.3.ld6
mod1.3.ld6_output <- as.data.frame(pairs(emmeans(mod1.3.ld6, ~evol_treat:st), by="st"))

#Subsetting data for Model 2.1 LD7.
#Just rows from ancestor and endpoint populations, no NST rows, and just rows from LD2 experiment
mod1.3_ld7_dat <- data_sum_recombined %>% subset(bac_gen != 500 & st != "NST" & pop_id == "LD7")

#Generating Model 2.1.LD2
mod1.3.ld7 <- lmer(data=mod1.3_ld7_dat, r_auc ~ evol_treat*st + (1|date))

#Running type III Anova on mod2.1.ld7
Anova(mod1.3.ld7, type="III")

#Running emmeans on mod1.3.ld7
mod1.3.ld7_output <- as.data.frame(pairs(emmeans(mod1.3.ld7, ~evol_treat:st), by="st"))

#Subsetting data for Model 2.1 LD8
#Just rows from ancestor and endpoint populations, no NST rows, and just rows from LD2 experiment
mod1.3_ld8_dat <- data_sum_recombined %>% subset(bac_gen != 500 & st != "NST" & pop_id == "LD8")

#Generating Model 2.1.LD2
mod1.3.ld8 <- lmer(data=mod1.3_ld8_dat, r_auc ~ evol_treat*st + (1|date))

#Running type III Anova on mod2.1.ld8
Anova(mod1.3.ld8, type="III")

#Running emmeans on mod1.3.ld8
mod1.3.ld8_output <- as.data.frame(pairs(emmeans(mod1.3.ld8, ~evol_treat:st), by="st"))

#Combining model outputs
mod1.3_output_comb <- rbind(mod1.3.hd1_output, mod1.3.hd2_output, mod1.3.hd3_output, mod1.3.hd4_output, mod1.3.hd5_output, mod1.3.hd6_output, mod1.3.hd7_output, mod1.3.hd8_output, mod1.3.ld1_output, mod1.3.ld2_output, mod1.3.ld3_output, mod1.3.ld4_output, mod1.3.ld5_output, mod1.3.ld6_output, mod1.3.ld7_output, mod1.3.ld8_output)

#Adjusting P-value for multiple comparisons (Tukey method for 80 comparisons)
mod1.3_output_comb$p_val_adj <- mod1.3_output_comb$p.value*80

#Testing for significance (adj P < 0.05?)
#Highlighting three supernatants that had variable responses to st across replicate days
#using "^"
mod1.3_output_comb$sig_test <- ifelse(mod1.3_output_comb$p_val_adj < 0.05, "*", ifelse(mod1.3_output_comb$st == "LD3_0250" | mod1.3_output_comb$st == "LD7_0250" | mod1.3_output_comb$st == "LD7_0500", "^", ""))
```

###### Pasting Model 1.3.XXX output

```
#Pasting results
mod1.3_output_comb
```

```
##  st       contrast estimate     SE df t.ratio p.value p_val_adj sig_test
##  HD1_0000 ANC - HD -0.00790 0.0369 29  -0.214  0.8319  66.55464         
##  HD1_0250 ANC - HD -0.02184 0.0369 29  -0.592  0.5584  44.67134         
##  HD1_0500 ANC - HD -0.19091 0.0369 29  -5.177 <0.0001   0.00124 *       
##  HD1_0750 ANC - HD -0.11374 0.0369 29  -3.084  0.0045   0.35641         
##  HD1_1000 ANC - HD -0.04682 0.0369 29  -1.270  0.2143  17.14618         
##  HD2_0000 ANC - HD  0.00621 0.0158 29   0.393  0.6974  55.79306         
##  HD2_0250 ANC - HD  0.00564 0.0158 29   0.357  0.7240  57.92246         
##  HD2_0500 ANC - HD -0.00947 0.0158 29  -0.598  0.5542  44.33917         
##  HD2_0750 ANC - HD -0.33094 0.0158 29 -20.917 <0.0001   0.00000 *       
##  HD2_1000 ANC - HD -0.06134 0.0158 29  -3.877  0.0006   0.04462 *       
##  HD3_0000 ANC - HD -0.00249 0.0174 29  -0.143  0.8871  70.97079         
##  HD3_0250 ANC - HD -0.42521 0.0174 29 -24.485 <0.0001   0.00000 *       
##  HD3_0500 ANC - HD -0.50239 0.0174 29 -28.929 <0.0001   0.00000 *       
##  HD3_0750 ANC - HD -0.48141 0.0174 29 -27.722 <0.0001   0.00000 *       
##  HD3_1000 ANC - HD -0.03136 0.0174 29  -1.806  0.0813   6.50354         
##  HD4_0000 ANC - HD  0.01669 0.0205 29   0.812  0.4232  33.85801         
##  HD4_0250 ANC - HD  0.00420 0.0205 29   0.205  0.8394  67.14841         
##  HD4_0500 ANC - HD -0.00189 0.0205 29  -0.092  0.9273  74.18293         
##  HD4_0750 ANC - HD -0.48360 0.0205 29 -23.541 <0.0001   0.00000 *       
##  HD4_1000 ANC - HD -0.54606 0.0205 29 -26.581 <0.0001   0.00000 *       
##  HD5_0000 ANC - HD  0.01465 0.0234 29   0.626  0.5360  42.87842         
##  HD5_0250 ANC - HD -0.28000 0.0234 29 -11.968 <0.0001   0.00000 *       
##  HD5_0500 ANC - HD -0.44904 0.0234 29 -19.194 <0.0001   0.00000 *       
##  HD5_0750 ANC - HD -0.01997 0.0234 29  -0.854  0.4003  32.02658         
##  HD5_1000 ANC - HD  0.00728 0.0234 29   0.311  0.7579  60.62970         
##  HD6_0000 ANC - HD  0.02159 0.0153 29   1.415  0.1677  13.41505         
##  HD6_0250 ANC - HD -0.35790 0.0153 29 -23.457 <0.0001   0.00000 *       
##  HD6_0500 ANC - HD -0.41733 0.0153 29 -27.352 <0.0001   0.00000 *       
##  HD6_0750 ANC - HD  0.00585 0.0153 29   0.384  0.7041  56.32728         
##  HD6_1000 ANC - HD -0.01802 0.0153 29  -1.181  0.2471  19.76666         
##  HD7_0000 ANC - HD  0.01903 0.0229 29   0.832  0.4120  32.96241         
##  HD7_0250 ANC - HD -0.43760 0.0229 29 -19.138 <0.0001   0.00000 *       
##  HD7_0500 ANC - HD -0.50988 0.0229 29 -22.299 <0.0001   0.00000 *       
##  HD7_0750 ANC - HD -0.37594 0.0229 29 -16.441 <0.0001   0.00000 *       
##  HD7_1000 ANC - HD -0.31572 0.0229 29 -13.807 <0.0001   0.00000 *       
##  HD8_0000 ANC - HD  0.08712 0.0174 25   5.011 <0.0001   0.00290 *       
##  HD8_0250 ANC - HD  0.03824 0.0246 25   1.555  0.1324  10.59402         
##  HD8_0500 ANC - HD  0.02789 0.0174 25   1.604  0.1212   9.69589         
##  HD8_0750 ANC - HD  0.04485 0.0174 25   2.580  0.0161   1.29190         
##  HD8_1000 ANC - HD  0.02942 0.0174 25   1.692  0.1031   8.24669         
##  LD1_0000 ANC - LD  0.07061 0.0284 29   2.487  0.0189   1.50902         
##  LD1_0250 ANC - LD -0.45344 0.0284 29 -15.974 <0.0001   0.00000 *       
##  LD1_0500 ANC - LD  0.04611 0.0284 29   1.624  0.1151   9.20822         
##  LD1_0750 ANC - LD  0.01900 0.0284 29   0.669  0.5086  40.68563         
##  LD1_1000 ANC - LD -0.00298 0.0284 29  -0.105  0.9172  73.37342         
##  LD2_0000 ANC - LD  0.06949 0.0226 29   3.076  0.0045   0.36320         
##  LD2_0250 ANC - LD -0.33821 0.0226 29 -14.972 <0.0001   0.00000 *       
##  LD2_0500 ANC - LD -0.30372 0.0226 29 -13.445 <0.0001   0.00000 *       
##  LD2_0750 ANC - LD  0.03275 0.0226 29   1.450  0.1579  12.63099         
##  LD2_1000 ANC - LD -0.00571 0.0226 29  -0.253  0.8023  64.18413         
##  LD3_0000 ANC - LD  0.08865 0.0516 29   1.719  0.0963   7.70537         
##  LD3_0250 ANC - LD -0.07504 0.0516 29  -1.455  0.1565  12.51623 ^       
##  LD3_0500 ANC - LD  0.05566 0.0516 29   1.079  0.2895  23.15654         
##  LD3_0750 ANC - LD  0.01635 0.0516 29   0.317  0.7535  60.27665         
##  LD3_1000 ANC - LD  0.01840 0.0516 29   0.357  0.7239  57.91186         
##  LD4_0000 ANC - LD  0.06177 0.0253 29   2.442  0.0209   1.67378         
##  LD4_0250 ANC - LD -0.38357 0.0253 29 -15.165 <0.0001   0.00000 *       
##  LD4_0500 ANC - LD -0.01183 0.0253 29  -0.468  0.6436  51.48463         
##  LD4_0750 ANC - LD  0.01382 0.0253 29   0.546  0.5889  47.11543         
##  LD4_1000 ANC - LD -0.03903 0.0253 29  -1.543  0.1337  10.69232         
##  LD5_0000 ANC - LD  0.05373 0.0201 29   2.677  0.0121   0.96671         
##  LD5_0250 ANC - LD  0.04866 0.0201 29   2.425  0.0218   1.74229         
##  LD5_0500 ANC - LD -0.40022 0.0201 29 -19.943 <0.0001   0.00000 *       
##  LD5_0750 ANC - LD -0.39379 0.0201 29 -19.623 <0.0001   0.00000 *       
##  LD5_1000 ANC - LD  0.00107 0.0201 29   0.053  0.9580  76.64181         
##  LD6_0000 ANC - LD  0.09783 0.0195 29   5.008 <0.0001   0.00199 *       
##  LD6_0250 ANC - LD -0.14479 0.0195 29  -7.412 <0.0001   0.00000 *       
##  LD6_0500 ANC - LD -0.15596 0.0195 29  -7.984 <0.0001   0.00000 *       
##  LD6_0750 ANC - LD  0.06987 0.0195 29   3.577  0.0012   0.09965         
##  LD6_1000 ANC - LD  0.03586 0.0195 29   1.836  0.0767   6.13419         
##  LD7_0000 ANC - LD  0.09167 0.0787 29   1.165  0.2533  20.26685         
##  LD7_0250 ANC - LD -0.12347 0.0787 29  -1.570  0.1273  10.18715 ^       
##  LD7_0500 ANC - LD -0.07618 0.0787 29  -0.968  0.3408  27.26461 ^       
##  LD7_0750 ANC - LD  0.03857 0.0787 29   0.490  0.6275  50.20289         
##  LD7_1000 ANC - LD -0.00521 0.0787 29  -0.066  0.9476  75.81115         
##  LD8_0000 ANC - LD  0.07613 0.0231 29   3.293  0.0026   0.20902         
##  LD8_0250 ANC - LD -0.35429 0.0231 29 -15.326 <0.0001   0.00000 *       
##  LD8_0500 ANC - LD -0.39988 0.0231 29 -17.298 <0.0001   0.00000 *       
##  LD8_0750 ANC - LD  0.04920 0.0231 29   2.128  0.0419   3.35400         
##  LD8_1000 ANC - LD  0.00467 0.0231 29   0.202  0.8412  67.29583         
## 
## Degrees-of-freedom method: kenward-roger
```

```
#Writing CSV file for the output of model 1.2 to be used for plotting significance values
write.csv(mod1.3_output_comb, "~/Documents/PhD Folder/Research/2025 Spring/NCEE - Prophage Paper/Stats Outputs/mod1.3_output_comb.csv", row.names = T)

#Generating model 1.3.1 in response to reviewer comments

#Removing rows for ancestral filtrate
mod1.3_output_comb <- mod1.3_output_comb %>% filter(!str_detect(st, "0000"))

#Generating column for pop_id in mod1.3_output_comb
mod1.3_output_comb$pop_id <- sub("_.*", "", mod1.3_output_comb$st)

#Generating column for treatment in mod1.3_output_comb
mod1.3_output_comb$treat <- substr(mod1.3_output_comb$pop_id, 1, 2)

#generating a 1/0 output from earlier significance tests
mod1.3_output_comb$eco_evo <- if_else(mod1.3_output_comb$sig_test == "", 0, 1)

#Combining all filtrates from eahc population
mod1.3_output_red <- mod1.3_output_comb %>% select(pop_id, treat, eco_evo) %>% group_by(pop_id) %>% summarise(eco_evo = max(eco_evo))

#adding treatment back in
mod1.3_output_red$treat <- substr(mod1.3_output_red$pop_id, 1, 2)

#generating contingency table for Fisher's exact test
#7/8 and 8/8 as frequencies of eco evo feedbacks based on mod1.3_output_red
HD_eco_evo <- c(7,1)
LD_eco_evo <- c(8,0)

mod1.3.1_table <- rbind(HD_eco_evo, LD_eco_evo)

colnames(mod1.3.1_table) <- c("yes", "no")

#Running Fisher's Exact test
mod1.3.1 <- fisher.test(mod1.3.1_table)

mod1.3.1
```

```
## 
##  Fisher's Exact Test for Count Data
## 
## data:  mod1.3.1_table
## p-value = 1
## alternative hypothesis: true odds ratio is not equal to 1
## 95 percent confidence interval:
##   0.00000 39.00055
## sample estimates:
## odds ratio 
##          0
```

###### Summarizing Model 1.3.XXX

1. In all populations excluding HD8, LD3, and LD7, rAUC for endpoint
   populations exceeds that of ancestral populations in filtrate from at
   least one evolutionary timepoint.
2. In all populations, the rAUC of ancestral populations is greater
   than or equal to endpoint populations in ancestral (gen = 0000)
   filtrate.
3. Focusing in on HD7, endpoint populations had higher rAUC than
   ancestral populations in filtrate from generation 500.
4. Based on Fisher’s Exact test, which considers the frequency of
   detecting eco-evolutionary feedbacks in each population combined across
   all filtrate timepoints, there was no detectable difference in the
   frequency of eco-evolutionary feedbacks detected across HD and LD
   populations.

#### Figure 2

The results section centered on Figure 2 involves the following
statistical models:

Model 2.1: relative-area-under-the-curve of ancestral and endpoint
populations in filtrate containing or lacking detectable hyperactive
phage

Model 2.2: area-under-the-curve of MPAO1 in ancestral and evolved
filtrate with or without 10 kDa amicon filtration.

Model 2.3: area-under-the-curve of MPAO1 in evolved filtrate and
ancestral filtrate with phage spiked-in.

##### Model 2.1

###### Importing data

```
st_pfu <- read.csv("~/Documents/PhD Folder/Research/2025 Spring/NCEE - Prophage Paper/ncee_prophage-paper_data/ncee_fil_pfus/ncee_st_pfus.csv")
```

###### Calculating PFU/mL and averaging across replicates

```
#Calculating pfu/mL based on plaque count, dilution, and volume
st_pfu$pfu <- (st_pfu$count*500)*(10^st_pfu$dil)

#making st_gen into gen number (removing x's that were in place to let excel show the necessary lagging zeros)
st_pfu$st_gen <- gsub("x", "", st_pfu$st_gen)

#Averaging PFU/mL across replicates
st_pfu_avg <- st_pfu %>% group_by(pop_id, st_gen, lawn_gen) %>% summarise(mean_pfu = mean(pfu))
```

```
## `summarise()` has regrouped the output.
## ℹ Summaries were computed grouped by pop_id, st_gen, and lawn_gen.
## ℹ Output is grouped by pop_id and st_gen.
## ℹ Use `summarise(.groups = "drop_last")` to silence this message.
## ℹ Use `summarise(.by = c(pop_id, st_gen, lawn_gen))` for per-operation grouping
##   (`?dplyr::dplyr_by`) instead.
```

###### Wrangling data

```
#Subsetting for PFUs on ancestral strain and averaging across replicates within days
st_pfu_anc <- subset(st_pfu, st_pfu$lawn_gen==0) %>% group_by(pop_id, st_gen, date) %>% summarize(pfu=mean(pfu)) %>% mutate(fil_id = paste(pop_id, st_gen, sep="_")) %>% mutate(fil_id_date = paste(fil_id, date, sep="_"))
```

```
## `summarise()` has regrouped the output.
## ℹ Summaries were computed grouped by pop_id, st_gen, and date.
## ℹ Output is grouped by pop_id and st_gen.
## ℹ Use `summarise(.groups = "drop_last")` to silence this message.
## ℹ Use `summarise(.by = c(pop_id, st_gen, date))` for per-operation grouping
##   (`?dplyr::dplyr_by`) instead.
```

```
#Retrieving growth curve parameters for ancestral strain 
data_sum_anc <- data_sum_combined %>% subset(evol_treat == "ANC")  

##Subsetting growth curve parameters for ancestral strain for only curves when ancestor was grown in supernatant
data_sum_anc <- subset(data_sum_anc, data_sum_anc$st!="NST")

#Making new column named "Ancestor" for the mean rAUC, which will be useful for plotting
data_sum_anc$Ancestor <- data_sum_anc$r_auc 

#generating fil_id_date column for merging dfs later
data_sum_anc <- data_sum_anc %>% mutate(fil_id_date = paste(st, date, sep="_"))

#Repeating the above process but for endpoint populations
data_sum_1000 <- data_sum_combined %>% subset(evol_treat != "ANC") %>% subset(bac_gen != 500)

data_sum_1000 <- subset(data_sum_1000, data_sum_1000$st!="NST")

data_sum_1000$Endpoint <- data_sum_1000$r_auc

#generating fil_id_date column for merging dfs later
data_sum_1000 <- data_sum_1000 %>% mutate(fil_id_date = paste(st, date, sep="_"))

#recombining growth curve datasets
data_sum_recomb <- cbind(data_sum_anc, data_sum_1000)
```

```
## New names:
## • `machine` -> `machine...1`
## • `date` -> `date...2`
## • `pop_id` -> `pop_id...3`
## • `evol_treat` -> `evol_treat...4`
## • `bac_id` -> `bac_id...5`
## • `bac_gen` -> `bac_gen...6`
## • `st` -> `st...7`
## • `Well` -> `Well...8`
## • `lag_time` -> `lag_time...9`
## • `max_percap` -> `max_percap...10`
## • `max_dens` -> `max_dens...11`
## • `auc` -> `auc...12`
## • `nst_lag_time` -> `nst_lag_time...13`
## • `nst_max_percap` -> `nst_max_percap...14`
## • `nst_max_dens` -> `nst_max_dens...15`
## • `nst_auc` -> `nst_auc...16`
## • `r_lag_time` -> `r_lag_time...17`
## • `r_max_percap` -> `r_max_percap...18`
## • `r_max_dens` -> `r_max_dens...19`
## • `r_auc` -> `r_auc...20`
## • `fil_id_date` -> `fil_id_date...22`
## • `machine` -> `machine...23`
## • `date` -> `date...24`
## • `pop_id` -> `pop_id...25`
## • `evol_treat` -> `evol_treat...26`
## • `bac_id` -> `bac_id...27`
## • `bac_gen` -> `bac_gen...28`
## • `st` -> `st...29`
## • `Well` -> `Well...30`
## • `lag_time` -> `lag_time...31`
## • `max_percap` -> `max_percap...32`
## • `max_dens` -> `max_dens...33`
## • `auc` -> `auc...34`
## • `nst_lag_time` -> `nst_lag_time...35`
## • `nst_max_percap` -> `nst_max_percap...36`
## • `nst_max_dens` -> `nst_max_dens...37`
## • `nst_auc` -> `nst_auc...38`
## • `r_lag_time` -> `r_lag_time...39`
## • `r_max_percap` -> `r_max_percap...40`
## • `r_max_dens` -> `r_max_dens...41`
## • `r_auc` -> `r_auc...42`
## • `fil_id_date` -> `fil_id_date...44`
```

```
#Generating new column for filtrate ID with a better name for the model
data_sum_recomb$fil_id_date <- data_sum_recomb$fil_id_date...22

#Combining datasets so I can relate PFU data to growth curve data
pfu_r_auc <- merge_dfs(data_sum_recomb, st_pfu_anc)
```

```
## Joining with `by = join_by(fil_id_date)`
```

```
#renaming some columns for convenience
pfu_r_auc <- pfu_r_auc %>% select("pop_id...3", "st_gen", "pfu", "Ancestor", "Endpoint", "date...2") %>% rename(pop_id=pop_id...3) %>% rename(date=date...2) %>% pivot_longer(cols=c("Ancestor", "Endpoint"), names_to="bac_id", values_to="mean_rAUC")

#Making a new column for phage presence
pfu_r_auc$phage_present <- ifelse(pfu_r_auc$pfu>0, "Present", "Not Present")
```

###### Generating Model 2.1

Model 2.1 compares the relative area-under-the-curve of ancestral and
endpoint populations in filtrate containing vs. lacking hyperactive
phages

```
#Generating Model 2.1
mod2.1 <- lmer(data=pfu_r_auc, mean_rAUC ~ bac_id*phage_present + (1|pop_id) + (1|date))

#Looking at overall effects
Anova(mod2.1, type="III")
```

```
## Analysis of Deviance Table (Type III Wald chisquare tests)
## 
## Response: mean_rAUC
##                         Chisq Df Pr(>Chisq)    
## (Intercept)          5911.091  1  < 2.2e-16 ***
## bac_id                 18.775  1   1.47e-05 ***
## phage_present        2645.369  1  < 2.2e-16 ***
## bac_id:phage_present 1462.254  1  < 2.2e-16 ***
## ---
## Signif. codes:  0 '***' 0.001 '**' 0.01 '*' 0.05 '.' 0.1 ' ' 1
```

```
#Examining differences between groups
pairs(emmeans(mod2.1, ~ bac_id:phage_present), by="bac_id")
```

```
## bac_id = Ancestor:
##  contrast              estimate     SE  df t.ratio p.value
##  Not Present - Present   0.3795 0.0074 625  51.305 <0.0001
## 
## bac_id = Endpoint:
##  contrast              estimate     SE  df t.ratio p.value
##  Not Present - Present  -0.0062 0.0074 625  -0.838  0.4026
## 
## Degrees-of-freedom method: kenward-roger
```

###### Model 2.1 Summary

1. The effect of phage presence on the impact of filtrate on
   bacterial growth varied across endpoint and ancestral
   populations.
2. Phage-containing filtrate significantly lowered the rAUC of
   ancestral populations, but not endpoint populations.

##### Model 2.2

###### Importing data

Both Model 2.2 and 2.3 concern data from the Phage Removal/Addition
experiment.

```
phage_ko_r1 <- read_wides(files = "~/Documents/PhD Folder/Research/2025 Spring/NCEE - Prophage Paper/ncee_prophage-paper_data/phage_ko_exp/gcs/2025.03.19/nh_2025.03.19_phage_ko_r1.csv", startrow=27, endrow=166) #Importing single plate of growth curve data (repeated below)

phage_ko_r1 <- trans_wide_to_tidy(wides=phage_ko_r1[,-c(1,2,4)], id_cols=c("Time")) #Converting imported growth curve to tidy format (repeated below)

phage_ko_r2 <- read_wides(files = "~/Documents/PhD Folder/Research/2025 Spring/NCEE - Prophage Paper/ncee_prophage-paper_data/phage_ko_exp/gcs/2025.03.20/nh_2025.03.20_phage_ko_r2.csv", startrow=27, endrow=166)

phage_ko_r2 <- trans_wide_to_tidy(wides=phage_ko_r2[,-c(1,2,4)], id_cols=c("Time"))

phage_ko_r3 <- read_wides(files = "~/Documents/PhD Folder/Research/2025 Spring/NCEE - Prophage Paper/ncee_prophage-paper_data/phage_ko_exp/gcs/2025.03.21/nh_2025.03.21_phage_ko_r3.csv", startrow=27, endrow=166)

phage_ko_r3 <- trans_wide_to_tidy(wides=phage_ko_r3[,-c(1,2,4)], id_cols=c("Time"))
```

###### Importing Design Files

```
phage_ko_r1_design<-import_blockdesigns(files=c("~/Documents/PhD Folder/Research/2025 Spring/NCEE - Prophage Paper/ncee_prophage-paper_data/phage_ko_exp/gcs/2025.03.19/phage_ko_r1_bac_id.csv", "~/Documents/PhD Folder/Research/2025 Spring/NCEE - Prophage Paper/ncee_prophage-paper_data/phage_ko_exp/gcs/2025.03.19/phage_ko_r1_date.csv", "~/Documents/PhD Folder/Research/2025 Spring/NCEE - Prophage Paper/ncee_prophage-paper_data/phage_ko_exp/gcs/2025.03.19/phage_ko_r1_st_evol_treat.csv", "~/Documents/PhD Folder/Research/2025 Spring/NCEE - Prophage Paper/ncee_prophage-paper_data/phage_ko_exp/gcs/2025.03.19/phage_ko_r1_st_pop_id.csv", "~/Documents/PhD Folder/Research/2025 Spring/NCEE - Prophage Paper/ncee_prophage-paper_data/phage_ko_exp/gcs/2025.03.19/phage_ko_r1_st.csv", "~/Documents/PhD Folder/Research/2025 Spring/NCEE - Prophage Paper/ncee_prophage-paper_data/phage_ko_exp/gcs/2025.03.19/phage_ko_r1_treat.csv"), block_names = c("bac_id", "date", "st_evol_treat", "st_pop_id", "st", "treat")) #Importing set of design files for one plate of growth curves (repeated below)
```

```
## Inferred 'into' column names as: bac_id, date, st_evol_treat, st_pop_id, st, treat
```

```
phage_ko_r2_design<-import_blockdesigns(files=c("~/Documents/PhD Folder/Research/2025 Spring/NCEE - Prophage Paper/ncee_prophage-paper_data/phage_ko_exp/gcs/2025.03.20/phage_ko_r2_bac_id.csv", "~/Documents/PhD Folder/Research/2025 Spring/NCEE - Prophage Paper/ncee_prophage-paper_data/phage_ko_exp/gcs/2025.03.20/phage_ko_r2_date.csv", "~/Documents/PhD Folder/Research/2025 Spring/NCEE - Prophage Paper/ncee_prophage-paper_data/phage_ko_exp/gcs/2025.03.20/phage_ko_r2_st_evol_treat.csv", "~/Documents/PhD Folder/Research/2025 Spring/NCEE - Prophage Paper/ncee_prophage-paper_data/phage_ko_exp/gcs/2025.03.20/phage_ko_r2_st_pop_id.csv", "~/Documents/PhD Folder/Research/2025 Spring/NCEE - Prophage Paper/ncee_prophage-paper_data/phage_ko_exp/gcs/2025.03.20/phage_ko_r2_st.csv", "~/Documents/PhD Folder/Research/2025 Spring/NCEE - Prophage Paper/ncee_prophage-paper_data/phage_ko_exp/gcs/2025.03.20/phage_ko_r2_treat.csv"), block_names = c("bac_id", "date", "st_evol_treat", "st_pop_id", "st", "treat"))
```

```
## Inferred 'into' column names as: bac_id, date, st_evol_treat, st_pop_id, st, treat
```

```
phage_ko_r3_design<-import_blockdesigns(files=c("~/Documents/PhD Folder/Research/2025 Spring/NCEE - Prophage Paper/ncee_prophage-paper_data/phage_ko_exp/gcs/2025.03.21/phage_ko_r3_bac_id.csv", "~/Documents/PhD Folder/Research/2025 Spring/NCEE - Prophage Paper/ncee_prophage-paper_data/phage_ko_exp/gcs/2025.03.21/phage_ko_r3_date.csv", "~/Documents/PhD Folder/Research/2025 Spring/NCEE - Prophage Paper/ncee_prophage-paper_data/phage_ko_exp/gcs/2025.03.21/phage_ko_r3_st_evol_treat.csv", "~/Documents/PhD Folder/Research/2025 Spring/NCEE - Prophage Paper/ncee_prophage-paper_data/phage_ko_exp/gcs/2025.03.21/phage_ko_r3_st_pop_id.csv", "~/Documents/PhD Folder/Research/2025 Spring/NCEE - Prophage Paper/ncee_prophage-paper_data/phage_ko_exp/gcs/2025.03.21/phage_ko_r3_st.csv", "~/Documents/PhD Folder/Research/2025 Spring/NCEE - Prophage Paper/ncee_prophage-paper_data/phage_ko_exp/gcs/2025.03.21/phage_ko_r3_treat.csv"), block_names = c("bac_id", "date", "st_evol_treat", "st_pop_id", "st", "treat"))
```

```
## Inferred 'into' column names as: bac_id, date, st_evol_treat, st_pop_id, st, treat
```

###### Merging gcs and design files and wrangling data

```
phage_ko_r1_merged <- merge_dfs(phage_ko_r1, phage_ko_r1_design) #merging dfs (repeated below)
```

```
## Joining with `by = join_by(Well)`
```

```
phage_ko_r2_merged <- merge_dfs(phage_ko_r2, phage_ko_r2_design)
```

```
## Joining with `by = join_by(Well)`
```

```
phage_ko_r3_merged <- merge_dfs(phage_ko_r3, phage_ko_r3_design)
```

```
## Joining with `by = join_by(Well)`
```

```
phage_ko_merged <- rbind(phage_ko_r1_merged, phage_ko_r2_merged, phage_ko_r3_merged) #merging all growth curves

#Splitting curves by date and treatment to subtract blanks from growth curves, as above
phage_ko_merged_list <- split(phage_ko_merged, list(phage_ko_merged$date, phage_ko_merged$treat))

phage_ko_merged_list <- list_drop_empty(phage_ko_merged_list)

for (i in 1:length(phage_ko_merged_list)) {
    phage_ko_merged_list[[i]]$Measurements <- phage_ko_merged_list[[i]]$Measurements - min(phage_ko_merged_list[[i]]$Measurements)
}

#recombining elements of the list
phage_ko_merged <- do.call("rbind", phage_ko_merged_list)

#removing blanks
phage_ko_merged <- phage_ko_merged[phage_ko_merged$"bac_id"!="bl",]

#removing NAs
phage_ko_merged <- phage_ko_merged[phage_ko_merged$"bac_id"!="na",]

#Converting time to minutes
phage_ko_merged$Time<-hms(phage_ko_merged$Time) #Converting time into Hour-Minutes-Seconds
phage_ko_merged$Time<-hour(phage_ko_merged$Time)*60+minute(phage_ko_merged$Time) #Converting time to just minutes (ignoring seconds)
```

##### calculating GC statistics

```
#Computing percapita derivative of each growth curve
phage_ko_merged <- mutate(
group_by(phage_ko_merged, date, st_pop_id, st, treat, Well),
percap_deriv = calc_deriv(y = Measurements, x = Time, percapita = TRUE,
blank = 0, window_width_n = 7))

#calculating growth curve parameters (lag time, max percapita growth rate, max density, and area under the curve) of each growth curve
data_sum <- phage_ko_merged %>% 
group_by(date, st_pop_id, st, treat, Well) %>% summarize(lag_time = lag_time(x = Time, y = Measurements, deriv = percap_deriv, trans_y="linear"),
max_percap = max(percap_deriv, na.rm = TRUE),
max_dens = max(Measurements),
auc = auc(y = Measurements, x = as.numeric(Time)))
```

```
## `summarise()` has regrouped the output.
## ℹ Summaries were computed grouped by date, st_pop_id, st, treat, and Well.
## ℹ Output is grouped by date, st_pop_id, st, and treat.
## ℹ Use `summarise(.groups = "drop_last")` to silence this message.
## ℹ Use `summarise(.by = c(date, st_pop_id, st, treat, Well))` for per-operation
##   grouping (`?dplyr::dplyr_by`) instead.
```

```
#Renaming some treatments for clarity
data_sum$treat <- data_sum$treat %>% replace(data_sum$treat == "anc_st + M9", "anc") %>% replace(data_sum$treat == "evol_st + M9", "evol") %>% replace(data_sum$treat == "anc_st + fil + M9", "anc + fil") %>% replace(data_sum$treat == "evol_st + fil + M9", "evol + fil") %>% replace(data_sum$treat == "evol_st + fil", "evol + fil") %>% replace(data_sum$treat == "evol_st + fil + phage", "evol + fil + Φ") %>% replace(data_sum$treat == "anc_st + phage", "anc + Φ")

#Reordering treatments for clarity
data_sum$treat <- factor(data_sum$treat, levels = c("anc", "evol", "anc + Φ", "anc + fil", "evol + fil", "evol + fil + Φ"))
```

###### Generating Model 2.2

Model 2.2 compares the area-under-the-curve of MPAO1 in ancestral and
evolved filtrate with or without 10 kDa amicon filtration.

```
#Subsetting data for just rows relevant to phage knock out experiment (anc, evol, anc + fil, evol + fil)
mod2.2_dat <- data_sum %>% subset(treat!="anc + Φ" & treat!="NST" & treat!="evol + fil + Φ")

#Creating new column for centrifugal filter treatment (FIL_Y = filtered, FIL_N = unfiltered)
#Creating new column for whether filtrate is ancestral (ANC) or from evolved pops (EVOL)
#and removing na rows
mod2.2_dat <- mod2.2_dat %>% mutate(cent_fil_treat = case_when(grepl("fil", treat) ~ "FIL_Y", TRUE ~ "FIL_N")) %>% mutate(evol_fil_treat = case_when(grepl("anc", treat) ~ "ANC", TRUE ~ "EVOL")) %>% subset(st != "na")

#Generating Model 2.2
mod2.2 <- lmer(data=mod2.2_dat, auc ~ evol_fil_treat*cent_fil_treat + (1|date))

#Summarizing Model 2.2
summary(mod2.2)
```

```
## Linear mixed model fit by REML. t-tests use Satterthwaite's method [
## lmerModLmerTest]
## Formula: auc ~ evol_fil_treat * cent_fil_treat + (1 | date)
##    Data: mod2.2_dat
## 
## REML criterion at convergence: 1527.8
## 
## Scaled residuals: 
##      Min       1Q   Median       3Q      Max 
## -2.53310 -0.25643 -0.01932  0.42186  3.08156 
## 
## Random effects:
##  Groups   Name        Variance Std.Dev.
##  date     (Intercept)  150.6   12.27   
##  Residual             6691.7   81.80   
## Number of obs: 134, groups:  date, 3
## 
## Fixed effects:
##                                        Estimate Std. Error      df t value
## (Intercept)                              705.01      24.65   51.44  28.596
## evol_fil_treatEVOL                      -209.24      25.87  127.93  -8.089
## cent_fil_treatFIL_Y                      213.53      40.90  127.93   5.221
## evol_fil_treatEVOL:cent_fil_treatFIL_Y   221.71      43.64  127.97   5.081
##                                        Pr(>|t|)    
## (Intercept)                             < 2e-16 ***
## evol_fil_treatEVOL                     4.00e-13 ***
## cent_fil_treatFIL_Y                    7.01e-07 ***
## evol_fil_treatEVOL:cent_fil_treatFIL_Y 1.30e-06 ***
## ---
## Signif. codes:  0 '***' 0.001 '**' 0.01 '*' 0.05 '.' 0.1 ' ' 1
## 
## Correlation of Fixed Effects:
##             (Intr) ev__EVOL c__FIL
## evl_fl_EVOL -0.874                
## cnt_f_FIL_Y -0.553  0.527         
## e__EVOL:__F  0.518 -0.593   -0.937
```

```
#Running Type III Anova on Model 2.2
Anova(mod2.2, type="III")
```

```
## Analysis of Deviance Table (Type III Wald chisquare tests)
## 
## Response: auc
##                                 Chisq Df Pr(>Chisq)    
## (Intercept)                   817.706  1  < 2.2e-16 ***
## evol_fil_treat                 65.429  1  6.024e-16 ***
## cent_fil_treat                 27.255  1  1.784e-07 ***
## evol_fil_treat:cent_fil_treat  25.815  1  3.758e-07 ***
## ---
## Signif. codes:  0 '***' 0.001 '**' 0.01 '*' 0.05 '.' 0.1 ' ' 1
```

```
#Running emmeans on Model 2.2 to generate CIs for AUC across filtrates
pairs(emmeans(mod2.2, ~ evol_fil_treat:cent_fil_treat), by="cent_fil_treat")
```

```
## cent_fil_treat = FIL_N:
##  contrast   estimate   SE  df t.ratio p.value
##  ANC - EVOL    209.2 25.9 128   8.089 <0.0001
## 
## cent_fil_treat = FIL_Y:
##  contrast   estimate   SE  df t.ratio p.value
##  ANC - EVOL    -12.5 35.2 128  -0.355  0.7234
## 
## Degrees-of-freedom method: kenward-roger
```

###### Summary of Model 2.2

1. Without centrifugal filtration, evolved filtrate reduces the
   growth of MPAO1 relative to ancestral filtrate
2. Centrifugal filtration of either class of filtrate increases the
   growth of MPAO1
3. Centrifugal filtration especially increases the growth of MPAO1
   in evolved filtrate
4. The growth of MPAO1 in filtrate from evolved and ancestral
   populations that has been treated with a centrifugal filter is
   indistinguishable.

###### Generating Model 2.3

Model 2.3 compared the area-under-the-curve of MPAO1 in evolved
filtrate and ancestral filtrate with phage spiked-in.

```
#Subsetting for just evolved filtrate and ancestral filtrate with phage spiked in
mod2.3_dat <- data_sum %>% subset(treat=="anc + Φ" | treat=="evol")

#removing rows with na
mod2.3_dat <- mod2.3_dat  %>% subset(st != "na")

#Generating Model 2.3
mod2.3 <- lmer(data=mod2.3_dat, auc ~ treat*st + (1|date))

#Running type III Anova on Model 2.3
Anova(mod2.3, type="III")
```

```
## Analysis of Deviance Table (Type III Wald chisquare tests)
## 
## Response: auc
##                Chisq Df Pr(>Chisq)    
## (Intercept) 1388.977  1    < 2e-16 ***
## treat          3.547  1    0.05965 .  
## st           817.164  9    < 2e-16 ***
## treat:st      20.847  9    0.01335 *  
## ---
## Signif. codes:  0 '***' 0.001 '**' 0.01 '*' 0.05 '.' 0.1 ' ' 1
```

```
# Running emmeans on Model 2.3 to look at CIs for each combination of treatment and filtrate ID
pairs(emmeans(mod2.3, ~treat:st), by = "st")
```

```
## st = HD1_0500:
##  contrast         estimate   SE df t.ratio p.value
##  evol - (anc + Φ)  -32.856 17.4 96  -1.883  0.0627
## 
## st = HD1_0750:
##  contrast         estimate   SE df t.ratio p.value
##  evol - (anc + Φ)  -32.810 17.4 96  -1.881  0.0630
## 
## st = HD7_0250:
##  contrast         estimate   SE df t.ratio p.value
##  evol - (anc + Φ)   12.780 17.4 96   0.733  0.4656
## 
## st = HD7_0500:
##  contrast         estimate   SE df t.ratio p.value
##  evol - (anc + Φ)   -0.167 17.4 96  -0.010  0.9924
## 
## st = HD7_0750:
##  contrast         estimate   SE df t.ratio p.value
##  evol - (anc + Φ)   41.208 18.3 96   2.250  0.0267
## 
## st = HD7_1000:
##  contrast         estimate   SE df t.ratio p.value
##  evol - (anc + Φ)    7.888 17.4 96   0.452  0.6522
## 
## st = LD6_0250:
##  contrast         estimate   SE df t.ratio p.value
##  evol - (anc + Φ)   11.036 18.3 96   0.603  0.5482
## 
## st = LD6_0500:
##  contrast         estimate   SE df t.ratio p.value
##  evol - (anc + Φ)   21.115 17.4 96   1.210  0.2291
## 
## st = LD8_0250:
##  contrast         estimate   SE df t.ratio p.value
##  evol - (anc + Φ)    1.492 17.4 96   0.086  0.9320
## 
## st = LD8_0500:
##  contrast         estimate   SE df t.ratio p.value
##  evol - (anc + Φ)  -43.602 17.4 96  -2.499  0.0141
## 
## Degrees-of-freedom method: kenward-roger
```

###### Summary of Model 2.3

1. The effect of filtrates on the performance of MPAO1 did not
   differ between phage-containing evolved filtrate and ancestral filtrate
   with phage spiked in.
2. There was a strong effect of filtrate ID on the performance of
   MPAO1. Filtrate IDs vary in both the evolved filtrate and the phage ID
   and phage titer spiked into the ancestral filtrate.
3. The impact of treatment (evol vs. anc + phage) on MPAO1
   performance varied across filtrate IDs. This implies that the impact of
   evol or anc + phage filtrates had a different effect in some filtrates
   (in some cases, evol inhibited MPAO1 more, or vice versa)
4. Once correcting for multiple comparisons, there was no detectable
   difference in the effect of evol vs. anc + phage filtrate for any
   filtrate ID.

#### Figure 3

The results section centered on Figure 3 involves the following
statistical models:

Model 3.1.1: AUC of ancestral strain grown in the presence of
hyperactive Pf isolates (across phage populations).

Model 3.1.2: AUC of ancestral strain grown in the presence of
hyperactive Pf isolates (across generations in HD7).

Model 3.2: Relationship between rAUC of ancestral strain in
phage-containing filtrate and the AUC of the ancestral strain when grown
in the presence of isolates from the same filtrate, as well as the titer
of phage in that filtrate.

##### Model 3.1

Model 3.1 compares the AUC of MPAO1 (ancestral strain) when
co-cultured with hyperactive phage isolates (moi: 0.01).

###### Importing Data

```
si_phage_r1 <- read_wides(files = "~/Documents/PhD Folder/Research/2025 Spring/NCEE - Prophage Paper/ncee_prophage-paper_data/si_phage_gcs/2025.03.25_si_phage_r1/nh_2025.03.25_si_phage_r1.csv", startrow=27, endrow=166) #Importing single plate of growth curve data (repeated below)

si_phage_r1 <- trans_wide_to_tidy(wides=si_phage_r1[,-c(1,2,4)], id_cols=c("Time"))  #Converting imported growth curve to tidy format (repeated below)

si_phage_r2 <- read_wides(files = "~/Documents/PhD Folder/Research/2025 Spring/NCEE - Prophage Paper/ncee_prophage-paper_data/si_phage_gcs/2025.03.26_si_phage_r2/nh_2025.03.26_si_phage_r2.csv", startrow=27, endrow=166)

si_phage_r2 <- trans_wide_to_tidy(wides=si_phage_r2[,-c(1,2,4)], id_cols=c("Time")) 

si_phage_r3 <- read_wides(files = "~/Documents/PhD Folder/Research/2025 Spring/NCEE - Prophage Paper/ncee_prophage-paper_data/si_phage_gcs/2025.03.30_si_phage_r3/nh_2025.03.30_si_phage_r3.csv", startrow=28, endrow=167)

si_phage_r3 <- trans_wide_to_tidy(wides=si_phage_r3[,-c(1,2,4)], id_cols=c("Time"))
```

###### Importing design files

```
si_phage_r1_design<-import_blockdesigns(files=c("~/Documents/PhD Folder/Research/2025 Spring/NCEE - Prophage Paper/ncee_prophage-paper_data/si_phage_gcs/2025.03.25_si_phage_r1/si_phage_r1_bac_id.csv", "~/Documents/PhD Folder/Research/2025 Spring/NCEE - Prophage Paper/ncee_prophage-paper_data/si_phage_gcs/2025.03.25_si_phage_r1/si_phage_r1_date.csv",  "~/Documents/PhD Folder/Research/2025 Spring/NCEE - Prophage Paper/ncee_prophage-paper_data/si_phage_gcs/2025.03.25_si_phage_r1/si_phage_r1_phage_evol_treat.csv", "~/Documents/PhD Folder/Research/2025 Spring/NCEE - Prophage Paper/ncee_prophage-paper_data/si_phage_gcs/2025.03.25_si_phage_r1/si_phage_r1_phage_gen.csv", "~/Documents/PhD Folder/Research/2025 Spring/NCEE - Prophage Paper/ncee_prophage-paper_data/si_phage_gcs/2025.03.25_si_phage_r1/si_phage_r1_phage_pop_id.csv", "~/Documents/PhD Folder/Research/2025 Spring/NCEE - Prophage Paper/ncee_prophage-paper_data/si_phage_gcs/2025.03.25_si_phage_r1/si_phage_r1_phage.csv", "~/Documents/PhD Folder/Research/2025 Spring/NCEE - Prophage Paper/ncee_prophage-paper_data/si_phage_gcs/2025.03.25_si_phage_r1/si_phage_r1_phage_pop_gen.csv"), block_names = c("bac_id", "date", "phage_evol_treat", "phage_gen", "phage_pop_id", "phage_id", "phage_pop_gen")) #Importing set of design files for one plate of growth curves (repeated below)
```

```
## Inferred 'into' column names as: bac_id, date, phage_evol_treat, phage_gen, phage_pop_id, phage_id, phage_pop_gen
```

```
si_phage_r2_design<-import_blockdesigns(files=c("~/Documents/PhD Folder/Research/2025 Spring/NCEE - Prophage Paper/ncee_prophage-paper_data/si_phage_gcs/2025.03.26_si_phage_r2/si_phage_r2_bac_id.csv", "~/Documents/PhD Folder/Research/2025 Spring/NCEE - Prophage Paper/ncee_prophage-paper_data/si_phage_gcs/2025.03.26_si_phage_r2/si_phage_r2_date.csv",  "~/Documents/PhD Folder/Research/2025 Spring/NCEE - Prophage Paper/ncee_prophage-paper_data/si_phage_gcs/2025.03.26_si_phage_r2/si_phage_r2_phage_evol_treat.csv", "~/Documents/PhD Folder/Research/2025 Spring/NCEE - Prophage Paper/ncee_prophage-paper_data/si_phage_gcs/2025.03.26_si_phage_r2/si_phage_r2_phage_gen.csv", "~/Documents/PhD Folder/Research/2025 Spring/NCEE - Prophage Paper/ncee_prophage-paper_data/si_phage_gcs/2025.03.26_si_phage_r2/si_phage_r2_phage_pop_id.csv", "~/Documents/PhD Folder/Research/2025 Spring/NCEE - Prophage Paper/ncee_prophage-paper_data/si_phage_gcs/2025.03.26_si_phage_r2/si_phage_r2_phage.csv", "~/Documents/PhD Folder/Research/2025 Spring/NCEE - Prophage Paper/ncee_prophage-paper_data/si_phage_gcs/2025.03.26_si_phage_r2/si_phage_r2_phage_pop_gen.csv"), block_names = c("bac_id", "date", "phage_evol_treat", "phage_gen", "phage_pop_id", "phage_id", "phage_pop_gen"))
```

```
## Inferred 'into' column names as: bac_id, date, phage_evol_treat, phage_gen, phage_pop_id, phage_id, phage_pop_gen
```

```
si_phage_r3_design<-import_blockdesigns(files=c("~/Documents/PhD Folder/Research/2025 Spring/NCEE - Prophage Paper/ncee_prophage-paper_data/si_phage_gcs/2025.03.30_si_phage_r3/si_phage_r3_bac_id.csv", "~/Documents/PhD Folder/Research/2025 Spring/NCEE - Prophage Paper/ncee_prophage-paper_data/si_phage_gcs/2025.03.30_si_phage_r3/si_phage_r3_date.csv",  "~/Documents/PhD Folder/Research/2025 Spring/NCEE - Prophage Paper/ncee_prophage-paper_data/si_phage_gcs/2025.03.30_si_phage_r3/si_phage_r3_phage_evol_treat.csv", "~/Documents/PhD Folder/Research/2025 Spring/NCEE - Prophage Paper/ncee_prophage-paper_data/si_phage_gcs/2025.03.30_si_phage_r3/si_phage_r3_phage_gen.csv", "~/Documents/PhD Folder/Research/2025 Spring/NCEE - Prophage Paper/ncee_prophage-paper_data/si_phage_gcs/2025.03.30_si_phage_r3/si_phage_r3_phage_pop_id.csv", "~/Documents/PhD Folder/Research/2025 Spring/NCEE - Prophage Paper/ncee_prophage-paper_data/si_phage_gcs/2025.03.30_si_phage_r3/si_phage_r3_phage.csv", "~/Documents/PhD Folder/Research/2025 Spring/NCEE - Prophage Paper/ncee_prophage-paper_data/si_phage_gcs/2025.03.30_si_phage_r3/si_phage_r3_phage_pop_gen.csv"), block_names = c("bac_id", "date", "phage_evol_treat", "phage_gen", "phage_pop_id", "phage_id", "phage_pop_gen"))
```

```
## Inferred 'into' column names as: bac_id, date, phage_evol_treat, phage_gen, phage_pop_id, phage_id, phage_pop_gen
```

###### Merging gcs and design files

```
si_phage_r1_merged <- merge_dfs(si_phage_r1, si_phage_r1_design) #merging growth curve data and design files, repeated below
```

```
## Joining with `by = join_by(Well)`
```

```
si_phage_r2_merged <- merge_dfs(si_phage_r2, si_phage_r2_design)
```

```
## Joining with `by = join_by(Well)`
```

```
si_phage_r3_merged <- merge_dfs(si_phage_r3, si_phage_r3_design)
```

```
## Joining with `by = join_by(Well)`
```

```
si_gc_merged <- rbind(si_phage_r1_merged, si_phage_r2_merged, si_phage_r3_merged) #merging all growth curves

#Splitting growth curves by data to subtract blanks, as above
si_gc_merged_list <- split(si_gc_merged, list(si_gc_merged$date))

si_gc_merged_list <- list_drop_empty(si_gc_merged_list)

for (i in 1:length(si_gc_merged_list)) {
    si_gc_merged_list[[i]]$Measurements <- si_gc_merged_list[[i]]$Measurements - min(si_gc_merged_list[[i]]$Measurements)
}

#recombining elemetns of the list
si_gc_merged <- do.call("rbind", si_gc_merged_list)

#removing blanks
si_gc_merged <- si_gc_merged[si_gc_merged$bac_id!="bl",]

#Converting time to minutes
si_gc_merged$Time<-hms(si_gc_merged$Time) #Converting time into Hour-Minutes-Seconds
si_gc_merged$Time<-hour(si_gc_merged$Time)*60+minute(si_gc_merged$Time) #Converting time to just minutes (ignoring seconds)
```

###### Computing growth curve parameters

```
#Computing percapita derivative of each growth curve
si_gc_merged <- mutate(
group_by(si_gc_merged, date, phage_id, phage_pop_id, phage_pop_gen, Well),
percap_deriv = calc_deriv(y = Measurements, x = Time, percapita = TRUE,
blank = 0, window_width_n = 7))

#calculating growth curve parameters (lag time, max percapita growth rate, max density, and area under the curve) of each growth curve
data_sum_si <- si_gc_merged %>% 
group_by(date, phage_id, phage_pop_id, phage_pop_gen, Well) %>% summarize(lag_time = lag_time(x = Time, y = Measurements, deriv = percap_deriv, trans_y="linear"),
max_percap = max(percap_deriv, na.rm = TRUE),
max_dens = max(Measurements),
auc = auc(y = Measurements, x = as.numeric(Time)))
```

```
## `summarise()` has regrouped the output.
## ℹ Summaries were computed grouped by date, phage_id, phage_pop_id,
##   phage_pop_gen, and Well.
## ℹ Output is grouped by date, phage_id, phage_pop_id, and phage_pop_gen.
## ℹ Use `summarise(.groups = "drop_last")` to silence this message.
## ℹ Use `summarise(.by = c(date, phage_id, phage_pop_id, phage_pop_gen, Well))`
##   for per-operation grouping (`?dplyr::dplyr_by`) instead.
```

```
#Making new columns for phage_gen and phage_rep by splitting phage_ID, and then adding phage_ID back in
phage_id <- data_sum_si$phage_id

data_sum_si <- data_sum_si %>% separate(phage_id,c(NA, "phage_gen", "phage_rep"))
```

```
## Warning: Expected 3 pieces. Missing pieces filled with `NA` in 9 rows [91, 92, 93, 184,
## 185, 186, 277, 278, 279].
```

```
data_sum_si$phage_id <- phage_id
```

###### Generating Model 3.1.1

Model 3.1.1 compares the AUC of ancestral strain grown in the
presence of hyperactive Pf isolates (across phage populations).

```
#Generating Model 3.1.1
mod3.1.1 <- lmer(data=data_sum_si[data_sum_si$phage_pop_id != "no_phage",], auc ~ phage_pop_id + (1|date) + (1|phage_pop_gen/phage_id))

#Testing for overall significance of phage_pop_id
Anova(mod3.1.1, type="III")
```

```
## Analysis of Deviance Table (Type III Wald chisquare tests)
## 
## Response: auc
##               Chisq Df Pr(>Chisq)    
## (Intercept)  346.83  1  < 2.2e-16 ***
## phage_pop_id  47.66  3  2.515e-10 ***
## ---
## Signif. codes:  0 '***' 0.001 '**' 0.01 '*' 0.05 '.' 0.1 ' ' 1
```

```
#testing which pairs of phage_pop_id are statistically distinguishable
pairs(emmeans(mod3.1.1, ~phage_pop_id))
```

```
##  contrast  estimate   SE df t.ratio p.value
##  HD1 - HD7   283.18 49.0  6   5.774  0.0048
##  HD1 - LD6    78.02 56.6  6   1.378  0.5540
##  HD1 - LD8   292.29 56.6  6   5.161  0.0083
##  HD7 - LD6  -205.16 49.0  6  -4.183  0.0224
##  HD7 - LD8     9.11 49.0  6   0.186  0.9975
##  LD6 - LD8   214.27 56.6  6   3.783  0.0347
## 
## Degrees-of-freedom method: kenward-roger 
## P value adjustment: tukey method for comparing a family of 4 estimates
```

###### Summary of Model 3.1.1

1. The AUC of MPAO1 when grown with Pf phage isolates varied
   depending on which population the isolates were sampled from.
2. Phage isolates from HD7 and LD8 decreased the AUC of MPAO1 more
   than phage isolates from HD1 and LD6.

###### Generating Model 3.1.2

Model 3.1.2 compares the AUC of ancestral strain grown in the
presence of hyperactive Pf isolates (across timepoints in phages derived
from population HD7).

```
#Generating Model 3.1.2
mod3.1.2 <- lmer(data=data_sum_si[data_sum_si$phage_pop_id == "HD7",], auc ~ phage_gen + (1|date) + (1|phage_id))

#Testing for overall significance of phage_pop_id
Anova(mod3.1.2, type="III")
```

```
## Analysis of Deviance Table (Type III Wald chisquare tests)
## 
## Response: auc
##               Chisq Df Pr(>Chisq)    
## (Intercept) 582.613  1  < 2.2e-16 ***
## phage_gen    41.857  3  4.302e-09 ***
## ---
## Signif. codes:  0 '***' 0.001 '**' 0.01 '*' 0.05 '.' 0.1 ' ' 1
```

```
#testing which pairs of phage_pop_id are statistically distinguishable
pairs(emmeans(mod3.1.2, ~phage_gen))
```

```
##  contrast                     estimate   SE df t.ratio p.value
##  phage_gen1000 - phage_gen250    78.67 28.9  8   2.725  0.0980
##  phage_gen1000 - phage_gen500   186.00 28.9  8   6.444  0.0009
##  phage_gen1000 - phage_gen750    86.03 28.9  8   2.980  0.0684
##  phage_gen250 - phage_gen500    107.32 28.9  8   3.718  0.0244
##  phage_gen250 - phage_gen750      7.35 28.9  8   0.255  0.9937
##  phage_gen500 - phage_gen750    -99.97 28.9  8  -3.463  0.0347
## 
## Degrees-of-freedom method: kenward-roger 
## P value adjustment: tukey method for comparing a family of 4 estimates
```

###### Summary of Model 3.1.2

1. The AUC of MPAO1 when grown with HD7-derived Pf phage isolates
   varied depending on which evolutionary timepoint isolates were sampled
   from.
2. HD7-derived phage isolates initially increased in virulence (AUC
   gen 500 < AUC gen 250), but virulence was attentuated in subsequent
   generations (AUC gen 500 < AUC gen 750 = AUC gen 1000).

##### Model 3.2

Model 3.2 examines the relationship between rAUC of the ancestral
strain in phage-containing filtrate and the AUC of the ancestral strain
when grown in the presence of isolates from the same filtrate as well as
the titer of phage in that filtrate.

###### Importing and wrangling data

```
#subsetting for no phage wells
data_sum_si_no_phage <- data_sum_si[data_sum_si$phage_pop_id=="no_phage",]

#calculating mean AUC of MPAO1 in absence of phage
data_sum_si_no_phage_sum <- data_sum_si_no_phage %>% group_by(date) %>% summarize(mean_AUC_np = mean(auc))

#Merging summarized no_phage df with phage_df by date
data_sum_merged <- merge_dfs(data_sum_si, data_sum_si_no_phage_sum, by="date")

#calculating rAUC of for virulent phage growth curves (AUC MPAO1 w/phage / AUC MPAO1 w/o phage within day)
data_sum_merged$rAUC <- data_sum_merged$auc/data_sum_merged$mean_AUC_np

#subsetting for curves with phage
data_sum_merged_phage <- data_sum_merged[data_sum_merged$phage_pop_id!="no_phage",]

#averaging across replicates and phage isolates (one rauc value per filtrate)
data_sum_merged_phage <- data_sum_merged_phage %>% group_by(phage_gen, phage_pop_id) %>% summarize(mean_rAUC_si_phage = mean(rAUC))
```

```
## `summarise()` has regrouped the output.
## ℹ Summaries were computed grouped by phage_gen and phage_pop_id.
## ℹ Output is grouped by phage_gen.
## ℹ Use `summarise(.groups = "drop_last")` to silence this message.
## ℹ Use `summarise(.by = c(phage_gen, phage_pop_id))` for per-operation grouping
##   (`?dplyr::dplyr_by`) instead.
```

```
#fixing phage_gen column
data_sum_merged_phage$phage_gen <- c("1000", "0250", "0250", "0250", "0500", "0500", "0500", "0500", "0750", "0750")

#Fixing order of rows
data_sum_merged_phage <- data_sum_merged_phage %>% arrange(phage_gen) %>% arrange(phage_pop_id)

#retrieving st_pfu data and subsetting for phage plated on lawns of anc & only those filtrates from which phage isolates were taken
st_pfu_anc <- st_pfu_avg[st_pfu_avg$lawn_gen==0,]

st_pfu_anc_hd1_focal <- st_pfu_anc[st_pfu_anc$pop_id=="HD1" & st_pfu_anc$st_gen=="0500" | st_pfu_anc$pop_id=="HD1" & st_pfu_anc$st_gen=="0750",]
st_pfu_anc_hd7_focal <- st_pfu_anc[st_pfu_anc$pop_id=="HD7" & st_pfu_anc$st_gen=="0250" | st_pfu_anc$pop_id=="HD7" & st_pfu_anc$st_gen=="0500" | st_pfu_anc$pop_id=="HD7" & st_pfu_anc$st_gen=="0750" | st_pfu_anc$pop_id=="HD7" & st_pfu_anc$st_gen=="1000",]
st_pfu_anc_ld6_focal <- st_pfu_anc[st_pfu_anc$pop_id=="LD6" & st_pfu_anc$st_gen=="0250" | st_pfu_anc$pop_id=="LD6" & st_pfu_anc$st_gen=="0500",]
st_pfu_anc_ld8_focal <- st_pfu_anc[st_pfu_anc$pop_id=="LD8" & st_pfu_anc$st_gen=="0250" | st_pfu_anc$pop_id=="LD8" & st_pfu_anc$st_gen=="0500",]

st_pfu_anc_focal <- rbind(st_pfu_anc_hd1_focal, st_pfu_anc_hd7_focal, st_pfu_anc_ld6_focal, st_pfu_anc_ld8_focal)

#Doing same thing but with rAUC values from bulk filtrate
data_sum_final_complete_anc <- data_sum_final_complete %>% subset(evol_treat == "ANC") %>% group_by(pop_id, st) %>% summarize(mean_r_auc = mean(mean_r_auc))
```

```
## `summarise()` has regrouped the output.
## ℹ Summaries were computed grouped by pop_id and st.
## ℹ Output is grouped by pop_id.
## ℹ Use `summarise(.groups = "drop_last")` to silence this message.
## ℹ Use `summarise(.by = c(pop_id, st))` for per-operation grouping
##   (`?dplyr::dplyr_by`) instead.
```

```
rauc_data_anc <- data_sum_final_complete_anc

rauc_data_anc_hd1_focal <- rauc_data_anc[rauc_data_anc$pop_id=="HD1" & rauc_data_anc$st=="0500" | rauc_data_anc$pop_id=="HD1" & rauc_data_anc$st=="0750",]
rauc_data_anc_hd7_focal <- rauc_data_anc[rauc_data_anc$pop_id=="HD7" & rauc_data_anc$st=="0250" | rauc_data_anc$pop_id=="HD7" & rauc_data_anc$st=="0500" | rauc_data_anc$pop_id=="HD7" & rauc_data_anc$st=="0750" | rauc_data_anc$pop_id=="HD7" & rauc_data_anc$st=="1000",]
rauc_data_anc_ld6_focal <- rauc_data_anc[rauc_data_anc$pop_id=="LD6" & rauc_data_anc$st=="0250" | rauc_data_anc$pop_id=="LD6" & rauc_data_anc$st=="0500",]
rauc_data_anc_ld8_focal <- rauc_data_anc[rauc_data_anc$pop_id=="LD8" & rauc_data_anc$st=="0250" | rauc_data_anc$pop_id=="LD8" & rauc_data_anc$st=="0500",]

rauc_anc_focal <- rbind(rauc_data_anc_hd1_focal, rauc_data_anc_hd7_focal, rauc_data_anc_ld6_focal, rauc_data_anc_ld8_focal)

#Adding rows from filtrate rAUC and PFUs
data_sum_merged_phage$pfu <- st_pfu_anc_focal$mean_pfu
data_sum_merged_phage$rAUC_fil <- rauc_anc_focal$mean_r_auc
```

###### Generating Model 3.2

Model 3.2 examines the relationship between rAUC of the ancestral
strain in phage-containing filtrate and the AUC of the ancestral strain
when grown in the presence of isolates from the same filtrate as well as
the titer of phage in that filtrate.

```
#Generating Model 3.2
mod3.2 <- lmer(data=data_sum_merged_phage, rAUC_fil ~ mean_rAUC_si_phage + log10(pfu) + (1|phage_pop_id))
```

```
## boundary (singular) fit: see help('isSingular')
```

```
#Generating output of Model 3.2
Anova(mod3.2, type="III")
```

```
## Analysis of Deviance Table (Type III Wald chisquare tests)
## 
## Response: rAUC_fil
##                      Chisq Df Pr(>Chisq)    
## (Intercept)         0.1010  1     0.7506    
## mean_rAUC_si_phage 30.3781  1  3.555e-08 ***
## log10(pfu)          0.3623  1     0.5472    
## ---
## Signif. codes:  0 '***' 0.001 '**' 0.01 '*' 0.05 '.' 0.1 ' ' 1
```

```
#Generating output of Model 3.2
summary(mod3.2)
```

```
## Linear mixed model fit by REML. t-tests use Satterthwaite's method [
## lmerModLmerTest]
## Formula: rAUC_fil ~ mean_rAUC_si_phage + log10(pfu) + (1 | phage_pop_id)
##    Data: data_sum_merged_phage
## 
## REML criterion at convergence: -16.4
## 
## Scaled residuals: 
##     Min      1Q  Median      3Q     Max 
## -1.2763 -0.6584 -0.2204  0.8262  1.1718 
## 
## Random effects:
##  Groups       Name        Variance Std.Dev.
##  phage_pop_id (Intercept) 0.000000 0.00000 
##  Residual                 0.003347 0.05785 
## Number of obs: 10, groups:  phage_pop_id, 4
## 
## Fixed effects:
##                     Estimate Std. Error        df t value Pr(>|t|)    
## (Intercept)         0.057147   0.179815  7.000000   0.318 0.759900    
## mean_rAUC_si_phage  0.760805   0.138036  7.000000   5.512 0.000895 ***
## log10(pfu)         -0.008875   0.014745  7.000000  -0.602 0.566218    
## ---
## Signif. codes:  0 '***' 0.001 '**' 0.01 '*' 0.05 '.' 0.1 ' ' 1
## 
## Correlation of Fixed Effects:
##             (Intr) m_AUC_
## mn_rAUC_s_p -0.851       
## log10(pfu)  -0.874  0.504
## optimizer (nloptwrap) convergence code: 0 (OK)
## boundary (singular) fit: see help('isSingular')
```

###### Summary of Model 3.2

1. rAUC of MPAO1 in bulk filtrate increases with increasing rAUC of
   MPAO1 in co-culture from isolates from the same filtrate.

#### Figure 5

The following model relate to results found in Figure 5.

Model 5.1: Twitch motility of ancestral, mutant, and endpoint
isolates.

###### Importing data

```
iso_twitch <- read.csv("~/Documents/PhD Folder/Research/2025 Spring/NCEE - Prophage Paper/ncee_prophage-paper_data/bac_iso_twitch/ncee_iso_twitch.csv")

#Renaming strain
iso_twitch$si_pf_pres <- replace(iso_twitch$si_pf_pres, iso_twitch$si_pf_pres=="PAO1", "MPAO1")

#Renaming strain
iso_twitch$si_pf_pres <- replace(iso_twitch$si_pf_pres, iso_twitch$si_pf_pres=="PAO1_pilA", "MPAO1ΔpilA")

#Reordering levels
iso_twitch$si_pf_pres <- factor(iso_twitch$si_pf_pres, levels=c("MPAO1", "LacZ", "MPAO1ΔpilA", "Pf+ Iso", "Pf- Iso"))
```

###### Generating Model 5.1

Model 5.1 compares the twitch motility of ancestral, mutant, and
endpoint isolates.

```
#Generating Model 5.1
mod5.1 <- lmer(data=iso_twitch, sqrt(area) ~ si_pf_pres + (1|date))

#Testing for overall effect of phage category (MPAO1, LacZ, MPAO1ΔpilA, Pf+ iso, Pf- iso)
Anova(mod5.1, type="III")
```

```
## Analysis of Deviance Table (Type III Wald chisquare tests)
## 
## Response: sqrt(area)
##              Chisq Df Pr(>Chisq)    
## (Intercept) 435.44  1  < 2.2e-16 ***
## si_pf_pres  433.48  4  < 2.2e-16 ***
## ---
## Signif. codes:  0 '***' 0.001 '**' 0.01 '*' 0.05 '.' 0.1 ' ' 1
```

```
#Testing which phage categories significantly differed in twitch motility
pairs(emmeans(mod5.1, ~si_pf_pres))
```

```
##  contrast                estimate     SE  df t.ratio p.value
##  MPAO1 - LacZ            0.348419 0.0526 246   6.624 <0.0001
##  MPAO1 - MPAO1ΔpilA      0.607791 0.0526 246  11.556 <0.0001
##  MPAO1 - (Pf+ Iso)       0.612947 0.0376 246  16.301 <0.0001
##  MPAO1 - (Pf- Iso)       0.349094 0.0429 246   8.129 <0.0001
##  LacZ - MPAO1ΔpilA       0.259372 0.0526 246   4.931 <0.0001
##  LacZ - (Pf+ Iso)        0.264527 0.0376 246   7.035 <0.0001
##  LacZ - (Pf- Iso)        0.000674 0.0429 246   0.016  1.0000
##  MPAO1ΔpilA - (Pf+ Iso)  0.005155 0.0376 246   0.137  0.9999
##  MPAO1ΔpilA - (Pf- Iso) -0.258698 0.0429 246  -6.024 <0.0001
##  (Pf+ Iso) - (Pf- Iso)  -0.263853 0.0222 246 -11.898 <0.0001
## 
## Note: contrasts are still on the sqrt scale. Consider using
##       regrid() if you want contrasts of back-transformed estimates. 
## Degrees-of-freedom method: kenward-roger 
## P value adjustment: tukey method for comparing a family of 5 estimates
```

###### Summarizing Model 5.1

1. twitch motility varied across isolate category (MPAO1, LacZ,
   MPAO1ΔpilA, Pf+ iso, Pf- iso)
2. For twitch motility, MPAO1 > LacZ = Pf- iso > Pf+ iso =
   MPAO1ΔpilA.
