## Supplementary Methods for "Mutations in filamentous bacteriophages spark eco-evolutionary feedbacks in *Pseudomonas aeruginosa*"

**Short Title**: Prophage mutations re-route evolution

**Authors**: Noah S. B. Houpt^1,*^, Catherine A. Hernandez^1,2,3^, Paul E. Turner^1,4,5,6^

**Keywords**: Niche construction, lysogenic bacteriophage, prophage, type IV pilus, motility, Pf4, Pf6, phage therapy, virulence evolution

^1^Department of Ecology & Evolutionary Biology, Yale University, New Haven, CT, USA

^2^Yale Institute for Biospheric Studies, Yale University, New Haven, CT, USA

^3^Department of Biological Sciences, University of South Carolina, Columbia, SC, USA

^4^Center for Phage Biology and Therapy, Yale University, New Haven, CT, USA

^5^Yale Quantitative Biology Institute, Yale University, New Haven, CT, USA

^6^Program in Microbiology, Yale School of Medicine, New Haven, CT, USA

*Supplementary Methods*

*Evolution experiment*

16 populations of *Pa* MPAO1 were experimentally evolved at either high (HD) or low (LD) population density for 1,000 generations. Half of the populations in each treatment were founded by a strain with a chromosomally inserted *lacZ* gene (MPAO1-*lacZ*, LacZ for short). This let us distinguish this strain from wild-type MPAO1 on agar plates supplemented with bromochloroindoxyl galactoside (X-Gal) where LacZ forms blue colonies, allowing us to detect cross-contamination between populations. The 8 wild-type MPAO1 and LacZ populations were founded from a single colony of each strain. All populations were grown in M9 media (22 mM KH_2_PO_4_, 48 mM Na_2_HPO_4_•H_2_O, 8.6 mM NaCl, 19 mM NH_4_Cl, 1 mM MgSO_4_, 100 µM CaCl_2_, 18 µM FeSO_4_) supplemented with 37 mM sodium succinate as the sole carbon source. Populations were grown in 1.5 mL cultures in 24 well tissue culture plates (CellTreat, #229524) wrapped in petri dish bags to minimize evaporation. Each day, populations were grown in well-mixed cultures (200 rpm) at 30 °C for 23 h before being transferred to fresh media (1 h transfer time). Aliquots of each population were stored at −80 °C in 20% glycerol every 250 generations.

The aim of our design was to manipulate population density while maintaining the daily population bottleneck and culture vessel consistent across treatments. To this end, population density was manipulated by varying the dilution regime across treatments. HD lines were grown in a single well of a 24 well plate and diluted 1:100 each day into fresh media (6.64 generations/day). Conversely, LD lines were split across 23 wells, grown for 23 h, and then combined and diluted 1:2300 into 23 wells filled with fresh media daily (11.17 generations/day). This resulted in an equivalent population bottleneck across treatments (~10^8^ cells), but a 23-fold lower starting density in LD lines each day. Consequently, LD populations experienced lower average population density during each growth cycle (Figure 1B). Each HD population was paired with an LD population on the same plate (e.g. HD1 paired with LD1). Paired populations were founded by oppositely marked ancestral strains such that cross contamination could be easily detected on plates supplemented with X-Gal. End-of-passage population density was measured every 2-7 passages by plating an aliquot of leftover culture from each population following its transfer to fresh media. Aliquots were diluted in M9 media and plated on M9 + succinate agar plates (1.5% agar) with 20 µg/mL X-Gal. Due to the different number of bacterial generations experienced each day across treatments, HD and LD lines were passaged for a different number of days (HD: 151 days, LD: 90 days).

*Microplate growth curves in filtrate*

To explore the evolution of organism-driven environmental change in our experiment, ancestral and endpoint bacteria were grown in microplate growth curves in the presence or absence of filtrate generated by bacteria sampled across evolutionary timepoints. Filtrate was generated by first reviving populations from frozen stocks in a 23-h culture. We then transferred revived populations to fresh media according to their evolved density treatment for another 23-h growth period. 1 mL from each culture was then transferred to a sterile 1.5 mL Eppendorf tube and centrifuged at 10,000 g for 10 min. The supernatant from each culture was then filtered using a syringe-tip 0.22 µm PDVF filter (Millex, SLGVR33R5) into a fresh Eppendorf tube, generating the cell-free filtrate. This filtrate should include cellular products smaller than the pore size, including any metabolites, toxins, extracellular proteins, or bacteriophages released by cells during growth or cell death.

The pellets of ancestral and endpoint populations were then resuspended in 1 mL of 1x M9 salt solution. To minimize carryover of spent media from the previous day’s culture, the resuspended bacteria was centrifuged, the supernatant was discarded, and pellets were resuspended again in 1 mL 1x M9 salt solution.

Growth curves were run in flat-bottom 96 well plates (Falcon, #351172). Curves in unmodified media were run in wells with 190 µL of 1x media and 10 µL of bacterial culture standardized to OD_600_ = 0.5 (starting density OD_600_ = 0.025). Conversely, curves in media with filtrate were run in wells composed of 2/3 media (126.7 µL) with 1.5x the normal concentration of succinate (55.5 mM sodium succinate), and 1/3 filtrate (63.3 µL). Concentrated media was used to ensure that the same amount of non-filtrate-derived carbon was supplied in wells with and without filtrate. Growth curves were run for 23 h at 30 °C with constant double-orbital shaking and optical density (OD_600_) was measured every 10 min in a BioTek Epoch 2 plate reader.

Performance of ancestral and endpoint populations in filtrate was used to quantify how niche construction varied over time and the extent to which endpoint populations adapted to organism-modified environments. The area-under-the-curve (AUC) was used to measure performance in each condition. The AUC was extracted from growth curves using the R package *gcplyr* (70 min bin widths used for computing moving average of optical density; Blazanin, 2024). The impact of filtrate on a population was quantified using the quotient of its AUC in the presence vs. absence of filtrate, which we called the relative AUC, or rAUC (1).

$${rAUC}_{p,f}= \frac{{AUC}_{p,f}}{{AUC}_{p,nf}} (1)$$

Where *AUC_p,f_* represents the AUC of population *p* in filtrate *f* and *AUC_p,nf_* represents the AUC of population *p* in the absence of filtrate. rAUC values exceeding 1 indicate filtrate benefited the population, whereas values below 1 indicate inhibition.

To quantify the extent to which endpoint populations were adapted to filtrate, the quotient of rAUC values from endpoint populations and ancestral populations was calculated, which we called the relative difference in AUC, or rdAUC (2).

$${rdAUC}_{p,f}= \frac{{rAUC}_{p\_end,f}}{{rAUC}_{p\_anc,f}} (2)$$

Where *rAUC_p_end,f_* represents the relative performance of the endpoint population in filtrate *f* and *rAUC_p_anc,f_* represents the relative performance of the ancestral population in filtrate *f*. rdAUC values exceeding 1 indicate that the endpoint population performed comparatively well in filtrate compared to unmodified media, relative to its ancestor.

*Phage quantification*

The density of free hyperactive phage in filtrate was measured by plating 10-fold dilution series of filtrate on lawns of MPAO1. Lawns were generated by adding 200 µL of an overnight culture of MPAO1 to 4 mL of lysogeny broth (LB) top agar (4 g/L agar) and spreading it on a LB agar plate (15 g/L agar). Filtrate was plated at dilution factors ranging from 10^0^-10^7^ in 2 µL spots in duplicate across each of two days (n = 4). Filtrate phage density was estimated by counting the number of plaques at the lowest dilution level at which individual plaques could be resolved.

*Phage removal/addition experiment*

To investigate whether hyperactive phage were the cause of inhibitory filtrate, hyperactive phage was experimentally removed from evolved filtrate or added phage into ancestral filtrate and measured the impact on MPAO1 in microplate growth curves. For each experiment, a subset of inhibitory filtrates was used, focussing on filtrates generated by two populations from each treatment that varied in the extent the which they inhibited MPAO1 in microplate growth curves. Specifically, all hyperactive phage-containing filtrates from populations HD1 (generation 500, 750) and LD6 (generation 250, 500), which weakly inhibited MPAO1, and HD7 (generation 250, 500, 750, 1000) and LD8 (generation 250, 500), which strongly inhibited MPAO1, were used. Filtrate was generated in the same way as described above, except here four replicate 23-h cultures were pooled before initial centrifugation to ensure enough filtrate was produced (pooled culture spun in 15 mL conical tube at 3,220 rpm for 10 min before filtration).

In the phage removal experiment, whether phage presence was necessary to explain the inhibitory effect of phage-containing filtrates on MPAO1 was tested. Phage were experimentally removed from filtrate using 4 mL cellulose centrifugal filters with a pore size of 10 kDa (centrifuged at 3,220 rpm for 10 min; Amicon Ultra 4, Millipore, UFC9010). Then, phage titers in filtrates post-centrifugal filtration was measured by plating dilutions of the modified filtrate on lawns of MPAO1, as above. Centrifugal filtration successfully removed phage in all but one replicate, which was consequently not considered in our analysis (Supplementary Figure 2). In this experiment, MPAO1 was grown in the presence of unmodified ancestral or hyperactive phage-containing filtrate, or the same filtrates after centrifugal filtration.

In the phage addition experiment, whether phage were sufficient to explain the inhibitory effect of phage-containing filtrates on MPAO1 was tested. This was done by adding a mix of three phage isolates collected from each inhibitory filtrate into ancestral filtrate. Phage isolates were collected by first plating filtrate on lawns of MPAO1. Then three random plaques from each filtrate was transferred into separate Eppendorf tubes containing 1 mL M9 media from which they were diluted and plated on fresh MPAO1 lawns. This was repeated three times to ensure each phage isolate was isogenic. Phage isolate stocks were generated by transferring a single plaque of each isolate into separate cultures of MPAO1, which were grown for 6 h before centrifugation and filtration as described above. Phage isolate stocks were titered as above. Phage isolates from each filtrate were mixed in equal proportion to generate phage isolate mixes, which were spiked into ancestral filtrate at densities approximating the densities detected in bulk filtrates (50 µL of 10x concentrated phage mix added to 450 µL of ancestral filtrate). To control for the small dilution caused by adding phage into filtrate, all other filtrates across both experiments were similarly diluted with M9 media. Phage densities were confirmed to be similar in evolved filtrates and in anc + Φ filtrates (Supplementary Figure 2). In this experiment, MPAO1 was grown in the presence of unmodified ancestral or phage-containing evolved filtrate (same curves as above) or ancestral filtrate with phage spiked-in. Microplate growth curves were carried out and analyzed as described above.

*Phage isolate sequencing and genomic analysis*

To identify the mutations responsible for the emergence of hyperactive phages, sequencing was attempted for the 30 phage isolates used in the phage addition experiment (3 isolates from each of ten filtrates from 4 total populations). Phage isolate stocks were prepared for sequencing by amplifying 20 µL of master phage stocks in 10 mL cultures of MPAO1 overnight. Samples were then centrifuged and filtered as described above and 6 mL of the filtrate was added to fresh 15 mL conical tubes with 1.5 mL of 5x phage precipitation solution (20% w/v PEG 8000, 2.5 M NaCl). Tubes were mixed and stored at 4 °C overnight. The next morning, tubes were centrifuged for 30 min at 3,220 rpm to pellet the precipitated phage. Pellets were resuspended in 360 µL of phage resuspension buffer (1M NaCl, 10 mM Tris•HCl (pH 7.6), 0.1 mM EDTA) and transferred to fresh Eppendorf tubes. Next, to degrade any host DNA or RNA present in the resuspension, 40 µL of 10x DNAse I buffer (New England Biolabs, B0303SVIAL), and 1 µL each of DNAse I (2,000 U/mL, New England Biolabs, M0303SVIAL) and RNAse A (20 mg/mL, New England Biolabs, T3018L) was added to each tube, which was then incubated for 30 min at 37 °C. Then, 10 µL of 0.1 M EDTA (pH 8.0) was added to chelate divalent metals.

Phenol chloroform DNA extractions were then performed following an established protocol (https://barricklab.org/twiki/bin/view/Lab/ProtocolsPhageGenomicDNA). 500 µL of phenol:chloroform:isoamyl alcohol (25:24:1) was added to each sample, which was then mixed by inversion for ~15 s and transferred into fresh tubes. Mixtures were centrifuged for 5 min at 16,000 g, and the aqueous phase was transferred into fresh tubes (~450 µL). Then, 50 µL of 3 M sodium acetate was added, followed by 1,000 µL of 100% ethanol prechilled to −20 °C. Samples were then incubated at −20 °C overnight, allowing DNA to precipitate from the solution.

After the overnight incubation (~18 hr), each sample was centrifuged for 15 min at 14,000 g and the supernatant was removed. DNA pellets were dislodged in 500 µL of 70% ethanol prechilled to −20 °C before samples were centrifuged again for 5 min at 14,000 g. The supernatant was then removed and the pellets were left to air dry before being resuspended in 200 µL of 10 mM Tris•HCl (pH 7.6). Extracted DNA was then purified using a DNA purification kit (Zymo Research DNA Clean & Concentrator Kit, D4013) and ssDNA concentrations were measured via spectroscopy (Nanopore One, Thermo Scientific). insufficient DNA was extracted from three phage isolates over two attempts, all derived from line HD1 (HD1_0500_ Φ2, HD1_0500_ Φ3, HD1_0750_ Φ3), and so these phages could not be sequenced. Extracted DNA was sent to Plasmidsaurus for Oxford Nanopore sequencing using their small linear dsDNA sequencing package. Despite having ssDNA genomes, this approach provided high quality reads with suitable coverage for all isolates (≥ 100-fold coverage of phage genomes). Raw sequencing reads are available on the NCBI Sequence Read Archive (PRJNA1420075).

Raw reads were filtered for quality using *fastplong* (mean base quality > Q20; Chen et al., 2018) and filtered reads were used to call variants relative to reference genomes of both known prophages present in MPAO1 (Pf4 and Pf6). Prophage reference genomes were generated by first extracting the known sequence of these phages from the reference genomes of PAO1 (for Pf4; NCBI Reference Sequence: NC_002516.2) and MPAO1 (for Pf6; NCBI Reference Sequence: NZ_CP027857.1). Prophage reference genomes were annotated using *pharokka* (using both *prodigal* and *phannotate* to call CDS’; Bouras et al., 2023) to identify coding regions and then gene names, locus IDs, and gene functions were manually adjusted based on relevant literature (Schmidt et al., 2022; Guo et al., 2024). Variants in the phage isolates relative to both reference genomes were called using *breseq* in nanopore mode (Deatherage and Barrick, 2014). Variants were classified as core genome mutations if they occurred within the region of homology shared by Pf4 (*pf4r* to *intF4*) and Pf6 (*pf6r* to *intF6*). Variants outside of this region were classified as accessory genome mutations. The proportion of reads mapping to either reference, as well as the number of variants was used to judge whether each phage isolate was Pf4- or Pf6-derived.

For two phage isolates, a high number of variants were detected when reads were compared to either reference genome. This led us to hypothesize that these isolates were recombinants of Pf4 and Pf6. This possibility was further investigated by mapping the raw reads from four phage isolates (two putative recombinants, two non-recombinants) to Pf4 and Pf6 simultaneously using the *bwa* package (Li, 2013). To test for recombination, regions mapping to one or the other reference phage genome were identified in the raw reads from a single phage isolate. The location of SNPs within phage isolate genomes were then visually compared to locations of apparent recombination to examine whether they overlapped, which would indicate that SNPs were called as a consequence of recombination.

To visualize phage isolate genomes, *de novo* assemblies of each isolate were generated using *flye* (Kolmogorov et al., 2019). Assemblies were manually reoriented in SnapGene by setting the orientation and starting sequence to be the same as the phage from which each isolate was derived. Coding regions were predicted using *pharokka* (using both *prodigal* and *phannotate* to call CDS’; Bouras et al., 2023) and manually annotated based on either the Pf4 or Pf6 reference genomes.

*Phage virulence measurement*

The virulence of our phage isolates was quantified using microplate growth curves. The rAUC of MPAO1 in the presence vs. absence of a phage isolate was used as a measure of virulence (as virulence increases, rAUC decreases). Each growth curve well consisted of 188 µL of media, 10 µL of MPAO1 culture adjusted to OD_600_ = 0.5 (starting OD_600_ = 0.025), and 2 µL of phage isolate stock standardized to 10^7^ PFU/mL (multiplicity of infection ~ 0.01). In no-phage control wells, 2 µL of M9 media was added instead of phage stock. Growth curves were programmed and analyzed as described above.

*Endpoint bacterial population sequencing*

Endpoint bacterial populations and ancestral strains were revived from glycerol stocks in 23 h cultures. They were then transferred to fresh media using dilution factors corresponding to their evolved treatment and grown for an additional 23 hr. The second growth step was intended to minimize any shifts in allele frequencies resulting from growth in the presence of glycerol from the frozen stocks. The genomic DNA was then extracted from these cultures using the protocol for Gram negative bacteria of a DNeasy Blood and Tissue DNA extraction kit (Qiagen, 69504). Remaining culture was diluted and plated on X-Gal plates to confirm there was no cross-contamination between oppositely marked populations. Short-read sequencing on extracted DNA was performed by SeqCenter on a NovaSeq X Plus Sequencer (Illumina), producing 2 x 151 bp paired-end reads. Raw sequencing reads are available on the NCBI Sequence Read Archive (PRJNA1420075).

*fastp* (Chen et al., 2018) was used to filter raw reads for quality using default parameters (Q >= 15, % unqualified bases >= 40%, minimum read length = 15 bp) and downsample to appropriate sequencing depth for analysis (downsampled to ~250-fold coverage for populations, ~50-fold coverage for ancestors). Then, variants in the sequenced populations relative to the MPAO1 reference genome (NCBI Reference Sequence: NZ_CP027857.1) were called using *breseq* in polymorphism mode (frequency cutoff = 0.05). Summary data for all variants in endpoint populations was then filtered for variants in genes related to type IV pilus biosynthesis and function.

*Generating endpoint isolate library*

To investigate the phenotypic consequences of hyperactive phage emergence, a library of three isolates from each endpoint population was generated. Isolates were randomly selected from separate colonies generated by plating frozen stocks of endpoint populations on LB agar plates. Isolates were re-streaked three times to ensure each represented a single genotype. Stocks of each isolate were generated by growing them overnight and storing them in 20% glycerol at −80 °C.

*Twitch assay*

To measure twitch motility in endpoint bacterial isolates, streak plates of our endpoint isolate library as well as both ancestral strains and MPAO1ΔpilA were generated. MPAO1ΔpilA lacks the gene encoding the major pilin of the TIVP and is incapable of twitch motility, making it an appropriate negative control for this assay. Each strain was grown in a 1.5 mL LB liquid culture from a single isolated colony for 23 hr. Subsequently, sterile wooden toothpicks were used to inoculate each strain into the center of a freshly poured LB plate (10 g/L agar). Toothpicks were stirred in each bacterial culture and then pushed to the bottom of each agar plate ~5 times, making contact with the plastic petri dish bottom each time. Plates were then incubated upside down for 48 h at 37 °C. Following incubation, the agar was removed from each plate using a sterile wooden stick and then stained the bottom of each plate using 0.1% w/v crystal violet. Plates were then scanned using an Epson Perfection V850 Pro scanner. Scans were then uploaded to ImageJ and manually outlined the stained region, which we used to measure the twitch area. The twitch area of each strain was then measured in a single technical replicate on each of 5 days (n = 5).

*Virulent phage susceptibility assay*

To measure the susceptibility of endpoint bacterial isolates to virulent (ie. obligately lytic) phages, working stocks of two virulent phages, LPS-5 and ΦKMV, were first generated. Then, each phage strain was plated on a lawn of MPAO1 as described above. From these lawns, a single plaque was randomly selected and amplified in a 10 mL culture of MPAO1 in LB for 6 hr. These phage amplifications were then centrifuged (3,220 rpm for 10 min) and the resulting supernatant was filtered using 0.22 µm syringe filters as described above. The titer of these stocks was then adjusted to 10^10^ PFU/mL by dilution with fresh LB.

Lawns of each endpoint isolate, both ancestors, MPAO1ΔpilA, and MPAO1 algC::Tn were generated. MPAO1 algC::Tn has a transposon inserted into the *algC* gene of MPAO1 and is consequently resistant to many lipopolysaccharide (LPS)-targeting phages, including LPS-5. Thus, it acted as a negative control in our phage susceptibility assay for LPS-5, while MPAO1ΔpilA acted as a negative control for ΦKMV, which targets the TIVP (Lavigne et al., 2003). Overnight cultures of each strain were grown from colonies in LB at 37 °C. These conditions were used to simulate clinical testing of phage susceptibility in *Pa* isolates. Lawns of each strain were generated as described above, but with 7.5 g/L LB top agar to reduce the size of plaques. Both virulent phage stocks were plated at dilution factors ranging from 10^0^-10^7^ in 10 µL spots. The relative number of plaques observed on each endpoint isolate or control strain compared with its relevant reference strain (MPAO1 or LacZ) was used to compute the efficiency of plating (EOP) of each phage (3):

${EOP}_{p,s}= \frac{{Estimated phage titer}_{p,s}}{{Estimated phage titer}_{p,r}}$	(3)

Where *p* represents the phage ID (LPS-5 or ΦKMV), *s* represents the focal bacterial strain, and *r* represents the appropriate reference strain for *s*.

*Statistics*

All statistical analyses were performed in R (version 4.2.0). Our modeling approach was to use linear mixed-effects models (*lme4* package; Bates et al., 2015) to compare phenotypes across groups while controlling for factors like date and population ID using random effects where appropriate. Type III ANOVAs were used for overall significance testing (type III Wald χ² tests) of main effects and post-hoc contrasts of relevant groups were carried out using the *emmeans* package. Where appropriate, we used Bonferroni corrections were used to adjust the P-values from post-hoc contracts. Please see Supplementary File 1 for more detailed information on the specific statistical models used. All non-sequencing data used in this study are included in supplementary files associated with this article.
