## Supplementary Figures for "Mutations in filamentous bacteriophages spark eco-evolutionary feedbacks in *Pseudomonas aeruginosa*"

**Short Title**: Prophage mutations re-route evolution

**Authors**: Noah S. B. Houpt^1,*^, Catherine A. Hernandez^1,2,3^, Paul E. Turner^1,4,5,6^

**Keywords**: Niche construction, lysogenic bacteriophage, prophage, type IV pilus, motility, Pf4, Pf6, phage therapy, virulence evolution

^1^Department of Ecology & Evolutionary Biology, Yale University, New Haven, CT, USA

^2^Yale Institute for Biospheric Studies, Yale University, New Haven, CT, USA

^3^Department of Biological Sciences, University of South Carolina, Columbia, SC, USA

^4^Center for Phage Biology and Therapy, Yale University, New Haven, CT, USA

^5^Yale Quantitative Biology Institute, Yale University, New Haven, CT, USA

^6^Program in Microbiology, Yale School of Medicine, New Haven, CT, USA

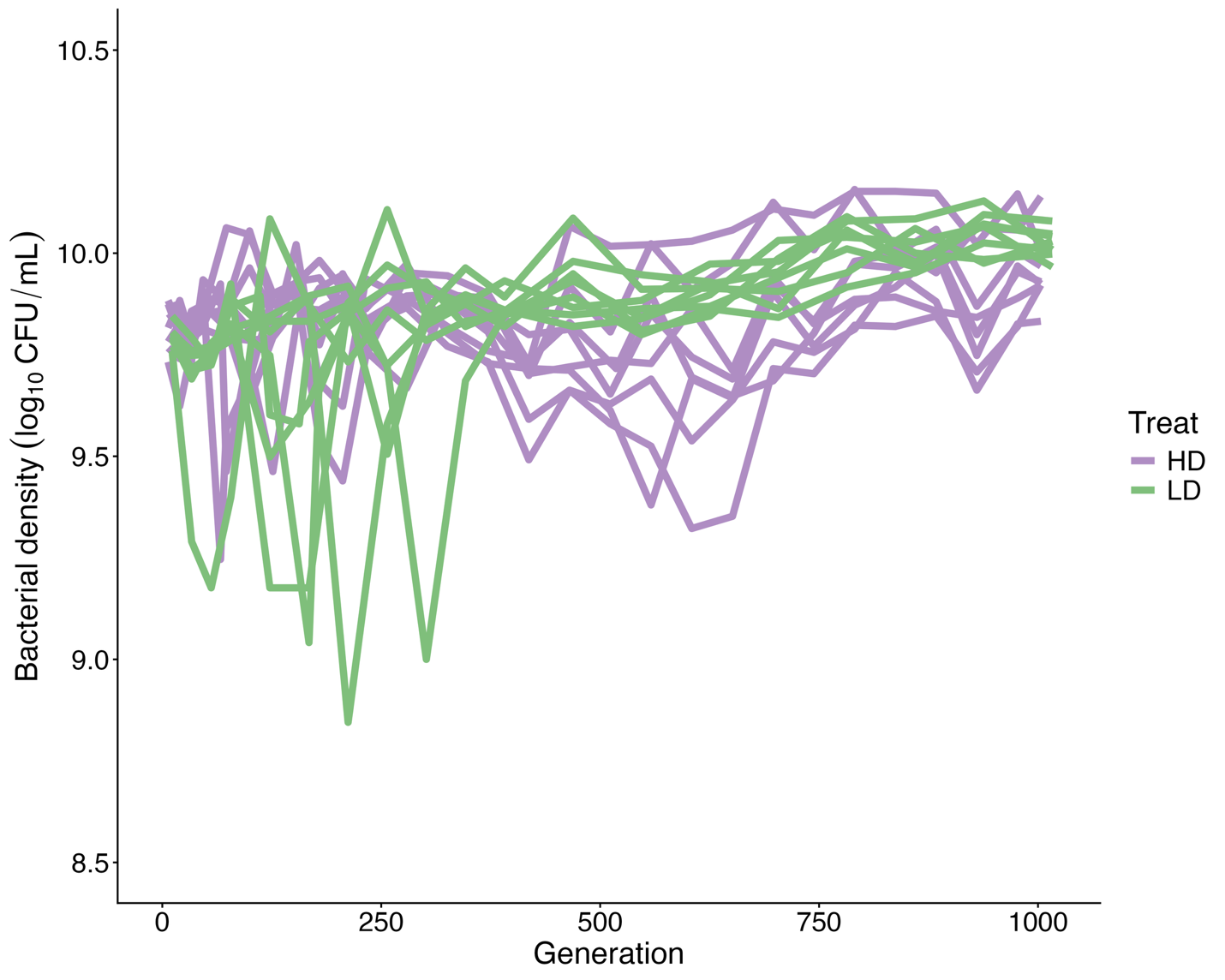

**Supplementary Figure 1:** **End-of-passage population density crashes and recovers in most HD and LD lines.** End-of-day passage density of high density (HD) and low density (LD) populations over generations (points on each line represent the average of two technical replicates). Each line represents one experimental lineage.

Alt text: Line graph visualizing the end-of-passage population densities of HD and LD lines.

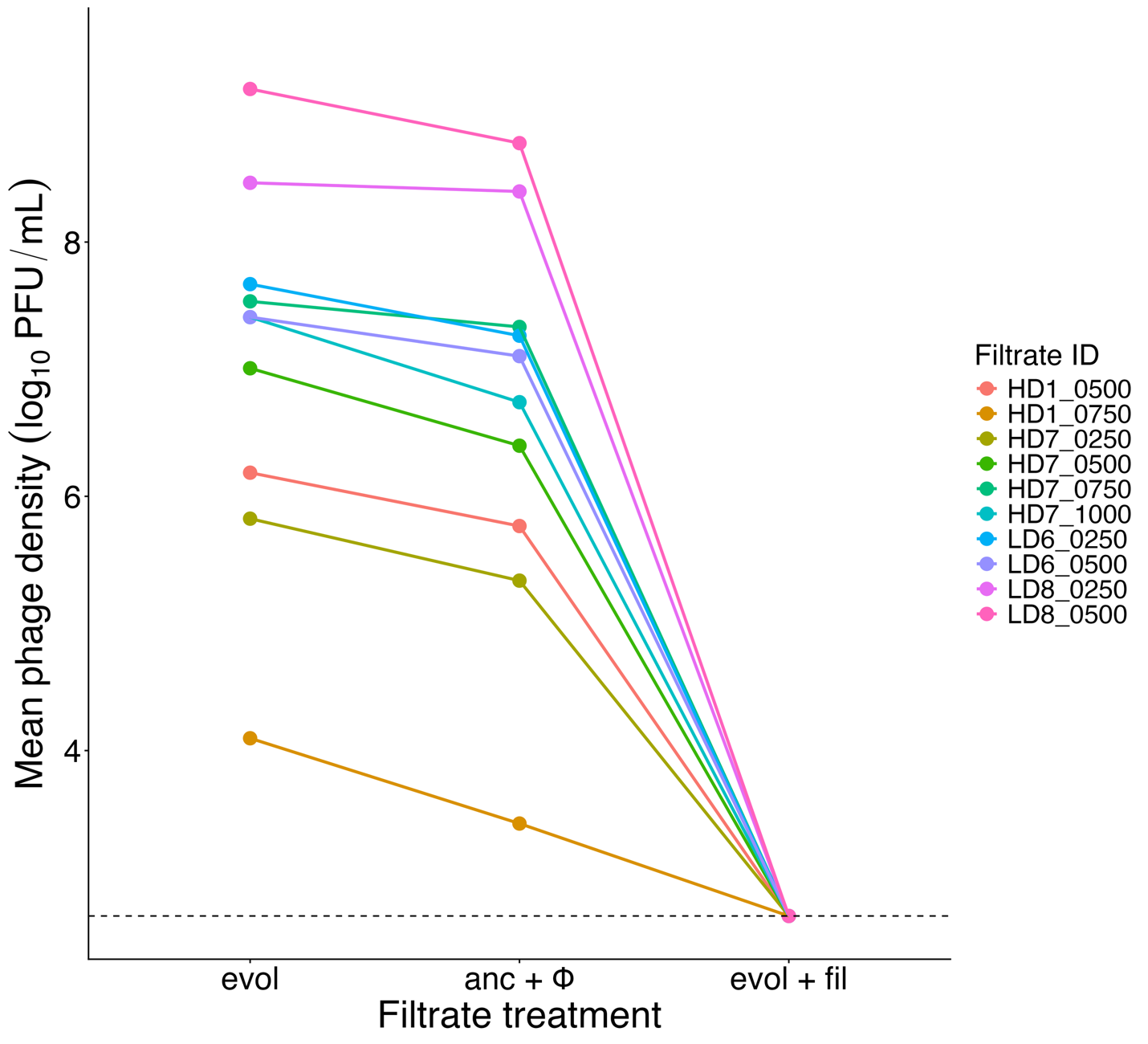

**Supplementary Figure 2: Centrifugal filters successfully removed virulent phage from filtrates.** Mean phage density of evolved filtrate (evol), ancestral filtrate with contemporaneous phage isolates spiked in (anc + Φ), and evolved filtrate that has been filtered through a 10 kDa centrifugal filter (evol + fil). Each point represents the mean of three replicates. Dashed line indicates the lower limit of detection.

Alt text: Line graph visualizing phage titers in bulk evolved filtrate, ancestral filtrate with phage isolates spiked-in, and evolved bulk filtrate following centrifugal filtration.

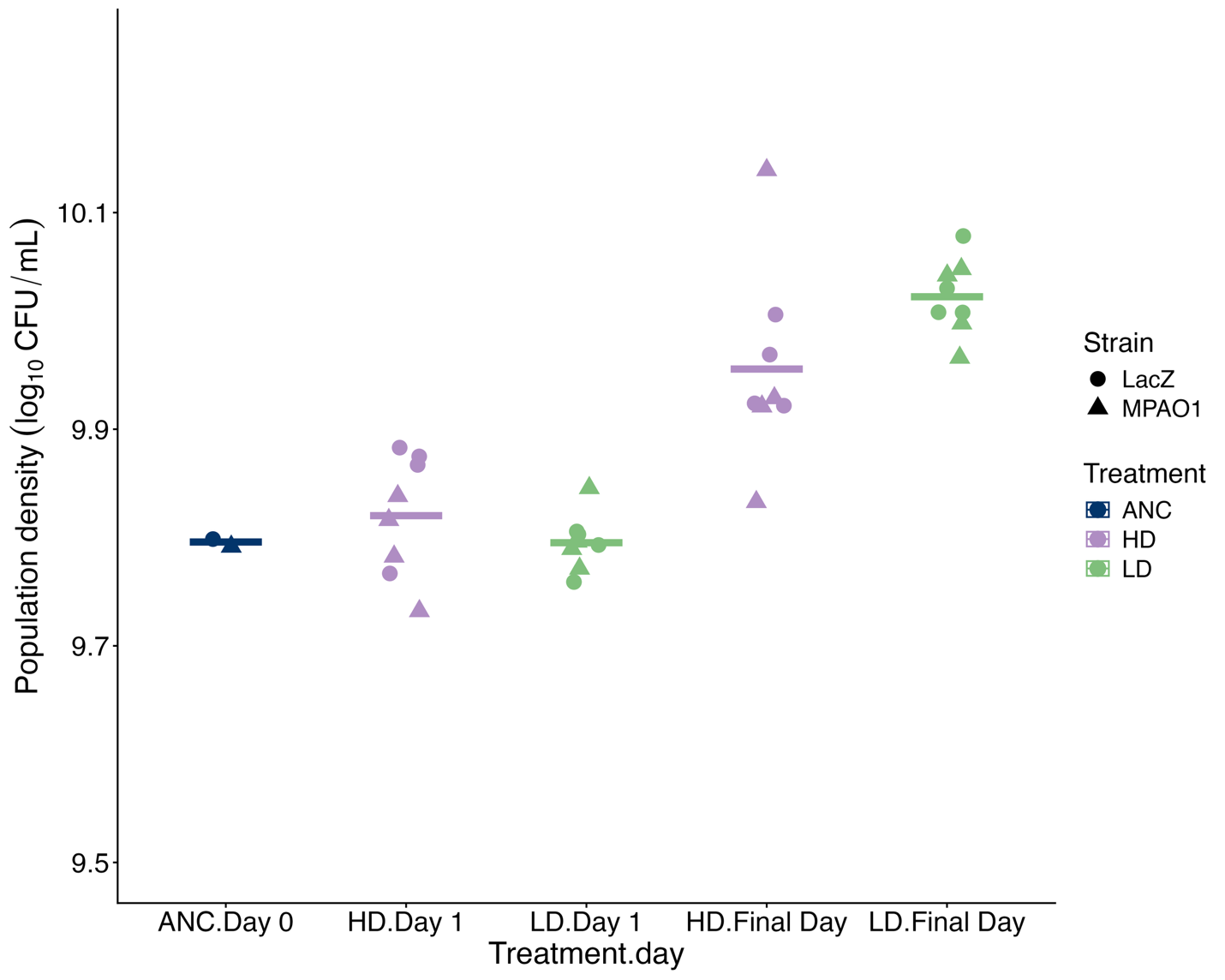

**Supplementary Figure 3: End-of-passage population density increases in both treatments over the evolution experiment.** Each point represents the mean of two technical replicates for each population (n = 2 for ANC, representing marked and unmarked ancestral strains; n = 8 for HD and LD populations). Horizontal bars show mean population density within groups. Population density of two ancestral cultures that initiated experimental populations was measured on “Day 0”. Final day was different across treatments (HD: Day 151, LD: Day 91) due to variation in the number of generations elapsed per day in HD and LD lines (~6.64 and ~11.17 gens/day, respectively).

Alt text: Jitter plot of end-of-passage population density of ancestral bacteria and HD and LD populations following the first and final passage.

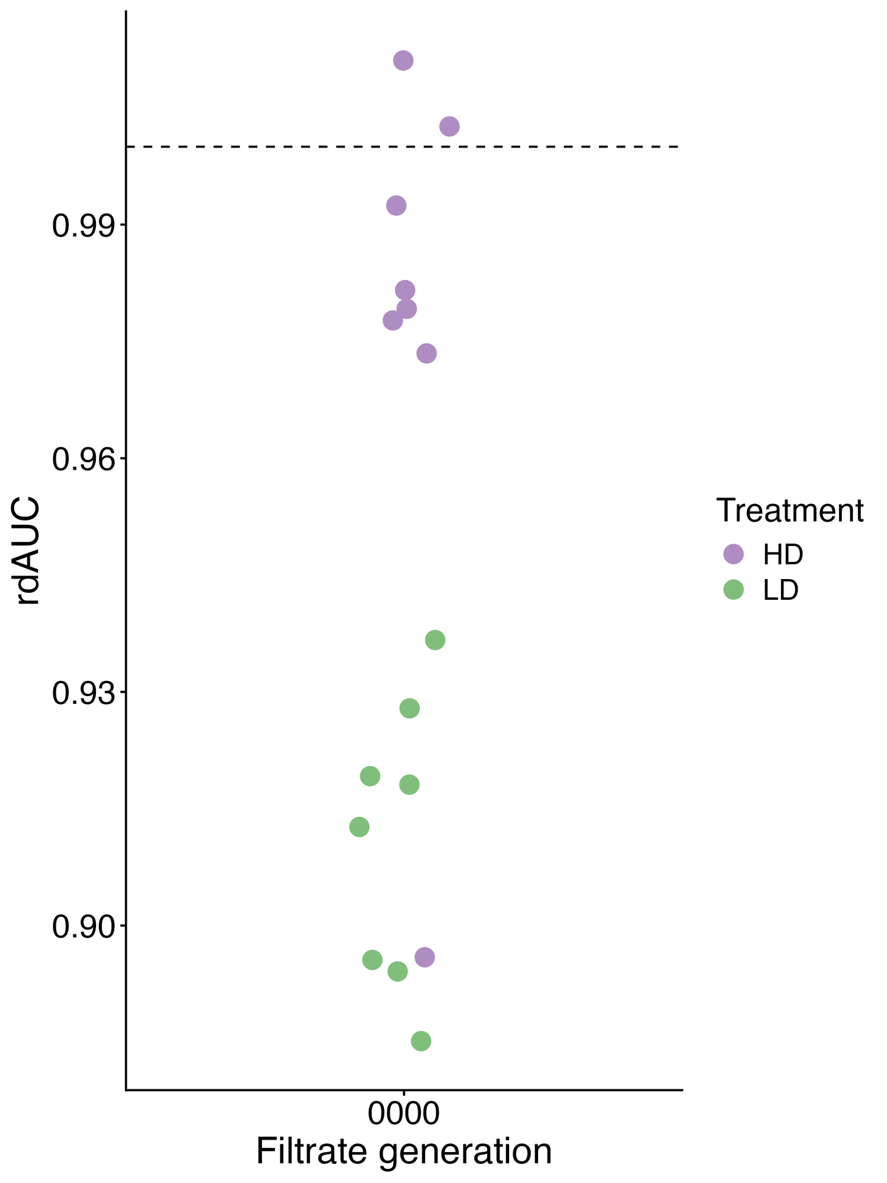

**Supplementary Figure 4: Endpoint populations do not have a relative advantage in ancestral filtrate.** Each point represents the average rdAUC value for one endpoint population. rdAUC values are calculated using the area-under-the-curve of 16 replicate growth curves (four replicates each of ancestral and endpoint populations grown in the presence or absence of the focal filtrate). Upper dashed line denotes an rdAUC value of 1, which would reflect filtrate having an equal effect on ancestral and endpoint bacteria.

Alt text: Jitter plot of rdAUC of HD and LD endpoint populations in ancestral filtrate.

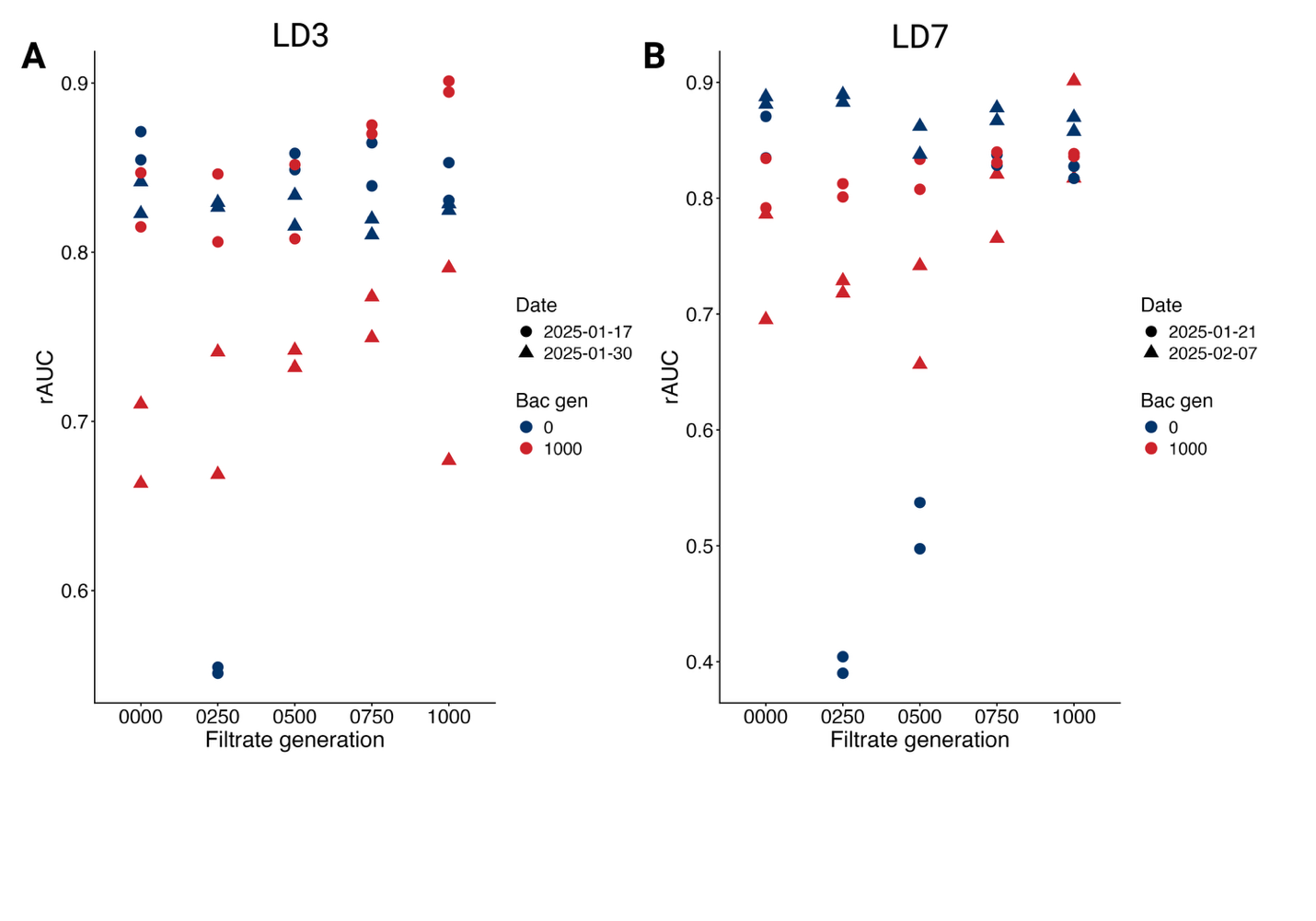

**Supplementary Figure 5: Effect of filtrate on ancestor strongly varies for some evolutionary timepoints across replicates.** Ancestor is strongly inhibited by filtrate prepared on one of two replicate days from **A**: generation 250 and **B**: generation 250 and 500 from lines LD3 and LD7, respectively. Created in BioRender. Turner, P. (2026) https://BioRender.com/tfe6eg9

Alt text: Two graphs showing the rAUC of ancestral and endpoint bacteria grown in the presence of various filtrates over time for populations LD3 and LD7. For each populations, ancestral bacteria are strongly inhibited by certain filtrates in some replicates but not others.

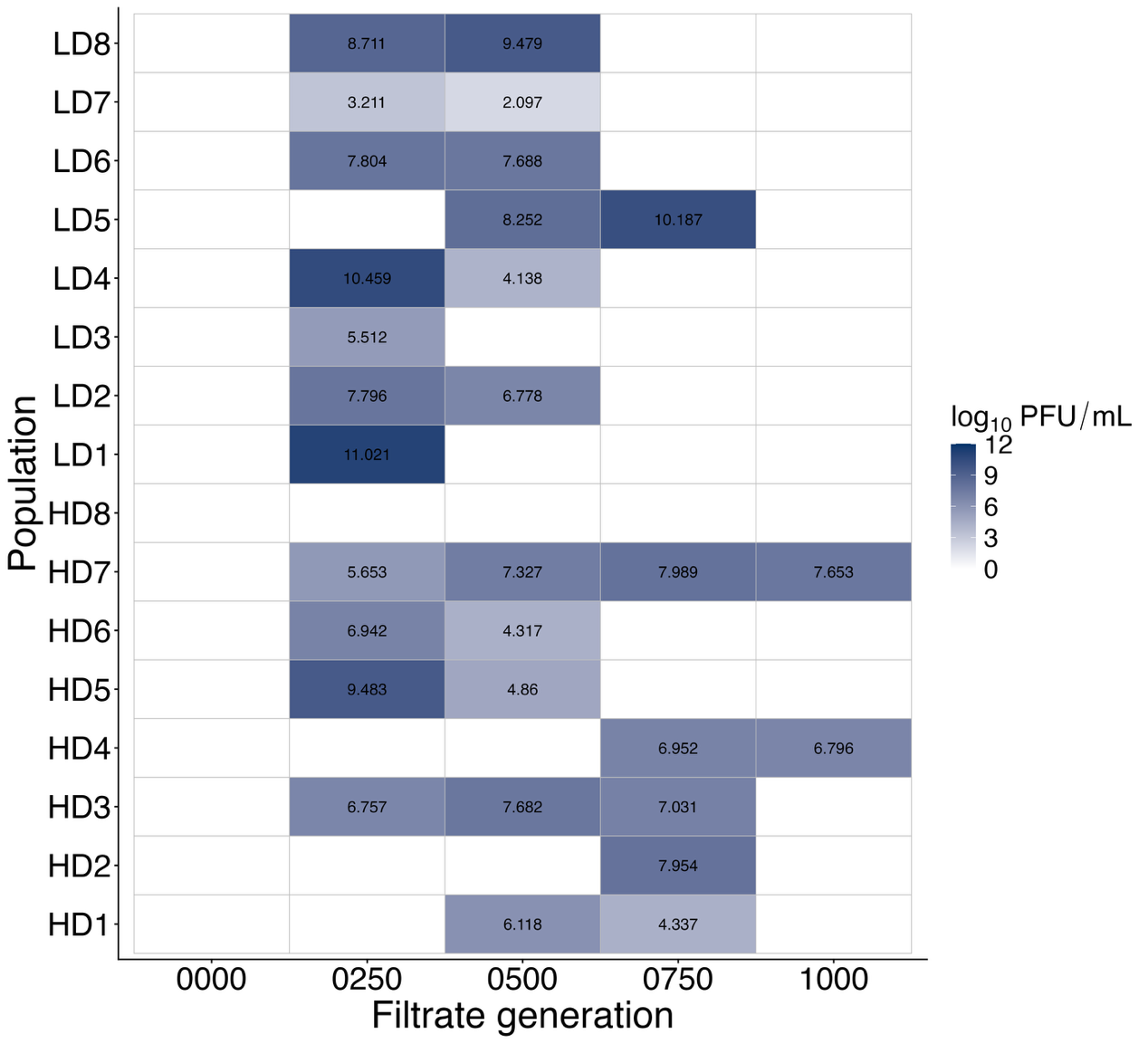

**Supplementary Figure 6: Filtrate phage density over generations.** Each value represents the mean phage density detected in filtrates across two independent cultures grown on separate days, each plated in duplicate (n = 4). Empty boxes represent filtrates in which phage were never detected.

Alt text: Tile plot summarizing the titers of hyperactive phage detected in filtrate across populations and evolutionary timepoints.

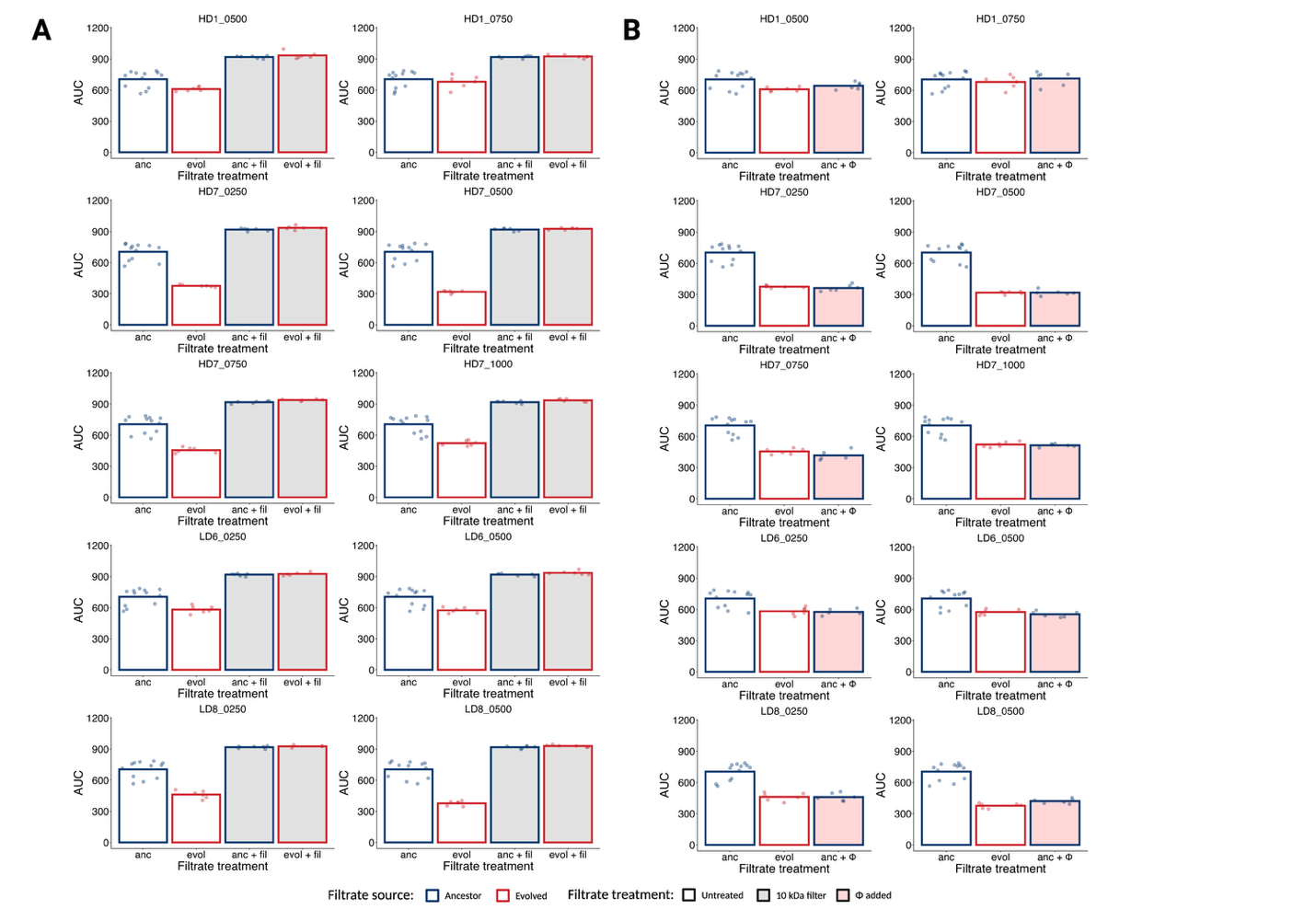

**Supplementary Figure 7: Phage presence explains inhibitory filtrate across populations**. **A:** Phage removal results across 10 filtrates sampled from four populations. AUC of MPAO1 when grown in ancestral (anc) or evolved (evol) filtrate that was either untreated or passed through a 10 kDa centrifugal filter to remove phage (+ fil). **B:** Phage addition results across 10 filtrates sampled from four populations. AUC of MPAO1 in ancestral (anc) or evolved (evol) filtrate, as well as ancestral filtrate with a mix of phage isolated from evolved filtrate spiked in (+ Φ). Created in BioRender. Turner, P. (2026) https://BioRender.com/xpzwtja.

Alt text: Multipaneled figure providing the AUC of MPAO1 grown in the presence of evolved and ancestral filtrate with or without centrifugal filtration as well as ancestral filtrate with a mix of hyperactive phage isolates spiked-in across populations.

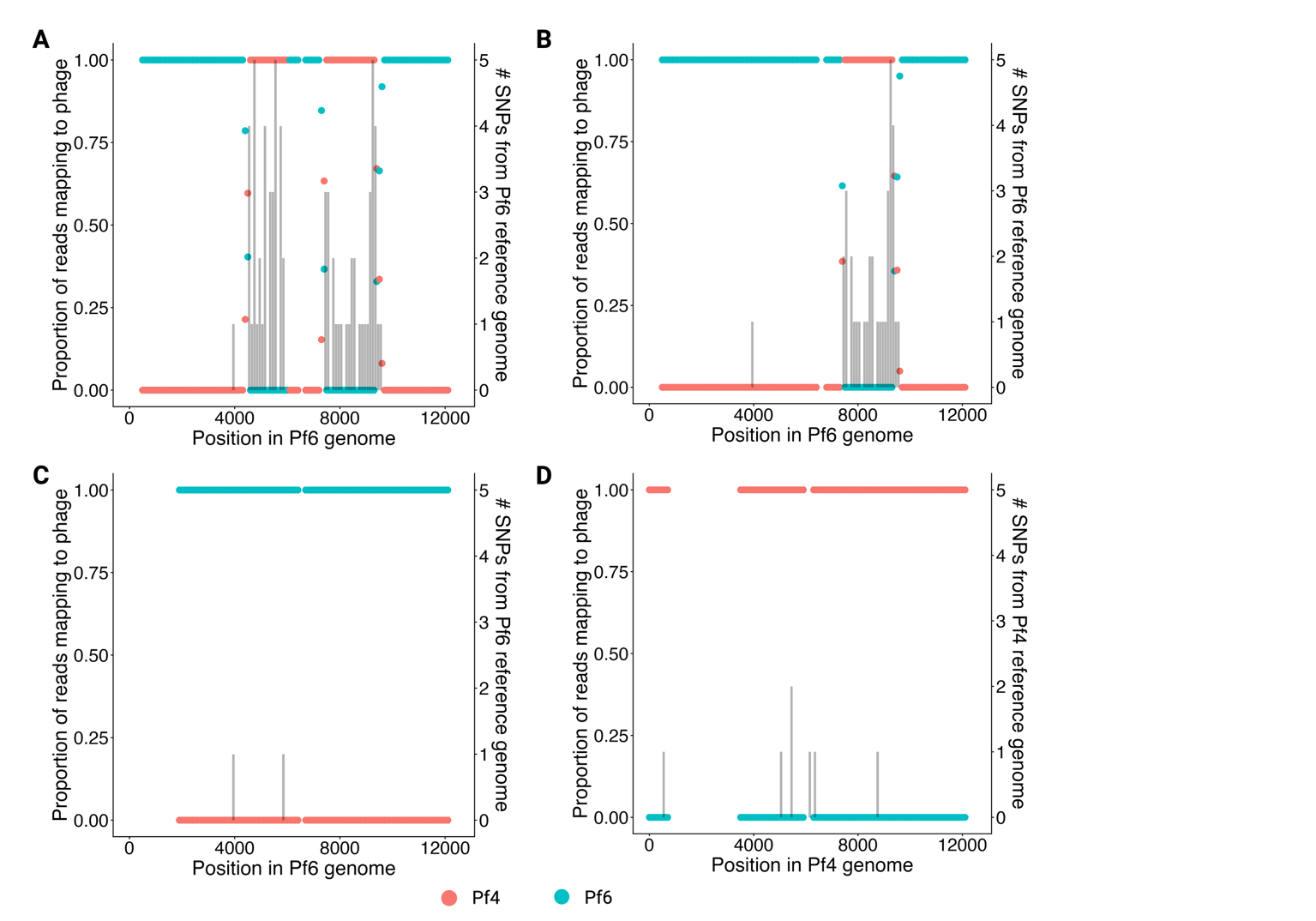

**Supplementary Figure 8: Phage isolates LD6_0250_Φ1 and LD6_0250_ Φ2 appear to be recombinants of Pf6 and Pf4**. Proportion of reads mapping to Pf4 vs. Pf6 reference genome for **A** LD6_0250_Φ1, **B** LD6_0250_Φ2, **C** LD6_0250_ Φ3, and **D** HD7_1000_ Φ1. Gaps in read mapping data indicate loci where reads did not map to either phage. Grey histograms indicate density of SNPs for each phage isolate relative to their respective reference genomes (Pf6 for **A**, **B**, **C**; Pf4 for **D**). Created in BioRender. Turner, P. (2026) https://BioRender.com/a9cfq1t.

Alt text: Bar plot visualizing the frequency of SNPs detected in the genome of four hyperactive filamentous phage isolates across regions of apparent Pf4-Pf6 recombination.

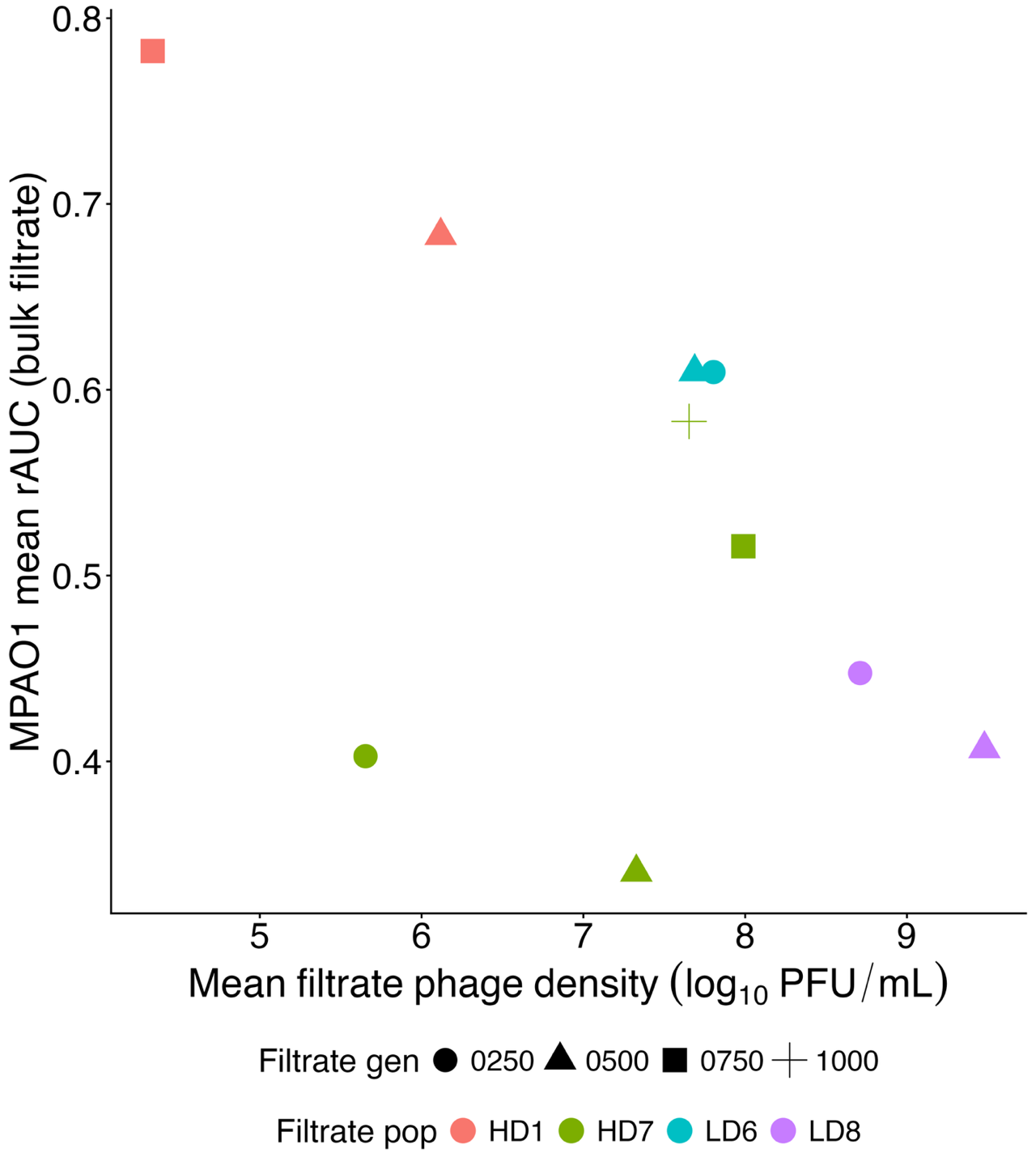

**Supplementary Figure 9: Variation in rAUC across filtrates not explained by phage density.** Relationship between filtrate phage density and the extent of inhibition caused by that filtrate in growth curves (rAUC of MPAO1). Each point represents the mean phage titer of filtrates across two replicates.

Alt text: Scatter plot of the relationship between MPAO1 rAUC in bulk filtrates to the mean filtrate phage density in the same filtrates.

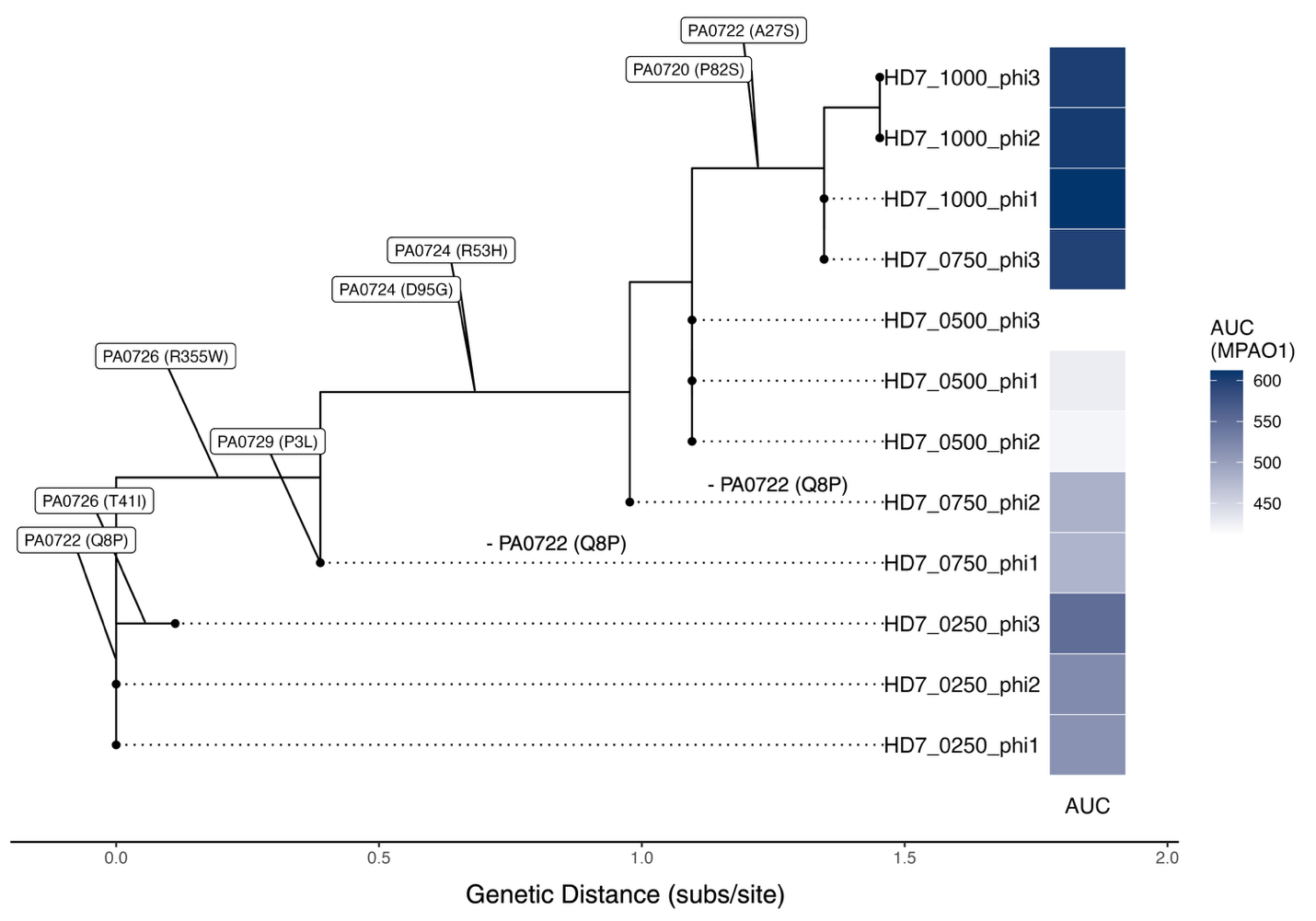

**Supplementary Figure 10: Mutations in PA0722 associated with transitions in virulence in hyperactive Pf4 isolates from HD7 filtrate.** Maximum likelihood phylogeny of hyperactive Pf4 strains isolated from HD7 filtrate with non-synonymous SNPs mapped onto branches. Performance of MPAO1 in growth curves in the presence of each phage isolate (multiplicity of infection ~ 0.01) plotted along tree tips. Created in BioRender. Turner, P. (2026) https://BioRender.com/7yp1h2u

Alt text: Phylogenetic tree of hyperactive Pf4 isolates collected from HD7 filtrates with each detected SNP annotated. AUC of MPAO1 in the presence of each isolate is plotted next to each tree tip.
